# *Let-7* restrains an oncogenic circuit in AT2 cells to prevent fibrogenic cell intermediates in pulmonary fibrosis

**DOI:** 10.1101/2024.05.22.595205

**Authors:** Matthew J. Seasock, Md Shafiquzzaman, Maria E. Ruiz-Echartea, Rupa S. Kanchi, Brandon T. Tran, Lukas M. Simon, Matthew D. Meyer, Phillip A. Erice, Shivani L. Lotlikar, Stephanie C. Wenlock, Scott A. Ochsner, Anton Enright, Alex F. Carisey, Freddy Romero, Ivan O. Rosas, Katherine Y. King, Neil J. McKenna, Cristian Coarfa, Antony Rodriguez

## Abstract

Analysis of lung alveolar type 2 (AT2) progenitor stem cells has highlighted fundamental mechanisms that direct their differentiation into alveolar type 1 cells (AT1s) in lung repair and disease. However, microRNA (miRNA) mediated post-transcriptional mechanisms which govern this nexus remain understudied. We show here that the *let-7* miRNA family serves a homeostatic role in governance of AT2 quiescence, specifically by preventing the uncontrolled accumulation of AT2 transitional cells and by promoting AT1 differentiation. Using mice and organoid models, we demonstrate genetic ablation of *let-7a1/let-7f1/let-7d* cluster (*let-7afd*) in AT2 cells prevents AT1 differentiation and results in accumulation of AT2 transitional cells in progressive pulmonary fibrosis. Integration of AGO2-eCLIP with RNA-sequencing from AT2 cells uncovered the induction of direct targets of *let-7* in an oncogene feed-forward regulatory network including BACH1/EZH2/MYC which drives an aberrant fibrotic cascade. Additional analyses using CUT&RUN-sequencing revealed an epigenetic role of *let-7* in induction of chromatin histone acetylation and methylation and maladaptive AT2 cell reprogramming. This study identifies *let-7* as a key gatekeeper of post-transcriptional and epigenetic chromatin signals to prevent AT2-driven pulmonary fibrosis.

## INTRODUCTION

Lung respiratory diseases rank as major global health challenges and are among the leading causes of death worldwide. Lung regeneration after injury is facilitated by epithelial progenitor stem cells (PSC)s found in airway and alveolar regions^1–3^. These cells are increasingly recognized as crucial factors in the onset of human lung diseases. Interstitial lung diseases (ILD)s are respiratory disorders characterized by excessive accumulation of extracellular matrix (ECM) and fibrotic tissue in alveoli leading to respiratory failure^4^. This condition is linked to abnormalities in facultative epithelial PSCs, especially alveolar type 2 (AT2) cells^5,6^. AT2 cells are crucial for lung alveolar tissue repair, serving dual roles: as PSCs that can differentiate into alveolar type 1 (AT1) cells and as homeostatic cells (hAT2) essential for surfactant production^7–9^. Recent studies in some lung injury models found that hAT2 cells become primed (pAT2), self-renew and give rise to heterogeneous population of KRT8^+^ transitional cells (hereafter referred to as ADI, alveolar differentiation intermediates) which can become AT1s^10–13^. Studies have shown that ADIs are characterized by the activation of P53, TGFβ, EMT signaling pathways and their pathological accumulation promotes lung injury and fibrosis^10–13^. A study identified a murine ADI cluster 7 subset (ADI-7) associated with stalled AT1 differentiation^13^. In the lungs of patients with Idiopathic Pulmonary Fibrosis (IPF), the most common endotype of ILDs, fibrogenic cells resembling murine ADIs -termed “aberrant basaloid” (AB) cells-persist and appear impaired in AT1-transdifferentiation^11–14^.

Studies in human IPF tissues and animal models have shown reduced expression of *let-7* microRNA (miRNA) family members^15,16^, as well as TGFβ/SMAD3 mediated repression of *let-7d* in epithelial cells ^15^. Systemic inhibition of *let-7* function with an antagomiR was found to trigger thickening of alveolar septa and remodeling^15^. *Let-7* is recognized as a tumor suppressor which prevents the proliferation and metastasis of cancer stem cells via unrestrained induction of multiway hub oncogenes such as EZH2, MYC, and KRAS^17–20^. Also, *let-7* has been shown to act as an orchestrator of key pathways and processes involved in pulmonary fibrosis, including PI3K/AKT/MTOR and epithelial mesenchymal transition (EMT)^20,21^.

This study delves into the molecular and cellular roles of the *let-7* family in AT2s, underscoring its vital role in lung tissue homeostasis, protection against injury, and fibrosis. We revealed the crucial role of *let-7* as a molecular brake to uncontrolled expansion, activation of AT2 PSCs and formation of ADIs. Loss of *let-7* activity in AT2s also hampers cell fate where ADIs are prevented from transitioning into AT1s. Mechanistic studies highlight *let-7* as a gatekeeper of physiologic expression levels of a profibrotic oncogene gene regulatory network (OGRN) comprised of BACH1 ^22^, EZH2 ^23^, and MYC ^24^ in preservation of AT2 cell plasticity and lung homeostasis. Overall, our studies highlight a new dimension for *let-7* as a potent regulator in alveolar stem cell fate and epigenetic perturbations in pulmonary fibrosis.

## RESULTS

### Time-dependent downregulation of *let-*7 family expression during peak formation of ADIs after bleomycin-induced lung injury

We examined *let-7* expression dynamics after bleomycin-induced lung injury in mice (GSE195773)^25^. Our analysis of this published small RNA-seq dataset revealed transient downregulation of *let-7* family members at 7- and 14-days relative to controls followed by return to baseline by 21-days post-bleomycin (Supplementary Fig. 1a). Total *let-7* activity decreased approximately 25% after 7-days post-bleomycin in lungs of mice vs controls (Supplementary Fig. 1b). Approximately 75% of total activity was comprised from members: *let-7a*, *let-7b*, *let-7c*, *let-7d*, and *let-7f*. These *let-7* members are transcribed, in part, from conserved *let-7b/let-7c2* (*let-7bc2*) and *let-7a1/let-7f1/let-7d* (*let-7afd*) gene clusters^26^ and contribute upwards of 20% and 55% respectively of total *let-7* activity in naive lung (Supplementary Fig. 1b,c). The temporal downregulation of *let-7* expression coincides with the transient appearance and peak formation of injury associated KRT8^+^ ADIs following bleomycin injury^11,12^. Based on these observations, we hypothesized that *let-7* family governs ADI cell formation in alveolar regeneration.

### Depletion of *let-7afd* in AT2 cells causes spontaneous lung injury acutely

To understand the mechanism of *let-7* function in AT2 cells, we utilized our *let-7bc2* and *let-7afd* conditional knockout (KO) mice^26^. We employed *let-7bc2^f/f^*;*Sftpc-Cre^ERT^*^2/+^, *let-7afd^f/f^;Sftpc-Cre^ERT^*^2^*^/+^* mice with and without the *R26R-LSL-tdTomato* reporter (*Sftpc-tdT*) for phenotypic analysis – hereafter referred to as *let-7bc2^AT^*^2^ and *let-7afd^AT^*^2^ mice (Supplementary Fig. 1c). Quantitative RT-PCR (qPCR) from flow sorted *Sftpc-tdT^+^* AT2 cells confirmed >90% excision of *let-7bc2* or *let-7afd* clusters after intraperitoneal tamoxifen administration (iTAM) (Supplementary Fig. 1d,e).

To assess the consequence of deletion of *let-7bc*2 or *let-7afd* clusters, lung histology and physiologic phenotypic analysis were carried out following 6-days of iTAM. Genetic loss of *let-7afd* in AT2 cells led to decreased arterial oxygen saturation (SpO2) following iTAM (Fig. 1a). Approximately 28% of *let-7afd^AT^*^2^ mice developed a pulmonary hemorrhage phenotype discerned by gross lung dissections and hematoxylin and eosin (H&E) stained sections (Fig. 1b,c). The histologic analysis of *let-7afd^AT^*^2^ mice revealed diffuse bleeding into the alveolar spaces and immune cell infiltration relative to controls (Fig 1c). The *let-7bc2^AT^*^2^ mice exhibited a milder temporal drop in SpO2 relative to *let-7afd^AT^*^2^ mice but no hemorrhage phenotype (Fig. 1a, Supplementary Fig. 1f).

**Fig. 1.**
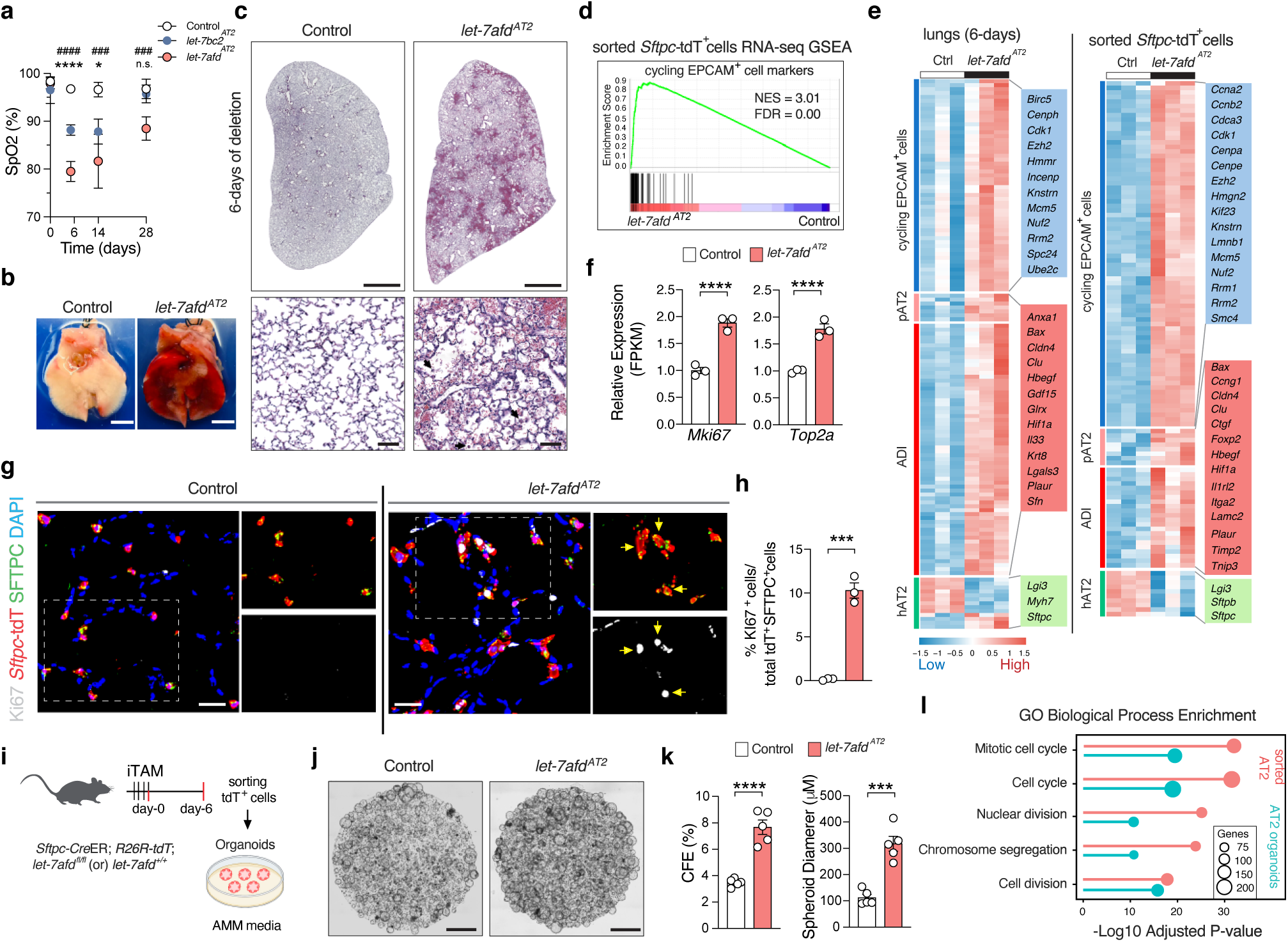
Deletion of *let-7afd* in AT2 cells promotes spontaneous lung injury and AT2 transitional cells acutely. **a** Quantitative measurements of Sp02 from control, *let-7bc2^AT^*^2^, *let-7afd^AT2^* mice (n = 5 per group) following iTAM. Data are mean±s.d. \**let-7bc2^AT2^* vs control, ^#^*let-7afd^AT2^* vs control; **, p < 0.01, ^###^, p < 0.001, **** and ^####^, p < 0.0001 by one way ANOVA with Tukey’s correction. Not significant (NS). **b**-**c** Representative lung dissections (**b**) and H&E-stained lung sections (**c**) from indicated mice 6-days after iTAM. Scale bars: 5 mm (**b**); 2 mm upper panel; 50 μm lower panel (**c**). Arrowheads indicate leukocytes. **d** GSEA plot shows induction of cycling epithelial genes in *let-7afd*^-/-^ *Sftpc-*tdT^+^ sorted AT2 cells (n = 3 samples per group). **e** Heatmap shows differentially expressed AT2 transition markers obtained by RNA-seq from whole lung or sorted *Sftpc-*tdT^+^ AT2 cells upon deletion of *let-7afd* vs controls after 6 days of iTAM. Adjusted p value < 0.05 vs control. **f** Transcript expression by RNA-seq in *Sftpc*-tdT^+^ AT2 cells after 6 days of iTAM (n = 3 samples per group). Data are mean±s.e.m. ****adjusted p < 0.0001. **g,h** Representative immunostaining (**g**) and quantification (**h**) of KI67^+^*SFTPC*^+^*Sftpc*-tdT^+^ cells in lungs of *let-7afd^AT2^* mice vs control mice 14-days after iTAM. KI67 (gray), SFTPC (green), *Sftpc-*tdT (red), DAPI (blue). (n = 3 mice per group). Data are mean±s.e.m. ***p < 0.001 by unpaired Student’s t test. Arrows indicate KI67^+^*SFTPC*^+^*Sftpc-*tdT^+^ cells (scale bar 25 μm). **i** Schematic representation to establish AT2 organoid cultures in mice. **j** Brightfield images show *let-7afd^-/-^* vs control AT2 organoids cultured in AMM. Scale bars 1 mm. **k** Quantification of CFE and spheroid diameters of organoids in panel (**j**). Data are mean±s.e.m (n = 5 mice per group). ***p < 0.001, ****p < 0.0001 by unpaired Student’s t test. Each dot represents one mouse. **l** GO BP enrichment analysis for overrepresented terms upon deletion of *let-7afd* in *Sftpc*-tdT^+^ AT2 cells (n= 3 samples per group) or organoid cultures (n = 2 mice per group) by RNA-seq. Scale shows adjusted p values. **a**-**c, g**-**h**, **j**, **k** Are representative of three independent experiments.

### Deletion of *let-7afd* in AT2 cells stimulates progenitor stem cell proliferation and formation of ADI cells acutely in the lung

We performed bulk RNA sequencing (RNA-seq) of whole-lung or flow cytometric sorting of *Sftpc-tdT*^+^ AT2s respectively between *let-7afd^AT2^* and controls following 6-days of iTAM. Gene set enrichment analysis (GSEA) showed significant induction of cell cycle related genes (*e.g*., *Mki67*), pAT2 (*e.g*., *Hif1a*) and ADI/ADI-7 (*e.g., Cldn4, Ctgf)* identity genes upon deletion of *let-7afd* in lungs and purified AT2s (Fig. 1d,e,f, Supplementary Fig. 2a,b,c). Lungs and purified AT2 cells from *let-7afd^AT^*^2^ mice also exhibit downregulation of hAT2 markers (Supplementary Fig 2a,b,c). Deletion of *let-7afd* led to downregulation of AT1 identity genes in sorted AT2s (Supplementary Fig. 2b). We additionally confirmed increased cycling AT2 lineage cells in lungs of *let-7afd^AT^*^2^ mice following 14-days of iTAM by immunofluorescence (IF) detection of nuclear KI67 in SFTPC^+^ *Sftpc-tdT^+^*cells (Fig. 1g,h).

To determine if transcriptomic changes arise, in part, due to intrinsic phenotypic changes in AT2 cells, we carried out colony forming efficiency (CFE) measurements in combination with bulk RNA-seq on *let-7afd^AT2^* and control AT2 cell-derived organoids grown in AT2 maintenance medium (AMM)^27^ (Fig. 1i). CFE and spheroid diameters were significantly increased in *let-7afd^AT2^* cells (Fig. 1j, k, Supplementary Fig. 2d). GSEA and gene ontology (GO) biological process enrichment analysis respectively revealed a striking parallel between *let-7afd^AT2^*organoids and *Sftpc-tdT* RNA-seq datasets in the induction cell cycle genes (Fig. 1l, Supplementary Fig. 2e). The *let-7afd^AT2^* organoids exhibited upregulation of pAT2 and ADI markers (Supplementary Fig. 2e). However, in contrast with sorted *let-7afd^-/-^* AT2 cells where hAT2 and AT1 markers appeared repressed, cultured *let-7afd^-/-^* AT2 organoids maintain similar marker expression as compared to controls (Supplementary Fig. 2e,f). The *let-7afd^-/-^* organoids did not exhibit significant induction of basal cell identity markers suggesting they are not redirected towards airway cell fate (Supplementary Fig. 2f). Collectively, this data indicates that acute loss of *let-7afd* expression triggers the uncontrolled proliferative expansion of AT2 PSCs and the formation of ADI cells in injured lungs acutely.

### Integrated biochemical and transcriptomics analysis in the generation of a *let-7* mRNA targetome in AT2 cells

To validate and identify functional targets of *let-7*, we performed enhanced Argonaute 2 UV-cross-linking immunoprecipitation-sequencing with enrichment of *let-7* targets (AGO2-eCLIP+let-7)^28^ in WT mice following 6-days of bleomycin injury (Fig. 2a). The libraries yielded 13,027 chimeric *let-7*:mRNA called peaks in the 3’untranslated region (3’UTR) of genes (Fig. 2b, Supplementary Data 1). Motif analysis further showed highly significant and specific enrichment of *let-7* “seed” region binding motifs within chimeric *let-7* peaks (Fig. 2c). Collapsing of chimeric *let-7* peaks into individual transcription units yielded 2009 target genes in acute bleomycin-injured lungs (Fig. 2d).

**Fig. 2.**
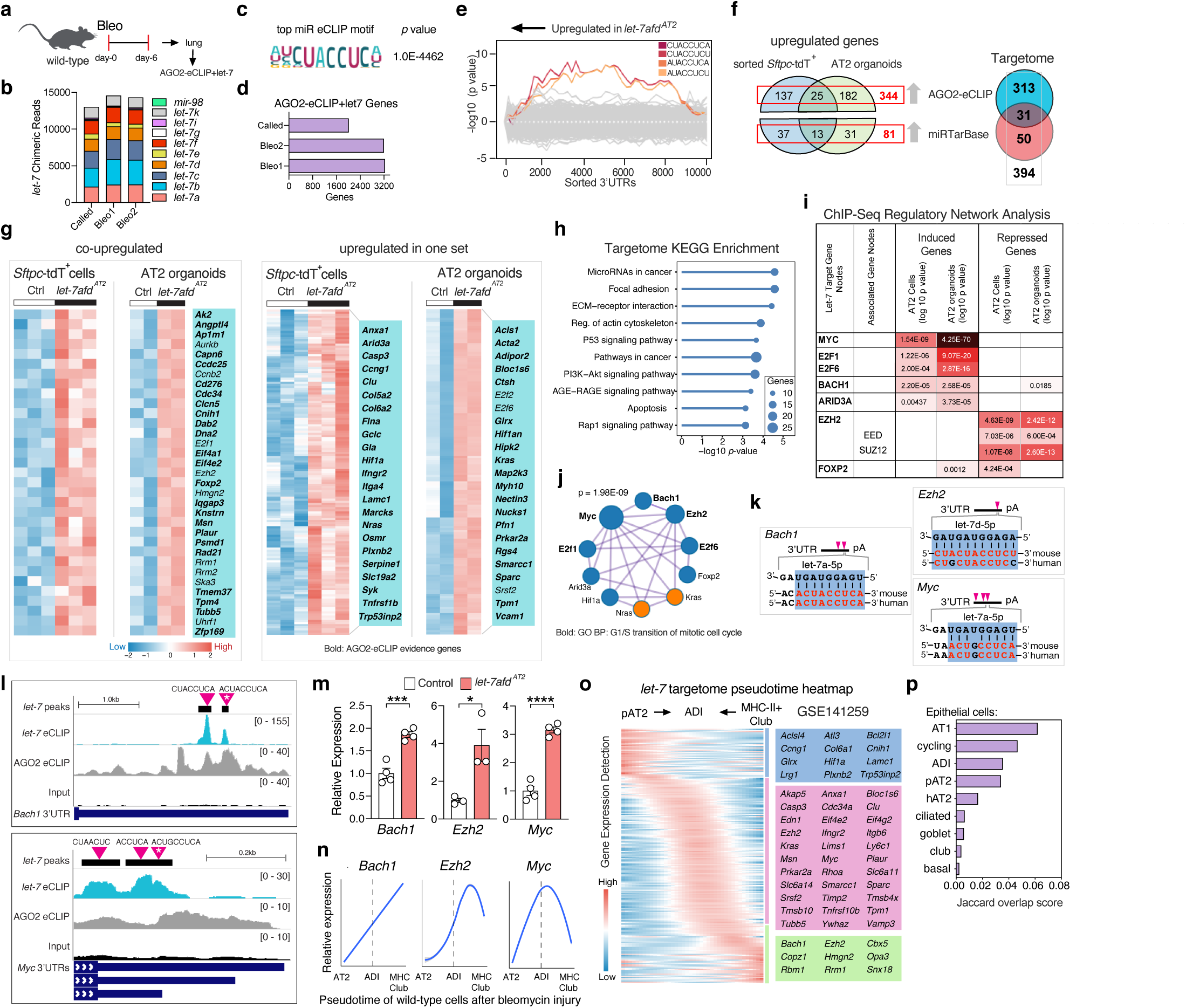
Chimeric AGO2 eCLIP and transcriptomics identify *let-7* targetome in AT2 cells. **a** Schematic of experimental design for AGO2-eCLIP+let-7 analysis from lungs of mice after bleomycin treatment. **b** Fractional representation of *let-7*:mRNA chimeric reads of *let-7* members. **c** The *let-7* family binding motif is highly enriched. **d** Total number of genes with *let-7*:mRNA chimeric peaks. **e** Sylamer analysis shows overrepresentation of *let-7* binding motifs in *let-7afd^-/-^ Sftpc*-tdT^+^ cells. **f** Venn Diagram shows gene totals to obtain the *let-7* AT2 cell targetome. **g** Heatmaps shows relative expression of significantly upregulated and validated *let-7* target genes upon loss of *let-7afd* in *Sftpc*-tdT^+^ AT2 cells (n= 3 samples per group) or AT2 organoids (n = 2 mice per group) bulk RNA-seq datasets. Adjusted p value < 0.05 vs controls. **h** KEGG pathway enrichment analysis for the *let-7* targetome. **i** ChIP-Seq cistromic analysis indicate significant enrichment of nodes corresponding to the transcription factor (TF) targets of *let-7* in sorted *let-7afd^-/-^ Sftpc*-tdT^+^ cells and organoids. **j** Functional pathway and protein interactome network shows significant connectivity of *let-7* hub targets. TFs (blue), enzymes (orange). Cytoscape/Enrichment Map P value. **k** Schematic alignments of *let-7* “seed” region with target mRNA sequences in mice and humans. **l** UCSC genome browser tracks from AGO2-eCLIP+let-7 indicates binding of *let-7* family to the 3’UTR of genes. Purple triangles indicate motifs of *let-7*. White asterisks indicate sequence alignment in panel (**k**). **m** qPCR analysis of indicated *let-7* targets from *let-7afd^-/-^* vs control sorted *Sftpc*-tdT^+^ cells 14-days after iTAM. (n = 3-4 per group). Data are mean±s.e.m. ****p < 0.0001, ***p < 0.001, *p < 0.05 by unpaired Student’s t test. **n** Line plots show smoothed relative expression levels of hub genes across the ADI pseudotime trajectories after bleomycin injury (GSE141259). Dashed line corresponds to *Krt8* peak and gray colors represent the 95% confidence interval derived from smoothing fit. **o** The expression pattern for 217 of 394 *let-7* targetome genes along the AT2 or MHC-II^+^ club cell trajectory into ADIs is based on inferred likelihood of detection. **p** Jaccard overlap scores were calculated between *let-7* targetome and epithelial cell types defined in GSE141259.

To enrich for functional targets of *let-7* in AT2 cells, we integrated the AGO2-eCLIP+let-7 targetome with the sorted *Sftpc-tdT^+^* and/or organoid transcriptome datasets. Initially, we applied Sylamer ^29^ on *let-7afd^-/-^* transcriptome data to ascertain *let-7* post-transcriptionally represses mRNA targets. Sylamer showed prominent miR-mediated mRNA destabilization since the motifs corresponding to *let-7* were enriched within the 3’UTRs of induced genes in *let-7afd^-/-^ Sftpc-tdT*^+^ cells compared to controls (Fig. 2e). Next, we made use of Venn Diagram intersection analysis to select biochemically validated, significantly upregulated, and experimentally validated *let-7* target genes in sorted *Sftpc*-tdT^+^ and organoid transcriptome datasets. The integrated approach revealed a *let-7* AT2 targetome comprised of 394 genes (Fig. 2f,g, Supplementary Data 2).

### Pathway discovery of *let-7* mRNA targetome identifies leading edge genes associated with stem cell renewal, cell growth, and cell differentiation

Functional enrichment analysis of the *let-7* targetome highlighted diverse networks of *let-7* targets associated with an array of cellular programs including stem cell renewal, growth, and differentiation (Supplementary Data 2,3). Interestingly, KEGG enrichment analysis showed the top pathways as miRNAs in cancer, focal adhesion, ECM-receptor interaction, and P53 signaling (Fig. 2h, Supplementary Data 3). MiRNAs typically govern cell programs upstream of transcription factors (TF) and multiway hub genes in gene regulatory networks (GRN)^18–20,29^. Annotation of the *let-7* targetome revealed TFs which includes gene activators (*e.g*. E2f1, *Hif1a*, *Myc*), gene repressors and context dependent inducers/repressors of gene expression (*e.g*., *Bach1*, *Ezh2, Foxp2*) (Supplementary Data 1,2,3). Major hub genes, *Kras* and *Nras* were also identified as targets of *let-7* (Supplementary Data 1,2,3). ChIP-Seq regulatory analysis on transcriptome datasets showed significant enrichment for MYC, E2F1, E2F6, and BACH1 targets in genes upregulated after depletion of *let-7afd* (Fig. 2i, Supplementary Data 4). Targets of EZH2 were overrepresented in downregulated genes upon loss of *let-7afd* (Fig. 2i). Intriguingly, the same is also true for the EZH2 partner proteins SUZ12/EED and core components of the Polycomb Repressive Complex 2 (PRC2) which mediate tri-methylation of histone H3 on lysine 27 (H3K27me3),^30,31^ implying that deletion of *let-7afd* promotes EZH2 mediated epigenetic silencing (Fig. 2i). Additionally, loss of *let-7afd* promotes the induction of KRAS pathway in AT2 cells (Supplementary Fig. 3a). Network analysis revealed strong connectivity of hub targets implicating the existence of a *let-7* dependent oncogenic GRN (OGRN) in AT2 cells (Fig. 2j). We note the unique properties of OGRN genes as sufficient drivers of uncontrolled cell growth and failed terminal differentiation in disease^30,32,33^. We annotated the location of the *let-7* chimeric peaks and binding motifs for *let-7* within 3’UTRs (Fig. 2k,l, Supplementary Fig. 3b,c, Supplementary Data 1,2). *Bach1*, *E2f1*, *Ezh2*, *Myc*, *Kras*, and *Nras* contain evolutionarily conserved sites and were previously identified as direct targets of *let-7* in non-AT2 cells^17–20,34^. *Hif1a* emerged as a direct target of *let-7*; however, the binding site is poorly conserved in humans (Supplementary Fig. 3b,c). We confirmed significant increased mRNA expression of OGRN genes by qPCR detection in sorted *let-7afd*^-/-^ *Sftpc-tdT^+^* cells following 14 days of iTAM (Fig. 2m, Supplementary Fig. 3d).

### Transcriptional convergence of alveolar and PSCs show enrichment of *let-7* targetome genes in ADI cells

We evaluated the expression dynamics of the *let-7* targetome during AT2 cell “bridging” into ADIs following bleomycin injury from a published scRNA-seq dataset^12^. At the same time, we also incorporated expression data for MHC-II^+^ club cells because they were associated with production of AT2 intermediate cells in bleomycin injury. Interestingly, *Bach1, Ezh2, Kras, Myc,* and *Nras* expression levels were increased along the differentiation trajectory from AT2 towards ADIs (Fig. 2n, Supplementary Fig. 3e). Conversion of MHC-II^+^ club cells into KRT8^+^ ADI showed a more nuanced pattern for *Ezh2* and *Bach1* indicated by an increase followed by a reduction of expression (Fig. 2n). Most *let-7* OGRN genes are upregulated in WT AT2 cells after 5-days of bleomycin treatment, suggesting they are co-regulated during lung regeneration (Supplementary Fig. 3f). Many *let-7* targets in our dataset exhibit a gradual increase and peak in expression in ADIs relative to both AT2 and MHC-II^+^ club cells (Fig. 2o). The *let-7* targetome exhibits greater enrichment in AT1s and transitional AT2s than hAT2s (Fig. 2p). Together, these results support the notion that *let-7* serves as a braking mechanism in generation of pAT2/ADIs following lung injury and alveolar regeneration upstream of a stress-inducible OGRN. These data prompted us to explore the phenotypic consequences of chronic absence of *let-7afd* in pulmonary fibrosis.

### Chronic deletion of *let-7afd* in AT2 cells induces progressive and spontaneous lung fibrosis

At 1-month of iTAM (Supplementary Fig. 4a), both male and female *let-7bc2^AT2^* and *let-7afd^AT2^* mice showed spatially heterogenous disruption of the distal lung architecture marked by enlargement of alveolar spaces and increased presence of alveolar and peribronchial leukocytes and hemosiderin-laden macrophage infiltrates (Fig. 3a,b,e, Supplementary Fig. 4b,c). Approximately 30% of *let-7afd^AT2^* mice furthermore exhibit heterogenous disorganization of alveoli including septal wall thickening and the appearance of interstitial cells (Fig. 3a,b). The *let-7afd^AT2^*mice additionally show isolated areas of alveolar septal destruction, fibroblastic foci, and collagen deposition primarily at the periphery of the lung (Fig. 3a,b,f,g). A parallel histologic evaluation of *let-7bc2^AT2^* mice at 1-month of iTAM indicates a milder phenotype without lung collagen deposition (Fig. 3a,b,f,g). When the *let-7afd^AT2^* mice were followed for 2-months of iTAM, they exhibit resolution of pulmonary fibrosis but retain enlargement of alveolar spaces vs controls (Supplementary Fig. 4d-g). Pulmonary biomechanics and spirometry measurements further showed that the *let-7afd^AT2^*mice exhibit a transient decrease in compliance and inspiratory capacity after 1-month, but fully recover by 2-months of iTAM (Supplementary Fig. 4h,i).

**Fig. 3.**
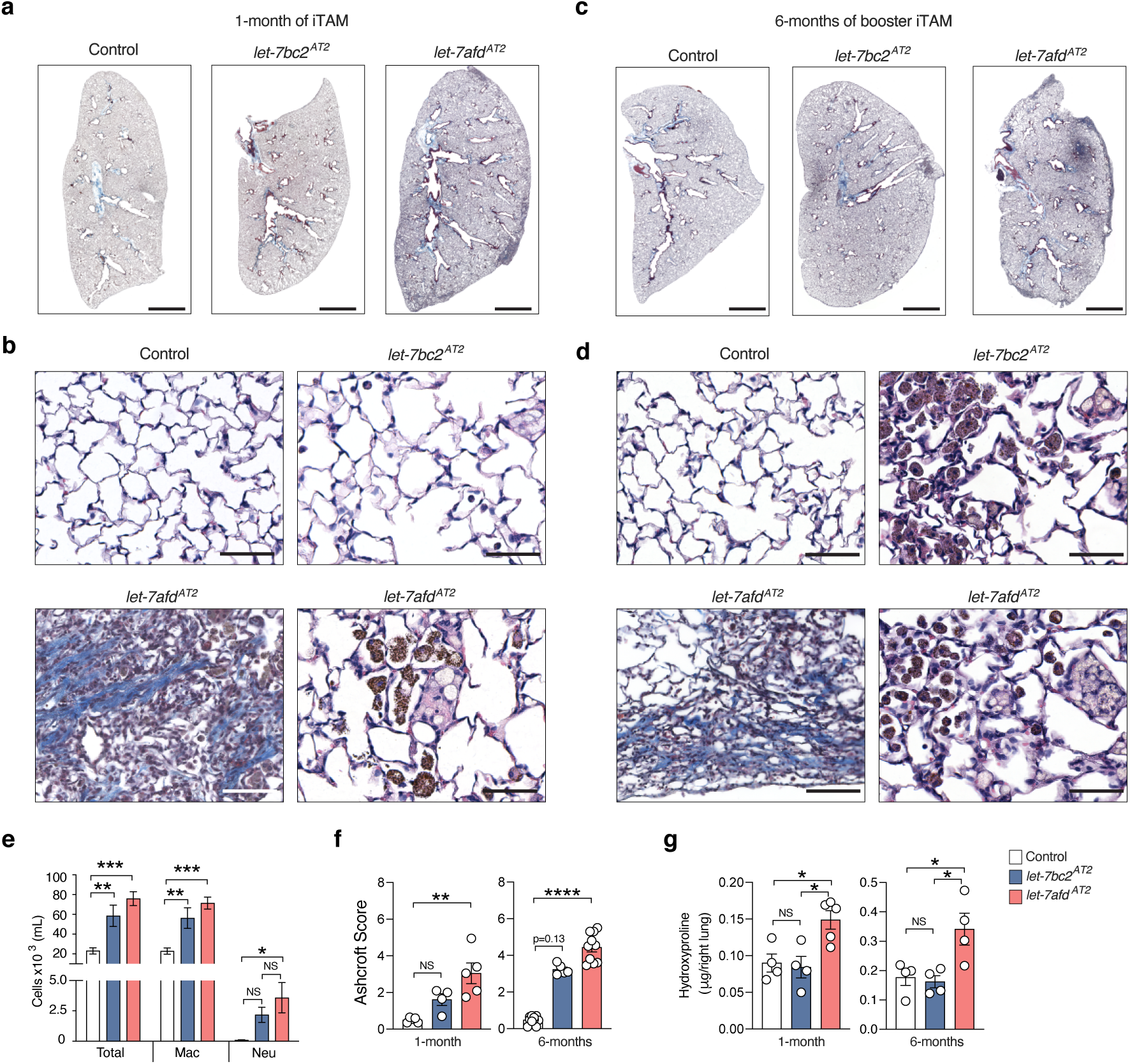
AT2-specific deletion of *let-7* clusters induce spontaneous progressive chronic inflammation and ILD. **a**-**d** Representative Masson’s trichrome-stained sections of lung lobes at 1-month iTAM (**a**,**b**) or 6-months (**c**,**d**) booster-iTAM. Scale bars: 2 mm upper panels; 50 μm lower panels (n = 16 mice per group). **e** Differential cell counts from BALF of mice is shown. Data are mean±s.e.m from individual mice (n = 7-8 mice per group). ***p < 0.001, **p < 0.01, *p < 0.05, by one way ANOVA with Tukey’s correction. Macrophages (Mac), Neutrophils (Neu). **f** Ashcroft score was used to evaluate lung injury after iTAM. (1-month, n = 4-5 per group; 6-months, n = 5 *let-7bc2^AT2^*, n = 10 controls and *let-7afd^AT2^*). Data are mean±s.e.m. ****p < 0.0001, **p < 0.01, by Dunn’s multiple comparison test. **g** Hydroxyproline levels from lungs of control, *let-7bc2^AT2^*, *let-7afd^AT2^* mice at 1-month iTAM (n = 4-5 mice per group) or 6-months of booster iTAM (4 mice per group). Data are mean±s.e.m.*p < 0.05 by one way ANOVA with Tukey’s correction. **f**,**g** Each circle represents a mouse. **a**-**g** Representative of three independent experiments. Not significant (NS).

To determine if continual Cre/loxP deletion of *let-7* clusters maintains pulmonary fibrosis over 1-month, *let-7bc2^AT2^* and *let-7afd^AT2^* mice received monthly iTAM boosters for up to 6-months (Supplementary Fig. 5a). With this regimen, the pulmonary inflammation, heterogeneous disruption of alveolar architecture and fibrotic phenotype persisted up to 6-months in >90% of *let-7afd^AT2^* mice (Fig. 3c,d,f,g). In contrast, the *let-7bc2^AT2^* mice displayed a milder ILD phenotype relative to *let-7afd^AT2^* mice but without collagen deposition (Fig. 3c,d,f,g). Pulmonary biomechanics demonstrated the *let-7afd^AT2^* mice exhibit more robust restrictive impairment in lung function vs controls and *let-7bc2^AT2^*mice (Supplementary Fig. 5b). To discern the transcriptome of the *let-7afd^AT2^* mice, we carried out bulk RNA-seq on whole lungs of mice at 6-months of booster iTAM. Differential gene expression analysis and GSEA indicated significant induction of inflammation and fibrosis pathways in lungs of *let-7afd^AT2^* mice (Supplementary Fig. 5c). Overall, these data establish that *let-7afd* deletion in AT2 cells models progressive spontaneous pulmonary fibrosis.

### Ablation of the *let-7afd* in AT2 stimulates the persistence of ADI cells

Based on this data, we hypothesized that AT2 cell loss of *let-7afd* drives the proliferative expansion of fibrogenic AT2 intermediates. We validated the presence of supernumerary AT2 lineage cells in lungs of *let-7afd^AT2^* mice compared to controls after 1-month of iTAM and 5-months of booster iTAM respectively by IF quantification of *Sftpc-*tdT*^+^* cells (Fig. 4a). Analysis of contiguous lung H&E sections for fibroblastic aggregates and IF detection of KRT8 revealed the spatial co-occurrence of ADIs in damaged areas with mesenchymal cell expansion and alveolar destruction in *let-7afd^AT2^* mice (Fig. 4b). Further quantitative IF analysis confirmed the persistence of KRT8^+^ ADIs in lung parenchyma of *let-7afd^AT2^* mice at 1- and 5-months of deletion (Fig. 4c,d). Cells expressing KRT8 exhibited distinct cell shape including elongated squamous morphology as recently described for ADIs (Fig. 4b,c)^11,12^. IF also confirmed increased *Sftpc-tdT^+^* labelled cells co-expressing KRT8 or CLDN4 in *let-7afd^AT2^* mice compared to controls (Fig. 4e,f). Reinforcing our hypothesis, at 6-months of booster iTAM lungs of *let-7afd^AT2^*mice furthermore exhibit prolonged expression of ADI and ADI-7 cell markers (Fig. 4g,h).

**Fig 4.**
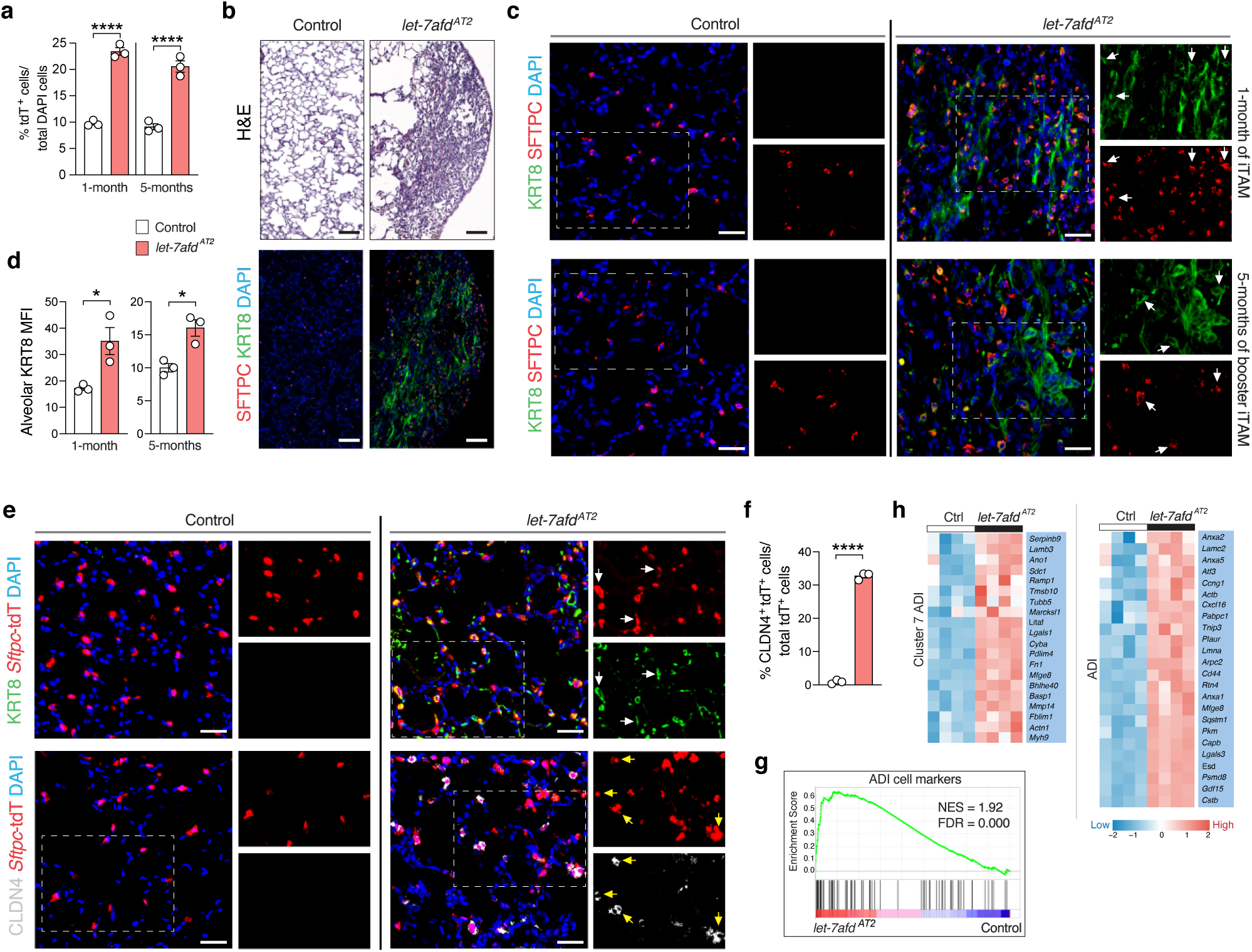
Ablation of *let-7afd* in AT2 cells stimulates the persistence of lung ADI intermediate cells. **a** Quantification of *Sftpc*-tdT^+^ cells from total DAPI cells in alveolar regions of *let-7afd^AT2^* mice compared to control mice at 1- or 5-months of booster iTAM (n = 3 mice per group). Data are mean±s.e.m. ****p < 0.0001 by unpaired Student’s t test. **b** Representative H&E-stained (upper panels) and immunostaining (lower panels) serial sections of lung after 1 month of iTAM in *let-7afd^AT2^* mice compared to control mice. IF shows KRT8 (green), SFTPC (red), DAPI (blue). Scale bars: 50 μm. **c** Representative immunostaining of KRT8 (green), SFTPC (red), DAPI (blue) at 1 month (upper panels) or 5 months of booster iTAM (lower panels) in *let-7afd^AT2^* mice vs controls. Arrowheads point to KRT8^+^SFTPC^+^ cells. Scale bars: 25μm. **d** Quantification of KRT8 mean fluorescence intensity (MFI) from alveolar regions excluding airways of mice groups in panel (**c**) (n = 3 mice per group). Data are mean±s.e.m. *p < 0.05 by unpaired Student’s t test. **e** Representative immunostaining of *Sftpc*-TdT^+^ traced KRT8^+^ or CLDN4^+^ cells in lungs of *let-7afd^AT2^* mice compared to controls 1-month post-iTAM. KRT8 (green), CLDN4 (gray), *Sftpc*-tdT (red), DAPI (blue). Arrowheads point to KRT8^+^ or CLDN4^+^ *Sftpc*-tdT^+^ cells. Scale bars: 25 μm. **f** Quantification of *Sftpc*-tdT^+^CLDN4^+^ from total *Sftpc*-TdT^+^ cells at 1-month post-iTAM (n = 3 mice per group). Data are mean±s.e.m. ****p < 0.0001, by unpaired Student’s t test. **g,h** GSEA and heat map plots derived from RNA-seq show significant induction of ADI genes in lungs of *let-7afd^AT2^* mice vs controls with 6-months of booster iTAM (n = 4 per group). Adjusted p value < 0.05 vs controls. **a**-**f** Representative of three independent experiments.

### Deficiency of *let-7afd* AT2 cell hypertrophy and stimulates the expression of profibrotic genes

To determine whether deletion of *let-7afd* promotes AT2 cell hypertrophy and/or other maladaptive ultrastructural cell changes associated with clinical ILD^5^, we made use of transmission electron microscopy (TEM) in lungs of mice after 3-months of booster iTAM. TEM stereological analysis revealed the *let-7afd^AT2^* mice exhibit enlarged or hypertrophic AT2 cells with increased area relative to controls (Fig. 5a,b). Notably, *let-7afd^-/-^* AT2 cells also displayed increased number and total cell area of lamellar bodies (LBs) with disorganized lamellae (Fig. 5a,b). The morphometric measurements of circularity furthermore show cell shape elongation of *let-7afd*^-/-^ AT2 cells compared to controls (Fig. 5a,b). Remarkably, *let-7afd^AT2^*mice also exhibit binucleated AT2 cells (11.1%, 5 of 45 cells) a unique feature not evidenced in control mice (0%, 0 of 44 cells) (Fig. 5a).

**Fig 5.**
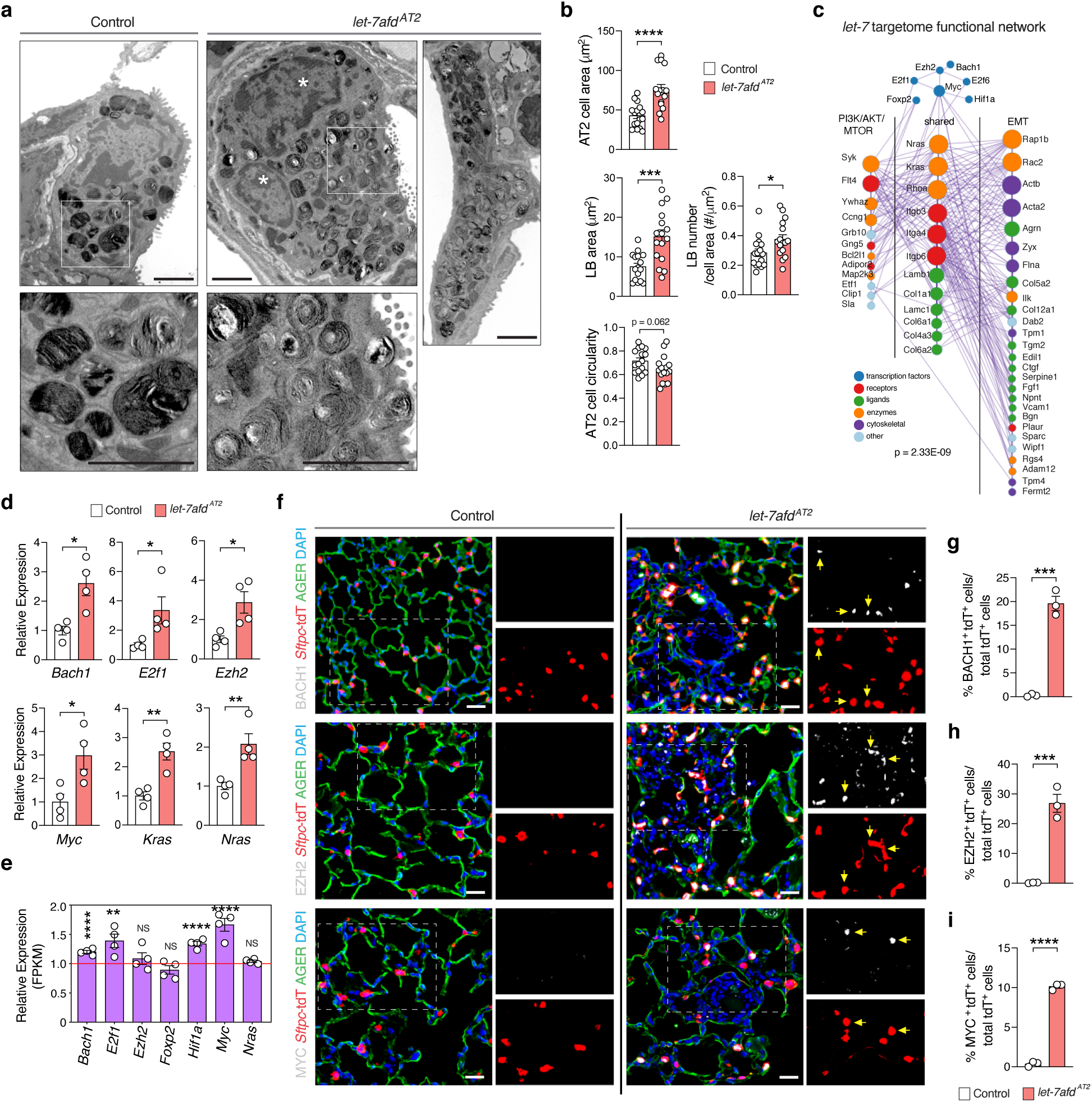
Loss of *let-7afd* stimulates AT2 cell hypertrophy and induces the expression of *let-7* target genes associated with cell hypertophy, ECM remodeling, and fibrosis. **a** Representative TEM image shows an enlarged *let-7afd*^-/-^ AT2 cell with increased abnormal lamellar bodies (LB)s and more than one nuclei (n = 3 mice per group). The white asterisks point to individual nuclei. An elongated *let-7afd*^-/-^ AT2 cell is shown on right. Scale bars: 2 μm. **b** TEM quantification of cell area and LB cell area (μm^2^), LB number per μm^2^ of cell area, and cell circularity index in each AT2 cell. Data are mean±s.e.m. Each dot represents one AT2 cell (control AT2 cells n = 16; *let-7afd*^-/-^AT2 cells = 16, from a total of 3 mice per group; ****p < 0.0001, ***p < 0.001, *p < 0.05, and p = 0.062 by unpaired Student’s t test. **c** Functional network of *let-7* target genes associated with PI3K/AKT/MTOR and/or EMT. Network p value was computed by Cytoscape/ Enrichment Map. **d** Expression levels of hub *let-7* targets were evaluated by qPCR after 2 months booster-iTAM from *let-7afd^-/-^* and control *Sftpc*-tdT^+^ cells (n = 4 samples per group; from pools of 2 mice). Data are mean±s.e.m. **p < 0.01, *p < 0.05, by unpaired Student’s t test. **e** *Let-7* target expression was determined by RNA-seq in lungs of *let-7afd^AT2^* and control mice after 6-months of booster iTAM (n = 4 samples per group). Data are mean±s.e.m plotted relative to control samples = 1.0. **adjusted p < 0.01, ****adjusted p < 0.0001. Not significant (NS). **f** Representative IF images show increased numbers of BACH1^+^ (top panels), EZH2^+^ (middle panels), and MYC^+^ *Sftpc*-tdT^+^ cells respectively (indicated by yellow arrows) in lungs of *let-7afd^AT2^* mice vs control mice lungs after 1-month of iTAM. BACH1 (gray); EZH2 (gray); MYC (gray), *Sftpc*-tdT (red); AGER (green); DAPI (blue). Scale bars: 25 μm. **g**-**i** Quantification of BACH1^+^ (**g**), EZH2^+^ (**h**), and MYC^+^ (**i**) *Sftpc*-tdT^+^ cells in total *Sftpc*-tdT^+^ cells (n = 3 mice per group). Data are mean ±s.e.m. ****p < 0.0001, ***p < 0.001 by unpaired Student’s t test. **f**-**i** Representative of three independent experiments.

We hypothesized that *let-7* with its vast OGRN targets and impact on cell size and shape might orchestrate fibrogenesis upstream of PI3K/AKT/MTOR and ECM receptor/EMT pathways. Correspondingly, analysis of our RNA-seq datasets revealed that absence of *let-7afd* activates these profibrotic pathways in AT2 cells (Supplementary Fig. 6a,b, Supplementary Data 3). Integrative pathway annotation of the *let-7* AT2 cell targetome highlights a functionally interconnected protein interactome network, consisting of over 50 gene targets of *let-7* directly or indirectly involved in PI3K/AKT/MTOR and EMT pathways (Fig. 5c). Notably, this interactome includes genes associated with fibrosis, such as TFs (*Myc*)^24^, receptors (*Itgb3*, *Plaur*), ligands (*Ctgf*, *Col5a2*), enzymes (*Nras*, *Kras*) and cytoskeletal proteins (*Tpm1*) (Fig. 5c, Supplementary Fig. 6c,d,e, Supplementary Data 1,2,3). Most OGRN genes (e.g., *Bach1*, *Ezh2*) were included in the network given their inductive roles in tissue fibrosis^22,23^ and engagement of PI3K/AKT/MTOR^33,35^ and EMT^21,36^ pathways respectively (Fig. 5c). To further explore the potential contribution of OGRN in pulmonary fibrosis, we sought to determine whether they remained significantly induced in *let-7afd^AT2^*mice with fibrosis. Consistent with our acute phenotypic analysis, transcript levels of most OGRN genes remained upregulated in sorted AT2 cells and lungs of *let-7afd^AT2^*mice with fibrosis over controls (Fig. 5d,e). Furthermore, *let-7afd^AT2^* mice with fibrosis exhibit increased numbers of BACH1^+^, EZH2^+^ or MYC^+^ *Sftpc*-tdT traced cells than controls (Fig. 5f,g,h,i, Supplementary Fig. 6f,g,h). These findings suggests that absence of *let-7afd* drives unchecked expression of OGRN at the top of the hierarchy and ECM structural proteins downstream.

### Genetic loss of *let-7afd* makes AT2 cells vulnerable to DNA damage, senescence, and apoptosis and impedes AT2 to AT1 terminal differentiation

Studies have shown that in different models of lung injury and fibrosis, AT2 cells can undergo cell senescence and/or apoptosis related to disruptions in quality control programs^5^. Furthermore, ADI cells are inherently prone to DNA damage and cell senescence during normal injury repair and in IPF^11^. Additionally, unchecked expression of BACH1, EZH2, or MYC in non-cancer cells can trigger DNA damage, senescence, and/or apoptosis ^36–41^. We examined lungs of mice with γH2AX as a marker of DNA damage in conjunction with the β-galactosidase activity assay for cell senescence. At 1-month of deletion, lung immunostaining showed significant accumulation and frequency of γH2AX^+^ SFTPC^+^ AT2s in *let-7afd^AT^*^2^ mice compared to controls (Fig. 6a,b,c). Correspondingly, *let-7afd^AT2^*mice showed robust accumulation of β-galactosidase substrate X-gal in lung parenchyma, including AT2s indicating induction of a widespread cellular senescence program (Fig. 6d). Then we assessed for apoptotic AT2 cells in lungs of mice by IF detection of active CASP-3. This analysis revealed increased active CASP-3 *Sftpc-*tdT^+^ labeled cells in *let-7afd^AT2^* mice relative to controls (Fig 6a,c). Congruent with these findings, sorted *Sftpc-* tdT cells and whole lungs of *let-7afd^AT2^*mice exhibit induction of DNA repair, senescence, senescence-associated secretory phenotype (SASP), and apoptotic genes than controls (Fig 6f,g, Supplementary Fig. 7a,b,c). Annotation of the *let-7* targetome identified *Casp3* and *Serpine1* as potentially important mediators of these cell processes (Supplementary Fig. 7d,e, Supplementary Data 1,2,3).

**Fig 6.**
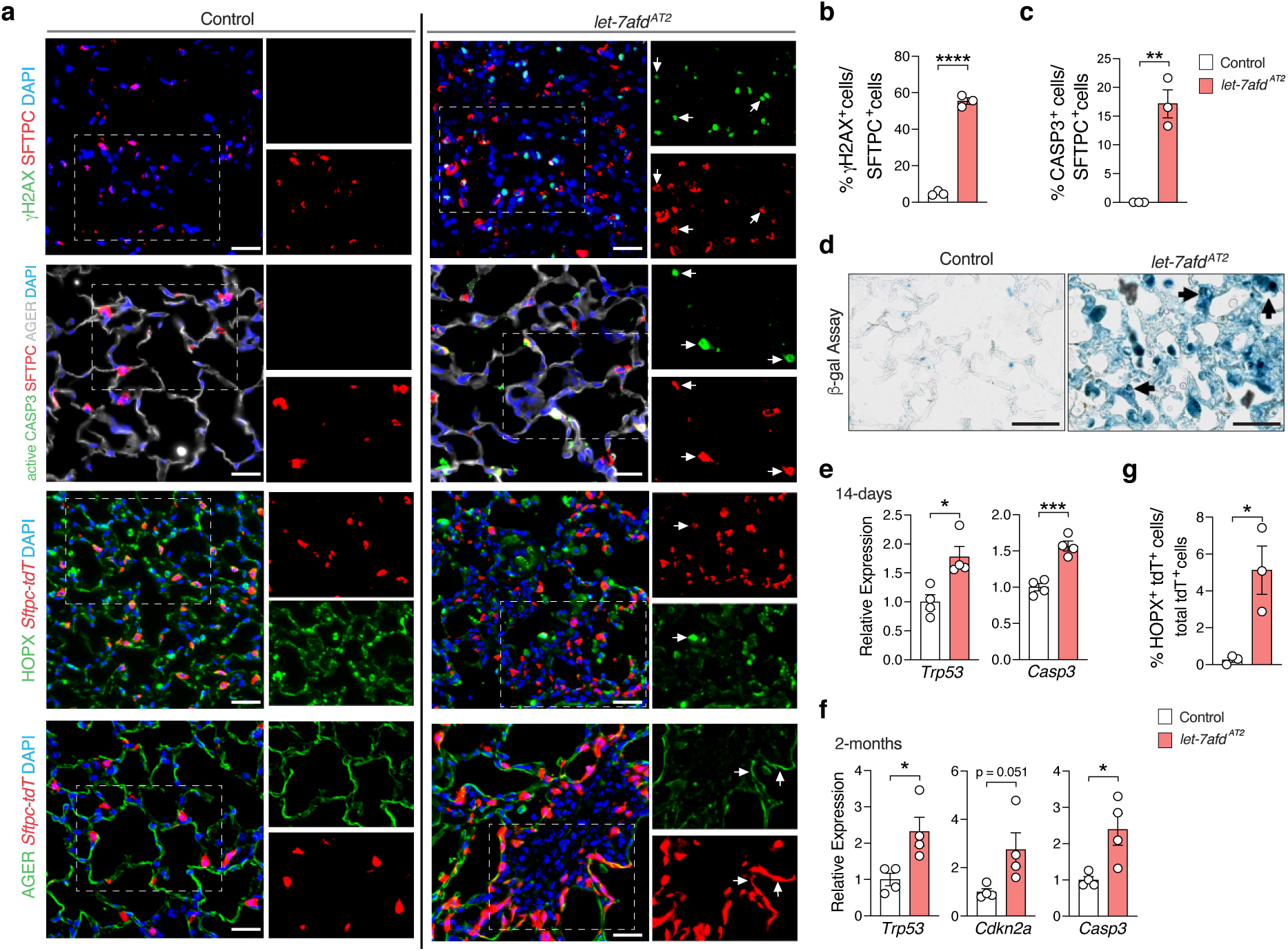
Loss of *let-7afd* is associated with AT2 cell DNA damage, senescence, and apoptosis. **a** Immunostaining for γH2AX (green), SFTPC (red), DAPI (blue) upper panels; active CASP3 (green), SFTPC (red), AGER (gray), DAPI (blue) second row panels; HOPX (green), *Sftpc*-tdT (red), DAPI (blue) third row panels; AGER (green), *Sftpc*-tdT (red), DAPI (blue) lower panels in lungs of *let-7afd^AT2^* and control mice after 1 month of iTAM. Arrows indicate γH2AX^+^SFTPC^+^ cells, active-CASP3^+^SFTPC^+^ cells, HOPX^+^*Sftpc*-tdT^+^, and AGER^+^*Sftpc*-tdT^+^cells respectively. Scale bars: 25 μm. **b**-**c** Quantification of γH2AX^+^SFTPC^+^ cells (**b**) or active-CASP3^+^ SFTPC^+^ cells (**c**) from total SFTPC^+^ cells (n = 3 mice per group). Data are mean±s.e.m. ****p < 0.0001, **p < 0.01, by unpaired Student’s t test. **d** Representative β-galactosidase staining in lungs of *let-7afd^AT2^* and control mice after 1-month of iTAM. Arrowheads point to AT2 cells in thickened alveolar septa (n = 6 mice per group). Scale bars: 50 μm. **e,f** Indicated genes were detected by qPCR from *let-7afd^-/-^* and control *Sftpc*-tdT^+^ cells after 14-days of iTAM (**e**) or 2-months (**f**) of booster iTAM (n = 4 samples per group from pools of 2 mice). Data are mean±s.e.m. *p < 0.05, ***p < 0.001, by unpaired Student’s t test. **g** Quantification of HOPX^+^*Sftpc*-tdT^+^ cells from total *Sftpc*-TdT^+^ cells in lungs of *let-7afd^AT2^* and control mice at 1-month after iTAM (n = 3 mice per group). Data are mean±s.e.m. *p < 0.05, by unpaired Student’s t test. **a-g** Representative of three independent experiments.

We posited that the persistence of ADIs in *let-7afd^AT2^* mice might relate to impaired AT1 trans-differentiation. Thus, we tested whether *let-7afd* is necessary for differentiation of AT2s into AT1s with lung IF with AT1 cell markers AGER and HOPX. The frequency of *Sftpc-*tdT^+^ cells co-expressing AT1 markers increased in *let-7afd^AT2^* mice vs controls after 1-month of deletion as an indication of AT1-transdifferentiation (Fig. 6a,g). However, consistent with the alveolar destruction and net loss of AT1 cells, AT1 identity markers were lower in lungs of *let-7afd^AT2^*mice than controls (Supplementary Fig. 7f,g). Club epithelial cell markers were also downregulated suggesting that *let-7afd* deletion also promotes damage within the airway (Supplementary Fig 7f,g).

To ascertain whether loss of *let-7afd* blunts AT2 to AT1 cell differentiation in an intrinsic manner, we compared the formation of ADIs and AT1 differentiation by IF in control and *let-7afd^-/-^* AT2 cell organoids grown in AT1 cell differentiation medium (ADM) (Fig. 7a). On day 7 of ADM culture, CLDN4^+^ and KRT8^+^ expression levels and positive cells were enhanced significantly in *let-7afd^-/-^*organoids over controls which corroborates the increased ADIs in lungs of *let-7afd^AT2^*mice (Fig. 7b,c, Supplementary Fig. 8a,b). In further support of abnormal AT1 trans-differentiation, *let-7afd^-/-^* organoids exhibited reduced AGER^+^, HOPX^+^, AQP5^+^ cells and expression in contrast with controls (Fig. 7d,e,f, Supplementary Fig. 8c,f,g). Of note, *let-7afd^-/-^*organoids also exhibit reduced expression and fewer LGALS3^+^ (a late ADI marker) cells vs controls which highlight a late block in AT1 differentiation (Fig. 7d,g, Supplementary Fig. 8d). By comparison, SFTPC^+^ cells and levels were enhanced in *let-7afd^-/-^* organoids suggesting that absence of *let-7* promotes not only the accumulation of ADIs but also reinforces retention of AT2 identity (Fig. 7d,h, Supplementary Fig. 8e). To ascertain whether *let-7afd* depletion promotes DNA damage or death in differentiation impaired *let-7afd^-/-^*organoids, we examined levels of γH2AX and active CASP3 by IF. The number of γH2AX^+^ and active CASP3^+^ cells were increased in *let-7afd^-/-^* AT2 cultures compared with controls (Supplementary Fig. 8f,h,i). Thus, *let-7afd* serves as a pivotal brake to the formation of lung injury inducing ADI cells, which are stalled in AT1 cell differentiation. Furthermore, loss of *let-7afd* renders AT2 cells prone to senescence and apoptosis in a cell intrinsic manner which could aggravate pulmonary fibrosis^42^.

**Fig 7.**
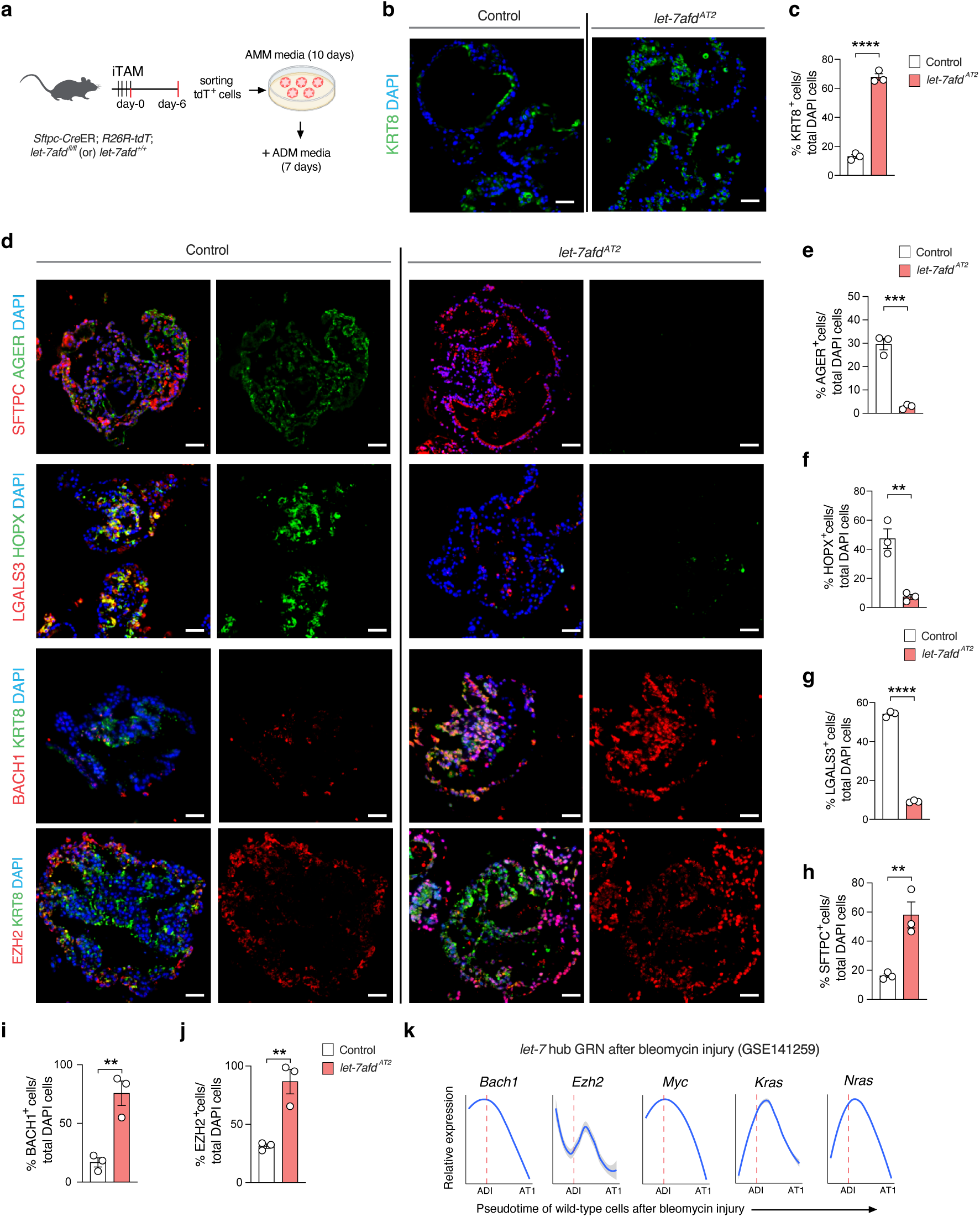
Absence of *let-7afd* impairs AT2 to AT1 differentiation. **a** Schematic representation of experimental design to examine the role of *let-7afd* in AT2 to AT1 differentiation in cultured AT2 organoids. **b**,**d** Representative immunostaining of ADI, AT1 markers and BACH1/EZH2 expression in *let-7afd^-/-^* vs control AT2 organoids grown as in panel (**a**) (n = 3 mice per group). Panel (**b**) shows KRT8 (green), DAPI (blue). Panel (**d**) shows AGER (green), SFTPC (red), DAPI (blue) top row panels; HOPX (green); LGALS3 (red); DAPI (blue) second row panels; BACH1 (red), KRT8 (green), DAPI (blue) third row panels; EZH2 (red), KRT8 (green), DAPI (blue) bottom row panels. Scale bars: 25 μm. **c**,**e**-**j** Quantification of KRT8^+^ (**c**), AGER^+^ (**e**), HOPX^+^ (**f**), LGALS^+^ (**g**), SFTPC^+^ (**h**), BACH1^+^ (**i**), or EZH2^+^ (**j**) cells respectively in total DAPI cells (n = 3 mice per group). Data are mean±s.e.m. ****p < 0.0001, ***p < 0.001, **p < 0.01, by unpaired Student’s t test. Each circle represents a mouse. Representative of three independent experiments. **k** Line plots show smoothed expression levels of selected *let-7* hub targets in the ADI to AT1 trajectory obtained from GSE141259 dataset. Grey line represents 95% confidence interval obtained from fit model. KRT8 peak represented by dashed line.

To explore potential mechanisms of impaired AT2 to AT1 trans-differentiation of *let-7afd^-/-^* AT2 cultures, we examined BACH1 and EZH2 expression by IF in ADM cultures. Interestingly, *let-7afd^-/-^*cultured AT2 organoids exhibited higher levels and frequency of BACH1^+^ and EZH2^+^ cells as compared to controls (Fig. 7d,i,j, Supplementary Fig. 8j,k). Both BACH1 and EZH2 have been associated with impaired epithelial cell differentiation upon overexpression^23,43^. Analysis of bleomycin-induced injury scRNA-seq dataset^12^ revealed a gradual decrease in dynamic expression BACH1/EZH2 and targetome as ADIs move in the direction of AT1 cell differentiation (Fig. 7k, Supplementary Fig. 8l). Taken together these analyses lend support that loss of *let-7afd* hampers AT2 cell “bridging” into AT1 cells, in part, via ectopic expression of BACH1/EZH2.

### *Let-7afd* orchestrates AT2 profibrotic dysfunction epigenetically via acetylation and methylation of histone H3 on lysine 27 (H3K27)

Pathologic overexpression of EZH2, BACH1, or MYC has been associated with epigenetic induction and repression of genes involved in cell state transitions and cell identity at the level of acetylation^44–46^ and methylation^31,47^ of H3K27 and other histones. We therefore hypothesized that deletion of *let-7* reprograms fibrogenic AT2s by promoting the deposition of H3K27ac on previously silent enhancers resulting in activation of cell growth and ADI genes, while simultaneously repressing key genes via H3K27me3 deposition.

To test this, we conducted cleavage under targets and release using nuclease (CUT&RUN)-DNA sequencing on chromatin to identify differential H3K27ac and H3K27me3 modifications in flow sorted *let-7afd^-/-^* compared to control *Sftpc-tdT^+^*cells on day 6 after iTAM treatment (Fig. 8a, Supplementary Fig. 9a, Supplementary Data 5). In *let-7afd^-/-^* AT2 cells, we observed increased H3K27ac peaks relative to control cells, which showed significant enrichment of cell cycle genes (Fig. 8b, Supplementary Data 6). Conversely, increased H3K27me3 marks in *let-7afd^-/-^*AT2 cells localized to development associated genes (Supplementary Fig. 9b, Supplementary Data 6). To identify potential TF regulatory networks, we performed motif analysis via HOMER. Several TF motifs, including BACH1, were significantly associated with H3K27ac modifications (Fig. 8c, Supplementary Data 7).

**Fig 8.**
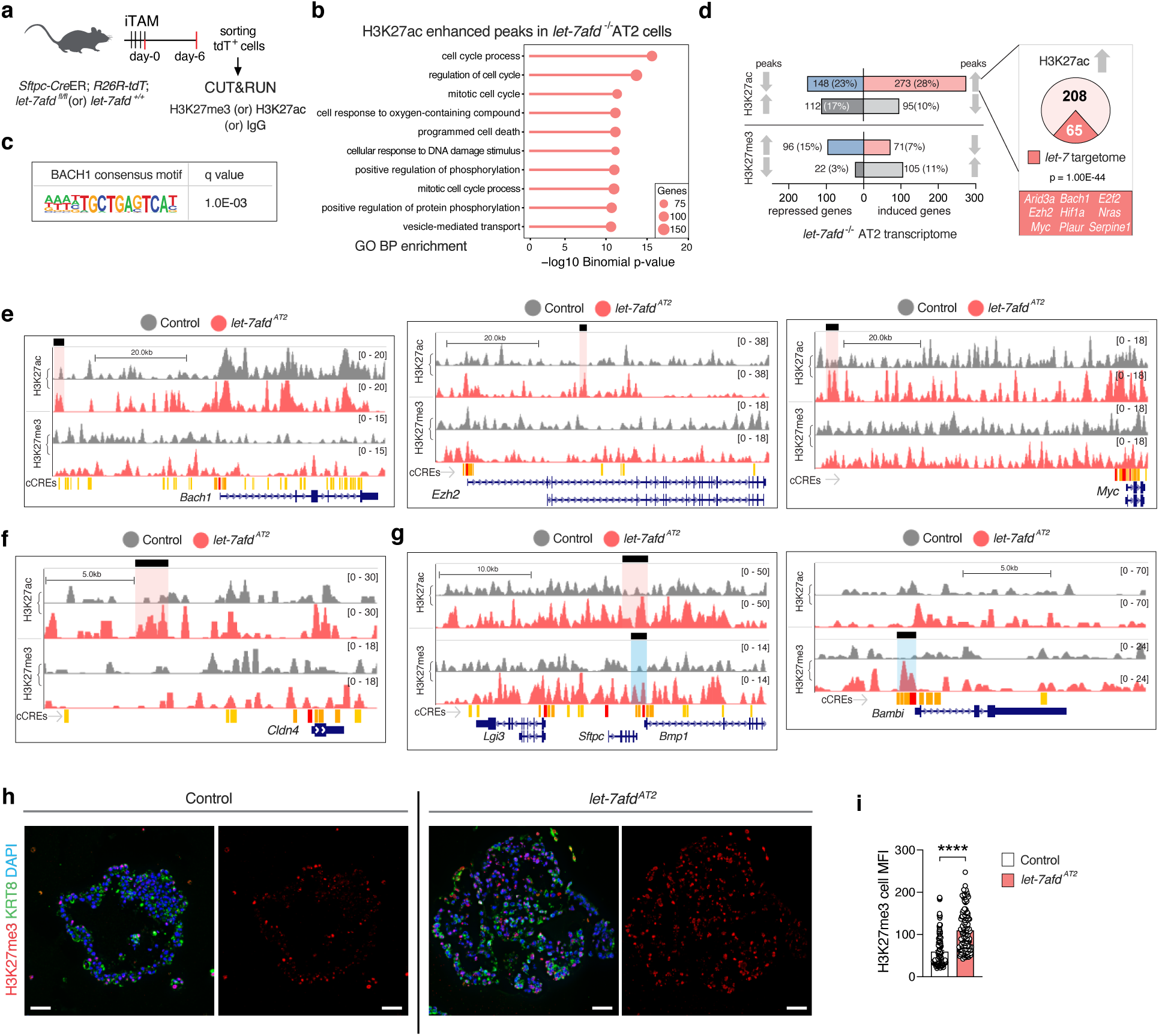
Loss of *Let-7afd* orchestrates AT2 fibrotic reprogramming, in part, epigenetically via H3K27 acetylation and methylation. **a** Experimental design scheme to carry out CUT&RUN from sorted *Sftpc*-tdT^+^ cells obtained from *let-7afd^AT2^* mice or control mice (n = 2 mice per group). **b,c** GO biological pathway (BP) (**b**) and HOMER transcription factor motif (**c**) enrichment of induced H3K27ac peaks in *let-7afd^-/-^* compared to control *Sftpc*-tdT^+^ cells. **d** Integration analysis of CUT&RUN with RNA-seq in *let-7afd^-/-^* vs control *Sftpc*-tdT^+^ cells shows correlation with transcription. Gene totals of congruent-expression change directionality (blue+red) and incongruent directionality (gray) are shown for H3K27ac and H3K27me3. Pie chart and ORA gene enrichment analysis shows that a high fraction of upregulated *let-7* targetome genes exhibit H3K27ac marks suggestive of increased enhancer activity in *let-7afd^-/-^ Sftpc*-tdT^+^ cells. **e-g** UCSC tracks show H3K27ac and H3K27me3 peaks in selected genes in *let-7afd^-/-^ Sftpc*-tdT^+^ cells compared to controls. Enriched H3K27ac peaks in *let-7afd^-/-^ Sftpc*-tdT^+^ cells are shaded in red. The enriched H3K27me peaks in *let-7afd^-/-^ Sftpc*-tdT^+^ cells vs controls are highlighted in blue. ENCODE candidate cis-regulatory elements (cCREs). **h** Representative immunostaining of H3K27me3^+^ cells in *let-7afd^-/-^* vs control AT2 organoids in ADM (n = 3 per group). H3K27me3 (red), KRT8 (green), DAPI (blue). Scale bars: 25 μm. **i** Quantification of H3K27me3^+^ nuclear MFI expression. Data are mean±s.e.m. Each dot represents one cell (n = 149 cells per genotype, 3 mice per group). Data are mean±s.e.m. ****p < 0.0001, by unpaired Student’s t test. Representative of three independent experiments.

To focus on differentially induced and repressed genes where H3K27 marks have a functional impact, we integrated the CUT&RUN analyses with our bulk transcriptomic dataset from sorted *let-7afd^-/-^* AT2 cells (Fig 8d, Supplementary 9c, Supplementary Data 8). This analysis revealed a positive correlation between enhanced H3K27ac modifications and *let-7afd*-dependent gene activation in *let-7afd^-/-^*AT2 cells (Fig. 8d). Notably, H3K27ac modifications were positively correlated with the *let-7* targetome including *Bach1*, *Ezh2*, *Hif1a*, and *Myc* (Fig. 8d,e, Supplementary Fig. 9d, Supplementary Data 5,8). Additionally, canonical ADI cell markers such as *Cldn4* displayed increased H3K27ac modifications in *let-7afd^-/-^* AT2 cells vs controls (Fig. 8f, Supplementary Fig. 9e, Supplementary Data 5,8). Interestingly, deletion of *let-7afd* also resulted in combined H3K27ac and H3K27me3 co-modifications in *Sftpc* and *Bmp1* which may indicate expression fine-tuning via bivalent chromatin (Fig. 8g). Of note, the antifibrotic and cell growth suppressor target of EZH2, *Bambi*,^48,49^ exhibits increased H3K27me3 deposition and reduced expression in *let-7afd^-/-^* AT2 cells compared to controls (Fig. 8g, Supplementary Data 5,8). To investigate whether the *let-7afd* deletion promotes H3K27me3, we examined subcellular localization by IF in cultured AT2 organoids. H3K27me3 nuclear expression levels were significantly enhanced in *let-7afd^-/-^*organoids over controls (Fig. 8h,i). In conclusion, these epigenomic data highlight the integral role of *let-7afd* chromatin remodeling, where aberrant enhancer H3K27ac activity and repressive H3k27me3 modifications lead to uncontrolled induction of cell growth and ADI marker genes.

### Human Aberrant Basaloid (AB) cells from IPF patients exhibit enrichment of the *let-7* interactome

We speculated that reduced *let-7* activity in lung epithelial cells of IPF patients might contribute to accumulation of AB cells. To investigate this hypothesis, we analyzed a published single-cell RNA sequencing dataset from human IPF lungs^14^. Few targets of *let-7* were upregulated in IPF AT2 relative to control AT2 cells (Fig. 9a,b, Supplementary Data 10). On the other hand, AB cells exhibited significant enrichment of the murine *let-7* interactome relative to AT1 and AT2 cells from IPF patients (Fig. 9a,b, Supplementary Data 10). Furthermore, upregulated targets of *let-7* in AB cells compared to control AT2s figure prominently in fibrosis pathways including ECM and EMT (Fig. 9c). Relative to AT1 and AT2 cells, AB cells furthermore exhibit enrichment of EMT genes (Fig. 9d). The *let-7* OGRN genes, *ARID3A*, *BACH1*, *EZH2*, *NRAS* were significantly upregulated in AB cells compared to control AT2 cells while FOXP2 and MYC were repressed (Fig. 9e and Supplementary Data 10). We also evaluated for consensus target signatures of EZH2 and BACH1 in IPF. This analysis found that AB cells show increased EZH2 and BACH1 target signatures compared to IPF AT2 cells (Fig. 9f,g). Our analyses suggest the *let-7* OGRN axis might contribute to the formation of AB cells in IPF; however further studies will be required for more direct evidence.

**Fig 9.**
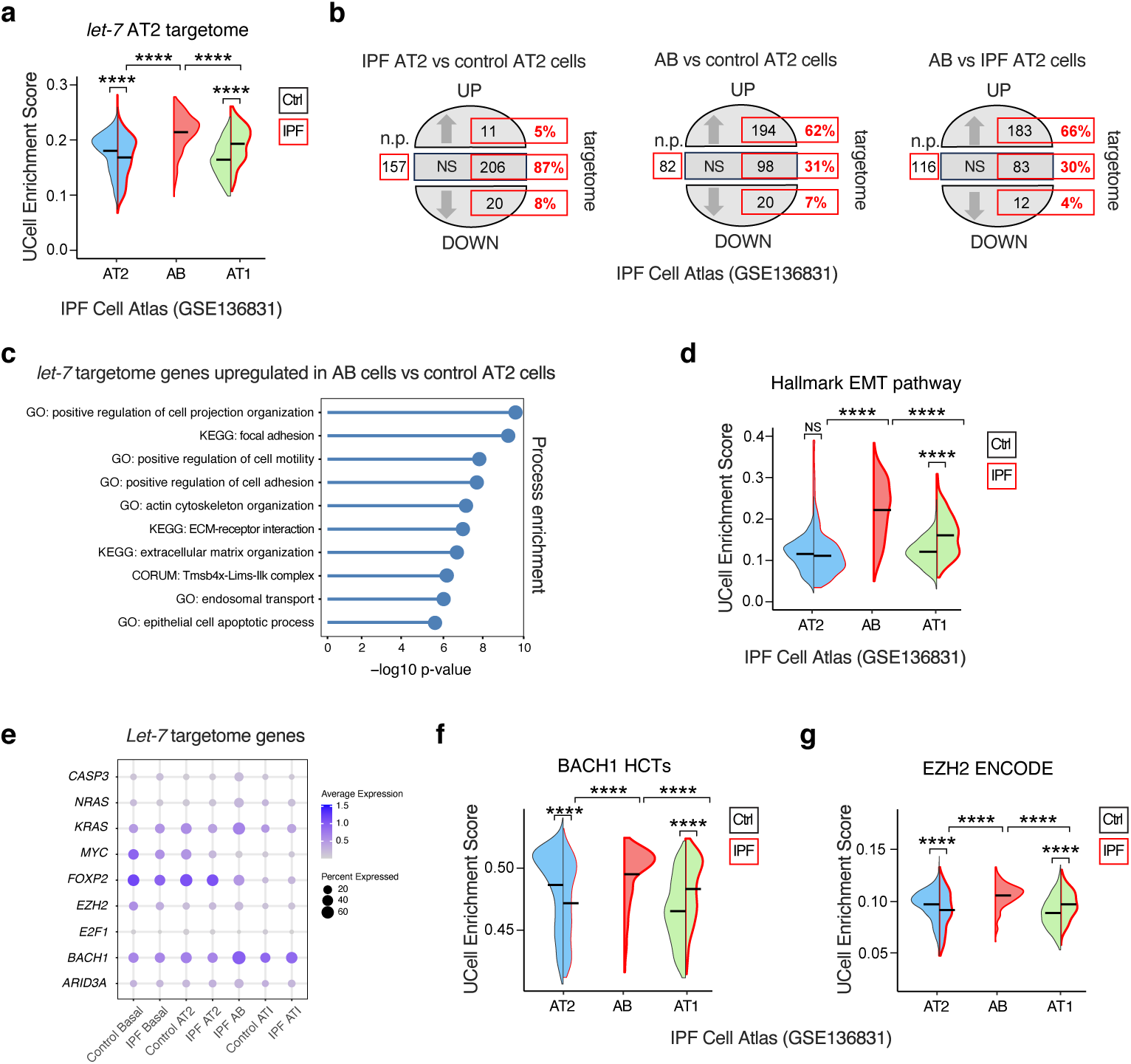
The *let-7* targetome is enriched in IPF “aberrant basaloid” (AB) cells. **a** UCell Enrichment scores for the murine *let-7* targetome in AT2, AB, and AT1 cells from control or IPF lungs were obtained from published scRNA-seq dataset GSE136831^14^. **b** Broken circle plots indicate the total number and proportions of *let-7* targets which are significantly upregulated (UP), downregulated (DOWN), or non significant (NS) between control (Ctrl) and IPF cells. Not present (n.p.) indicates gene targets missing in dataset GSE136831^14^. **c** Gene process enrichment analysis for upregulated *let-7* targets in AB cells compared to control AT2 cells. **d** The Hallmark EMT pathway enrichment scores in control vs IPF AT2, AB and AT1 cell samples. **e** Dot plots show the relative expression of selected *let-7* targets and identity markers in AT2, AB, and AT1 cells from control and IPF patients. **f,g** Consensus targets of BACH1 and EZH2 were evaluated for UCell Enrichment Scores in AT2, AB and AT1 cells in control and IPF patients. High confidence targets (HCT)s. **a**,**d**,**f,g** Data are mean±s.e.m half violin plots. ****p < 0.0001, by one way ANOVA with Sidak correction. Data was derived from published scRNA-seq dataset GSE136831^14^.

## DISCUSSION

Our study lends support to the importance of *let-7* miRNA in AT2s as a molecular gatekeeper in lung homeostasis, injury, and healing. We identified transient and coordinated downregulation of *let-7* family after bleomycin-induced lung injury, indicating a tightly regulated post-transcriptional response which paves the way for the emergence of repair-associated AT2s. This interpretation aligns with our studies indicating that AT2-cell specific genetic ablation of *let-7afd* removes a “brake” on AT2 activation resulting in uncontrolled cell growth and accumulation of ADIs in pulmonary hemorrhage which precedes a temporary chronic fibrotic remodeling phase. Additionally, we discovered that *let-7* not only restricts AT2 cells from transitioning into ADI intermediates but it is also crucial for exiting this state. Interestingly, with a booster Cre/loxP deletion protocol, we observed that ADIs and pulmonary fibrosis were maintained in *let-7afd^AT2^*mice. We speculate that extensive cell death and senescence of *let-7afd^-/-^* AT2s may secondarily recruit airway epithelial PSCs^12,50–52^ which can repair the parenchyma.

The *let-7* miRNA family exerts broad cellular effects by silencing hundreds of target mRNAs in GRNs^20^. Here we demonstrate that *let-7* simultaneously regulates an extensive GRN in AT2 cells, including oncogenes such as BACH1, EZH2, and MYC previously linked to uncontrolled cell growth and fibrosis^22–24,33,53^. Our data favors that this OGRN is persistently upregulated in AT2 cells due to impaired *let-7* post-transcriptional regulation which would support progressive pulmonary fibrosis disease. We note that the *let-7* OGRN likely functions as a powerful coherent or synergistic feedforward-loop because these genes converge on similar phenotypic disturbances in diseased epithelial cells. Impressively, OGRN *let-7* targets are enriched in cycling and pAT2 and ADIs but their expression decreases as cells progress toward AT1 identity. With permanent loss of *let-7afd*, our data indicates that AT2 cell identity is held within ADI state, where the *let-7* OGRN genes are most highly expressed.

The leading-edge targets of *let-7*, such as EZH2, BACH1, MYC, are associated with epithelial stem cell growth, transformation, and enhanced metastatic potential involving processes which include EMT^30,32,36^. Moreover, published studies on *let-7* have shown that its loss stimulates cell growth and EMT programs, while its overexpression limits these processes^20,21^. Intriguingly, BACH1, EZH2, and MYC have also been associated with partial EMT in cancer, where epithelial cells do not fully transition into mesenchymal cells but exhibit a continuum of epithelial and mesenchymal traits which potentiate metastasis^54^. Lineage tracing and scRNA-seq studies indicate that AT2 transitional cells do not transform into fibroblasts via EMT^10–13^; instead, they adopt a mixed epithelial and mesenchymal gene expression program. Recent studies showed that the accumulation of AT2 transitional cells can drive lung injury and ILD^11–13^. Given these observations, we propose a model where loss of *let-7afd* contributes to partial EMT in transitional AT2s via the OGRN.

Prior studies have highlighted the importance of BACH1, EZH2, and MYC on heritable chromatin remodeling leading to phenotypic changes in cells^44–46^. Overexpression of BACH1, EZH2, or MYC promote histone modifications including H3K27ac at active enhancers to facilitate oncogenic cell signaling^44–46^. Furthermore, recent studies have shown that BACH1 and EZH2 can promote H3K27me3 epigenetic silencing cooperatively in cells^47,55^. According to our data, the loss of *let-7afd* enhances H3K27ac modifications which may lead to opening of chromatin and increased accessibility of pioneer TFs to the enhancers of *let-7* targetome, thereby boosting not only mRNA and protein production post-transcriptionally but also transcription output. This likely amplifies the positive feedback loop with *Bach1*, *Ezh2*, *Myc* themselves acting as coherent chromatin remodeling TF drivers in the establishment of a profibrotic AT2 epigenetic landscape. However, the specific roles of BACH1, EZH2 and other OGRN targets as necessary and/or sufficient mediators in the persistence of ADIs and pulmonary fibrosis remain largely undetermined. Nonetheless, recent research on *Ezh2* has shown that its deletion hampers the growth of lung epithelial cells and reduces the emergence of cycling epithelial cells and KRT8^+^ transitional epithelial cells in murine organoids ^56^.

Our genetic studies to reduce *let-7* activity in AT2 cells relied primarily on phenotypic studies of *let-7afd* but did not provide a mechanistic perspective on the contribution of *let-7bc2* and we did not examine the role of other clusters in AT2 cells. Another limitation of our study is that although we demonstrated dynamic changes in AT2 cells upon loss of *let-7afd* by integrated approaches, we did not achieve transcriptomic and epigenomic depth at a single cell resolution which would have allowed more insight on the role of *let-7* on AT2 cell plasticity and alveolar cell niche.

In conclusion, our study provides insights into *let-7* as a braking mechanism to oncogene dysregulation in AT2 cell-driven pulmonary fibrosis. Our observation that AB cells from IPF patients express higher levels of the *let-7* targetome offers the enticing possibility that targeted delivery of *let-7* could be used as a treatment. We point that several reports indicate the effectiveness of targeted *let-7* delivery in mitigating fibrosis ^57,58^, and pharmacologic targeting of EZH2 is employed against both cancer and fibrosis ^59,60^. Furthermore, since AT2 cells are significant source of lung adenocarcinomas and IPF patients are at high risk of cancer; we propose that existing anti-cancer drugs might offer new approaches to target AB cells in patients with IPF.

## MATERIALS AND METHODS

### Mice and tamoxifen administration

Mice of both sexes were used for experiments. We recently described the creation and PCR genotyping strategy of the C57BL/6 isogenic *let-7bc2^fl/fl^* or *let-7afd^fl/fl^* mice^26^. Mice were crossed to *S*ftpc*^tm1(cre/ERT)Blh^* (*Sftpc-CreER^T2^*), *Rosa26R-CAG-LSL-tdTomato (R26tdT)* purchased from JAX and genotyped by PCR. For all phenotypic studies, mice aged 8-12 weeks old were given an initial iTAM regimen defined as 4 doses of 160 mg/kg tamoxifen (Sigma, T5649) dissolved in corn oil (Sigma, C8267) by intraperitoneal injections (i.p.) every other day. In some chronic phenotypic experiments, mice received iTAM boosters following four weeks after the initial iTAM. The boosters consisted of 4 doses of 80 mg/kg iTAM every other day and these were administered monthly. *Let-7bc2^f/f^*, *let-7afd^f/f^* or wild-type (WT) *Sftpc-tdT* mice were used as controls and treated with iTAM in parallel with experimental mice. Mice were genotyped by PCR from ear clippings with published primers^26^. All studies with mice were approved by the Baylor College of Medicine Institutional Animal Care and Use Committee and followed the National Research Council Guide for the Care and Use of Laboratory Animals.

### Physiologic measurements

Arterial oxygen saturation (SpO2) measurements were collected using the STARR Life Sciences MouseOx Plus (Oakmont, Pennsylvania) on awake mice with a neck cuff sensor to obtain five readings over a 30 min interval. The mice were shaved on the neck and allowed 45 minutes to acclimate without restrains with a neck cuff sensor prior to good quality recordings. The lung respiratory biomechanics values were obtained with the flexiVent (Scireq) system^61^. Mice undergoing flexiVent procedure were subjected to anesthesia, followed by tracheostomy and cannulation with an 18G metal cannula. The mice were placed under mechanical ventilation at a respiratory rate of 150 breaths/min, a tidal volume of 10 mL/kg and a PEEP set at 3cmH2O in enclosed chamber for pulmonary function measurements following manufacturer guidelines. The average of three measurements with a coefficient of determination ≥ 0.95 were calculated for each mouse.

### Lung histology and histomorphometry analysis

Mouse lungs were perfused with sterile PBS via right ventricle cardiac perfusion. For formalin-fixed paraffin embedding tissues, lungs were inflated with instillation of 4% paraformaldehyde (PFA) via a tracheal cannula at 25-cm H_2_O pressure for 15mins and then tied by nylon suture. The lungs were then transferred into 50 mL polypropylene tube containing PFA for overnight fixation at 4°C. Then lung lobes were separated, transferred into tissue cassettes, and then immersed in 70% ethanol at 4°C. Samples were provided to the Human Tissue Acquisition and Pathology (HTAP) Core at BCM for paraffin embedding, cutting of 5μM thick sections, and staining by H&E or Masson’s trichrome. The PFA-fixed lung cryosections were obtained in similar manner, but after the overnight fixation step in PFA at 4°C, the lungs were equilibrated with 30% sucrose for 1-2 days, prior to embedding in OCT. Samples were then cut into 7-10uM thick sections for the analysis. β-galactosidase (X-gal) lung staining was performed on frozen lung sections using the Senescence Cells Histochemical Staining Kit (Sigma CS0030) following manufacturer instructions. Hydroxyproline Colorimetric Assay Kit (Abcam, ab222941) was used to measure collagen in lung lobes following kit instructions and optical density was measured using a Varioskan LUX plate reader (Thermo Fisher Scientific, VL0000D0).

### Bronchoalveolar lavage and Prussian blue staining

Bronchoalveolar lavage for measurements of total and differential cell counts was done as we described ^26^. Prussian Blue staining of BALF cytospins was performed using the Iron Stain Kit (Sigma, HT20-1KT) following kit instructions.

### Immunostaining

For immunostaining of paraffin embedded lung tissues or organoid sections, antigen retrieval was performed with 1X BioCare Diva Decloaker RTU (BioCare, DV2004G1) buffer in a Decloaking Chamber NxGen (BioCare, DC2012) at 95°C or 110°C for 15mins. Slides were then allowed to cool in antigen retrieval buffer for 10 mins before washing in distilled water. Both paraffin embedded and thawed cryo-sectioned slides were permeabilized with 0.3% Triton X-100 in 1% Normal Goat Serum (Abcam, ab7481) for 10 mins. Antigen blocking was performed in 5% Normal Goat Serum for 1 hour. Samples were then incubated overnight at 4°C with the following primary antibodies diluted in 1% Normal Goat Serum: Rat anti-Ki67 conjugated FITC (1:50, Invitrogen, 11-5698-82), Mouse anti-active/pro-caspase3 (1:20, Invitrogen, MA1-91637), Mouse anti-EZH2 (1:100, Invitrogen, 14-9867-82), Rabbit anti-EZH2 (1:200, Cell Signaling, 5246), Rabbit anti-BACH1 (1:200, Novus Biological, NBP2-55113), Mouse anti-γH2AX (1:200, Novus Biological, NB100-74435), Rabbit anti-CLDN4 (1:200, Invitrogen, 36-4800), Rabbit anti-pro-SFTPC (1:200, Millipore, AB3786), Rabbit anti-RFP/tdTomato (1:200, Rockland, 600-401-379), Rat anti-Krt8/TROMA-I (1:20, DSHB, TROMA-I-s), Rat anti-Galectin-3/LGALS3 (1:500, Cedarlane, CL8942AP), Rat anti-RAGE/AGER (1:100, R&D Systems, MAB1179-100), Rabbit anti-H3K27me (1:200, Invitrogen, MA5-11198), Rabbit anti-Aqp5 (1:100, Invitrogen, PA5-36529), and Mouse anti-HopX (1:200, Santa Cruz, sc-398703). The next day, samples were incubated for 1 hour at room temperature in the following secondary antibodies diluted in 1% Normal Goat Serum with DAPI (1:10000, Sigma, D9542): Alexa Fluor 488 goat anti-mouse (1:2000, Thermo Fisher Scientific, A11029), Alexa Fluor 488 goat anti-Rabbit (1:2000, Thermo Fisher Scientific, A-32731), Alexa Fluor 488 goat anti-Rat (1:2000, Thermo Fisher Scientific, A11006), Alexa Fluor 555 goat anti-Mouse (1:2000, Thermo Fisher Scientific, A32727), Alexa Fluor 555 goat anti-Rabbit (1:2000, Thermo Fisher Scientific, A21428), Alexa Fluor 555 goat anti-Rat (1:2000, Thermo Fisher Scientific, A21434), Alexa Fluor 647 goat anti-Rabbit (1:2000, Thermo Fisher Scientific, A21244), Alexa Fluor 647 goat anti-Rat (1:2000, Thermo Fisher Scientific, A21247), Alexa Fluor 790 Goat anti-Rabbit (1:2000, Thermo Fisher Scientific, A11369). In some cases, paraffin embedded lung sections were treated after secondary staining with TrueBlack Lipofuscin Autofluorescence Quencher (Biotium, 23007) for 1 min following product instructions to block autofluorescence. Slides were mounted with ProLong Glass Antifade Mountant (Thermo, P36980).

### Lung cell suspensions and purification of AT2 cells

Isolation of *Sftpc-tdT^+^*lineage derived AT2s was performed as previously described with minor modifications ^27,62^. Briefly, mouse lungs were perfused with cold PBS via right ventricle of the heart and then inflated with 1 mL digestion solution containing 5 units/mL dispase (Corning, 354235), 450 units/mL collagenase type I (Gibco, 17100-017), 0.33 units/mL DNase I (Sigma, 10104159001), and 1X antibiotic-antimycotic (Thermo Fisher Scientific, A5955-100ML) in Advanced DMEM/F-12 (Thermo Fisher Scientific, 12634028). Lung lobes were then minced into 1-2 mm^3^ pieces, transferred into 4 mL digestion solution, and incubated at 37°C with rotation for 40 mins. Samples were dissociated by vigorously mixing by P1000 pipette tip until no large pieces remained. Samples were filtered through a 70µm strainer, pelleted, and then resuspended in 1 mL RBC Lysis Buffer (Invitrogen, 00-4333-57) for 1-min before washing with 5 mL Advanced DMEM/F-12 with 1X antibiotic-antimycotic. Lung cell suspensions were stained with Live/Dead Fixable Blue stain (Invitrogen, L34962), Alexa Fluor 700 anti-mouse CD45.2 (Biolegend, 109822), Brilliant Violet 711 anti-mouse EpCam (Biolegend, 118233), Alexa Fluor 700 anti-mouse CD31 (Biolegend, 102444), Brilliant Violet 421 anti-mouse Pdgfrα (Biolegend, 135923) antibodies and incubated for 30 minutes in dark on ice. Cells were suspended in ice cold 1X PBS with 2% FBS and filtered through a 40µm strainer followed by sorting of AT2 cells with *Sftpc-tdT* tracer on a FACSAria (BD Biosciences). The purity of sorted cells was >97% and it was confirmed via immunostaining with SFTPC (Supplemental Fig. 10). In initial AT2 organoid culture experiments, lung cell suspensions were incubated on antibody-coated dishes at 37°C with 5% CO_2_ for 45 mins to remove adherent cells and immune cells as an intermediate step prior to flow sorting of AT2 cells as detailed above. This intermediate step was done by coating 100mm tissue culture dishes with 42 µg anti-Mouse CD45 (BioLegend, 103101) and 16 µg anti-Mouse CD32 (BioLegend, 156402) in 7 mL PBS for 24-48 hours at 4°C as previously described.^62^

### RNA extraction and quantitative real-time PCR (qPCR)

Cell pellets or lung lobes from individual mice were snap frozen in 350µl of 1:100 β-mercaptoethanol in the RLT Lysis Buffer from the RNAeasy mini plus Kit (Qiagen, 74134). For extraction of RNA in bulk RNA-seq lineage labelled *Sftpc*-tdT^+^ AT2 cells (∼700,000) were pooled from three mice per genotype by flow sorting as indicated above. Cell samples were thawed and vortexed while lung samples were homogenized for total RNA extraction intended for qPCR or bulk RNA-seq. The RNAeasy mini plus Kit (Qiagen, 74134) was used for removal of genomic DNA and RNA isolation. Quality control was performed with NanoDrop 2000C Spectrophotometer (Thermo Fisher Scientific). Equal amount of input RNA was used for preparation of cDNA with the High-Capacity cDNA Reverse Transcription Kit with RNase Inhibitor (Thermo Fisher Scientific, 4374966). Pre-designed Taqman probes were used with TaqMan Fast Advanced Master Mix (Thermo Fisher Scientific, 4444557). *Gapdh* served to normalize gene expression by ΔΔCT method. The qPCR was conducted and analyzed on ViiA 7 System (Applied Biosystems). The following Taqman probes were used: *Bach1* (Mm01344527_m1), *Casp3* (Mm01195085_m1), *Cdkn2a* (Mm00494449_m1), *E2f1* (Mm00432939_m1), *Ezh2* (Mm00468464_m1), *Kras* (Mm00517492_m1), *Myc* (Mm00487804_m1), *Nras* (Mm03053787_s1), *Trp53* (Mm01731287_m1), *Gapdh* (Mm99999915_g1). The validated and published *pri-let-7b/c2* and *pri-let-7a1/d/f1* transcript Taqman probes ^26^ were custom designed by Applied Biosystems (44411114, areptx2, arfvmhy).

### AT2 3D organoid cultures and differentiation

Culture of 3D organoids was carried out as previously described with minor modifications ^27,63^. Lineage labeled *Sftpc*-tdT^+^ AT2 cells were purified from 8-10 week-old mice on day 6-day after iTAM treatment. AT2 cells were resuspended in thawed Cultrex UltiMatrix Matrigel (Biotechne, BME001-05) at 1,000 cells/µL. For each biological replicate, organoids were initiated in each well of 6-well plate into five 20 µl domes containing ∼20,000 cells and incubated at 37°C with 5% CO_2_ for 20 mins without media. When domes hardened, 2mL of alveolar maintenance media (AMM) without Interleukin-1β was added to each well. The organoids were fed AMM without Interleukin-1β every other day and grown for 10-14-days for subsequent analysis. For RNA-seq, the domes were washed with 1 mL sterile PBS and then incubated in 1 mL Cultrex Organoid Harvesting Solution (Biotechne, 3700-100-01) on ice for 1 hour. Domes were gently detached from 6-well plates by spatula, resuspended in 10 mL PBS, pooled for each biological replicate, and then pelleted at 300 x g for 10 mins at 4°C. Cells were washed and pelleted several times with 10 mL sterile PBS to remove Matrigel. In AT2 to AT1 differentiation experiments, organoids were switched from AMM into AT2-Differentiation medium (ADM) on day 10 followed by an additional 7-days as previously described ^27,63^. For immunostaining analysis, the AT2 organoids were fixed and sectioned as in previous publication^63^. Organoid colony forming efficiency (CFE) and alveolosphere size diameters were quantified using FIJI (ImageJ, v2.14.0).

### Bulk RNA-seq processing, pathway enrichment, and regulatory network analysis

Purified RNA (> 800 ng) from each of the samples was used for the bulk RNA-seq analysis. RNA samples with RIN scores greater than 7 were selected for RNA-seq with the Agilent Bioanalyzer 2100. Samples were submitted to the Genomic and RNA Profiling Core at BCM for RNA-seq and library preparation. Nonstrand-specific, polyA+ -selected RNA-seq libraries were generated using the Illumina TruSeq protocol. Libraries were sequenced to a median depth of 35-40 million 100-bp single-end reads on a NovaSeq sequencer (Illumina). The FASTQ files underwent quality control metrics (trimming), read mapping, and gene annotation (GRCm38/mm10 genome) with default parameters with the RNA-seq Alignment v2.0.2 pipeline from Illumina’s BaseSpace (https://basespace.illumina.com/). The RNA-Seq Differential Expression v1.0.1 workflow (Illumina BaseSpace) was used with default parameters to extract gene counts and to obtain differentially expressed genes (DEG)s with an adjusted p value (FDR) < 0.05 between control and experimental samples. Gene Ontology and pathway enrichment analysis was carried out with web based tools Enrichr (https://maayanlab.cloud/Enrichr/) ^64^ or with Metascape (https://metascape.org/) ^65^. The gene set enrichment analysis was performed with GSEA v4.2.3 Mac application using standard settings. The epithelial cell marker gene sets used for GSEA were obtained from Mouse Lung Injury & Regeneration web tool (https://schillerlabshiny.shinyapps.io/Bleo_webtool/) ^12^ (Supplementary Data 10). The Jaccard overlap scores for the *let-7* targetome were calculated by taking the size of the intersect and dividing it by the size of the union from epithelial cell types defined in GSE141259 ^12^. AT2 and ADI pseudotime trajectory values were obtained from Strunz et al.,^12^ Heatmaps and plots were generated with R (v2022.12.0). To predict transcription factors whose function is impacted by *let-7*, ChIP-Seq regulatory network analysis was performed on the *let-7afd*^-/-^ RNA-seq datasets as previously described ^66^. Published raw data files corresponding to total RNA-seq of purified AT2 cells from control vs bleomycin treated mice (n = 3 mice per group) (GSE115730) ^6^ were extracted from NCBI SRA toolkit and then processed for differential gene expression.

### Small RNA-seq data analysis

The published raw data files corresponding to small RNA-seq from lungs of mice after vehicle or bleomycin treatment (GSE195773)^25^ were extracted with the NCBI SRA toolkit. The SRA data files were analyzed with miRNA Analysis v1.0.0 application workflow (Illumina BaseSpace). To obtain mature *let-7* miRNA normalized counts, standard settings were used corresponding to murine GRCm38/mm10 genome and miRbase DB v21 database. Plots were generated with R (v2022.12.0) by normalizing bleomycin treated samples to untreated samples for each timepoint.

### CUT&RUN sequencing, processing, and data analysis

The CUT&RUN sequencing protocol was done as previously described ^67,68^. Lineage labelled *Sftpc*-tdT^+^ AT2 cells (∼100,000) were flow sorted from mice as indicated above. Cells were pooled from two mice per genotype for CUT&RUN sequencing (n = 2). Cells were washed with wash buffer (50 ml total H_2_O with 20 mM HEPES pH 7.5, 150 mM NaCl, 0.5 mM Spermidine, and one Roche Complete protein inhibitor tablet (Millipore, 11873580001). Cells were then incubated with activated concanavalin-A beads (Bang Laboratories, L200731C) for fifteen minutes at room temperature. Slurry was then separated with a Miltenyi MACSiMAG separator (Miltenyi 130-092-168), washed once, and resuspended in wash buffer containing 0.05% digitonin (Digitonin Wash buffer, Millipore, 300410). Digitonin wash buffer was used from here on out and cells kept on ice. The samples were placed on chilled tube rack kept on ice when not rotating for remainder of protocol. The samples were incubated with primary antibodies overnight at 4°C rotating and then washed three times and incubated on rotator with Digitonin wash buffer containing pAG-MNase (Epicypher 15-1016) for one hour at 4°C. Antibodies used were Rabbit anti-H3K27ac (Cell Signaling Technologies, 8173; 1:50 concentration), Rabbit anti-H3K427me3 (Cell Signaling Technologies, 9733; 1:50 concentration), and Rabbit anti-mouse IgG (Jackson ImmunoResearch, 315-005-003; 1:100). After, samples were washed three times they were resuspended in 100uL Digitonin wash buffer and chilled on ice rack for two minutes. 2μL of 100mM CaCl_2_ was added to each sample to catalyze MNase cleavage. Cells were then kept at 4°C for 40 minutes. Cleavage was stopped by adding equal volume stopping buffer (340 mM NaCl, 20 mM EDTA, 4 mM EGTA, 0.05% Digitonin, 0.05 mg/mL glycogen, 5 ug/mL RNase A, 2 pg/mL heterologous spike-in DNA (Epicypher 18-1401)), and samples were heated to 37°C for 20 minutes for MNase/Histone/DNA complex release into supernatant. Slurry was separated with magnetic stand, and supernatant with DNA was collected into new tubes. DNA was purified with phenol chloroform extraction (Thermo Fisher Scientific, 15593031) and resuspended in ultrapure water (Invitrogen, 10-977-023). The samples were then submitted for library generation and sequencing to Admerahealth (South Plainfield, NJ).

The amount of DNA was determined by Qubit 2.0 DNA HS Assay (ThermoFisher, Massachusetts, USA) and quality was assessed by Tapestation High sensitivity D1000 DNA Assay (Agilent Technologies, California, USA). The KAPA HyperPrep kit (Roche, Basel, Switzerland) was used for library preparation. Assessment of library quality and quantity was done with Qubit 2.0 DNA HS Assay (ThermoFisher, Massachusetts, USA), Tapestation High Sensitivity D1000 Assay (Agilent Technologies, California, USA) for sequencing on an Illumina HiSeq 4000 (Illumina, California, USA) with a read length configuration of 150 paired-end (PE).

CUT&RUN sequencing data was trimmed for low quality reads and Illumina adapters using the trim galore (0.6.10) package (https://github.com/FelixKrueger/TrimGalore). Data was mapped using bowtie2 against the mouse genome UCSC build mm10. Peaks were called using MACS2 with IgG serving as control in each sample. Differential peaks between control and *let-7afd^-/-^* samples were determined using DiffReps, with a significance cutoff and FDR adjusted q value < 0.05 and fold change exceeding 1.5x for H3K27ac peaks and a q value of < 0.10 with fold change exceeding 1.5x for H3K27me3 peaks. Signal tracks were generated using DeepTools v2.1.0 https://deeptools.readthedocs.io/en/2.1.0/index.html) and BEDTOOLS v2.31.0 suite (https://bedtools.readthedocs.io/en/latest/content/bedtools-suite.html) was used for annotation of differential peaks overlapping genes +/-10kb from its genebody ^69^. GO terms for differential peaks were determined using GREAT v4.0.4 ^70^. Enriched motifs were derived using the findMotifsGenome.pl HOMER tool ^71^. Data peaks were visualized with the UCSC genome browser.

### AGO2 eCLIP and *let-7* interactome network analysis

To obtain lung samples for AGO2-eCLIP+let-7, eight-week-old wild-type C57BL/6 mice were administered with constant flow of 3.5% isoflurane during intratracheal administration of 4U/kg bleomycin. The mice were then euthanized 6-days after treatment. Mouse lungs were cleared by cardiac perfusion and then inflated with 1mL RNAse free PBS prior to harvesting and snap freezing with liquid nitrogen. Samples were provided to EclipseBio (San Diego, California) for generation of AGO2-eCLIP+let-7 libraries. The AGO-eCLIP+let-7 was performed in duplicate from 2 lungs of mice with chimeric ligation of miRNA and mRNA as previously described but with following modifications for lung sample processing and the enrichment of *let-7* targets ^28,72^. Briefly, approximately 20×10^6^ cells from the lung were plated on a dish then UV crosslinked, then pelleted and suspended in 1 ml volume of eCLIP lysis mix. The lysates were then sonicated (QSonica Q800R2) for 5 minutes, 30 seconds on / 30 seconds off with an energy setting of 75% amplitude, followed by digestion with RNase-I (Ambion). A primary mouse monoclonal AGO2/EIF2C2 antibody (sc-53521, Santa Cruz Biotechnology) was pre-coupled for 1hr with Sheep Anti-Mouse IgG Dynabeads (Thermo Fisher Scientific, 11202D) and then added to the homogenized lysate for a 2hr immunoprecipitation (IP) at 4°C. After the IP, a 2% portion of the sample was taken as paired Input and the remainder of lysate washed with eCLIP high stringency wash buffers. The miRNA:mRNA chimeric ligation step was done at room temperature for 1hr with T4 RNA ligase (NEB). The IP samples were then treated with alkaline phosphatase (FastAP, Thermo Fisher) and then T4 PNK (NEB) for ligation of barcoded RNA adapters and isolation of AGO2-RNA complexes ^28,72^. To generate AGO2-eCLIP libraries enriched for *let-7* chimeras, 5’ biotinylated ssDNA probes antisense to murine *let-7a-5p*, *let-7b-5p*, *let-7d-5p*, *let-7f-5p* miRNAs were ordered from Integrated DNA Technologies and resuspended in nuclease-free water to a final concentration of 100µM. Each probe was bound individually to Capture Beads (Eclipse Bioinnovations) and afterward mixed. The resulting probe-bound bead mixture was added to the reserved AGO2-eCLIP enrichment fractions and allowed to hybridize over a 1.5-hour period, with incrementally decreasing temperatures. Beads were washed to remove nonspecific RNAs and afterward, bound RNAs were eluted from beads using a DNase treatment. Eluted RNA was purified using an RNA Clean and Concentrator-5 Kit (Zymo Research, cat. #R1015/R1016) and taken through reverse transcription, DNA adapter ligation, and PCR amplification to generate sequencing libraries. Library sequencing was performed as SE122 on the NextSeq 2000 platform. BedGraph files of *let-7* chimeric peaks were generated for visualization with UCSC genome browser.

The AGO2-eCLIP+let-7 processing, and bioinformatics analysis was done by Eclipsebio (San Diego, California) with a proprietary analysis pipeline v1 developed from several published eCLIP publications ^28,72^. The original bioinformatic pipelines are available (https://github.com/yeolab/eclip; https://github.com/YeoLab/chim-eCLIP). A brief description follows: The umi_tools (v1.1.1) tool was used to prune Unique molecular identifier (UMIs) from reads. Next, cutadapt (v3.2) was used for trimming 3’ adapters. Repetitive elements and rRNA sequences were removed and non-repeat mapped reads mapped to murine GRCm38/mm10 using STAR (v2.7.7a). PCR duplicates were removed with umi_tools (v1.1.1). miRbase (v22.1) miRNAs were reverse mapped to reads with bowtie (v1.2.3) and miRNA portion of each read trimmed. STAR (v2.7.7a) was used to map the miRNA portion of each read to the genome. PCR duplicates were trimmed, and miRNA target cluster peaks were identified with CLIPper (v2.1.0). Individual clusters were annotated with corresponding miRNAs names and peaks annotated to transcripts GENCODE release M25 (GRCm38.p6). Peaks present in both biological samples were selected for downstream analysis. De novo HOMER (v4.11) ^71^ motif analysis was also applied to confirm the *let-7* seed region as the most enriched motif in dataset. *Let-7* chimeric peaks with overlap to the 3’UTR regions of protein coding genes were designated as the AGO2-eCLIP *let-7* interactome. The Sylamer test was done as previously described^73^. Upregulated (p < 0.05) genes in *let-7afd^-/-^ Sftpc*-tdT and/or cultured organoids were overlayed with AGO2-eCLIP *let-7* interactome and experimentally validated and published *let-7* targets in miRTarbase database ^74^ with Excel software (Microsoft). Pathway enrichment analysis for the *let-7* interactome was carried out with Enrichr ^64^ and Metascape ^65^. Protein-protein interaction networks were derived from Metascape (using parameter of Combined Core) and then visualized and organized with EnrichmentMap/Cytoscape (v3.10.1).

### Human IPF Atlas scRNA-seq integration

A published scRNA-seq dataset deposited in IPF Cell Atlas from control and IPF patients was used^14^. The cell annotations for cell clusters were provided by the authors ^14^. Cell-level expression for individual genes was obtained with R package Seurat ^75^. Gene signature scores were generated using the UCell method. Violin plots for individual genes or gene signatures in specific cell types were generated using the R statistical package v4.4.0. Dotplot expression of individual gene in selected cell types were generated using Seurat ^75^. The Hallmark EMT and ENCODE Consensus ChIP-X gene targets for EZH2 used for UCell enrichment were obtained from Enrichr^64^. BACH1 high confidence transcriptional targets were derived with the ChIP-Seq regulatory network toolkit ^66^.

### Transmission electron microscopy (TEM) and analysis

We utilized a published protocol for fixation ^6^, processing and embedding of lung tissues for preservation of AT2 cell lamellar body (LB) ultrastructure. Mice were euthanized, followed by an incision of the IVC artery and descending aorta to allow for cardiac draining of blood. Lungs were inflated with 1.2 mL of freshly prepared room temperature TEM fixative (4% paraformaldehyde, 2.5% glutaraldehyde, 0.02% picric acid in 0.1 M sodium cacodylate buffer, pH 7.3), tie sutured and then fixed overnight at 4°C. Strips of lung tissue were dissected, washed in 0.1M sodium cacodylate buffer, fixed in a solution of 1% OsO_4_ and 1.5% potassium ferricyanide for 1hr with gentle agitation, washed several times with 0.1M sodium cacodylate buffer and then incubated overnight at 4 °C. Samples were then washed with ddH_2_0, stained with 1.5% uranyl acetate in the dark for 1hr, dehydrated with a graded series of ethanol, infiltrated with and embedded in EMbed812 epoxy resin (#14120, Electron Microscopy Sciences) and then sectioned at 100 nm thickness with a Leica EM UC7 ultramicrotome. Images were collected with a JEOL JEM-1400Flash TEM operating at 120 kV of accelerating voltage and equipped with a high-contrast pole piece and an AMT NanoSprint15 Mk-II sCMOS camera. Images were examined with FIJI (ImageJ, v2.9.0) to quantify AT2 cell area (mm2), LB area (mm2), LB number/cell area (#/mm2) and AT2 cell circularity.

### Microscopy, image acquisition, and quantification

Slides were imaged on a Leica DM6000B (Leica, Wetzlar, Germany) using cameras CoolSNAP HQ2 (Teledyne Photometrics, USA) or Infinity3-6UR (Teledyne Lumenera, USA). A Keyence BZ-X800 microscope (Keyence, USA), an Olympus IX83 microscope (Olympus, USA), and a Nikon A 1Rs confocal microscope were used for imaging. Image processing was done with BZ-X Series Advanced Analysis software (Keyence) or cellSens image deconvolution software (Olympus). Brightfield color images captured by Leica microscope where white balance with FIJI (ImageJ, v2.14.0), and then stitched using AIVIA software (Leica, v11.0). Organoid domes were imaged on 6-well plates with a BioTek LionHeart LX microscope (Agilent, USA). Images were captured with 10X objective and stitched with built-in software. Cell counts were done manually based on markers and DAPI with FIJI software. Mean Fluorescence Intensity (MFI) was measured by FIJI software. Images of interest where then prepared using Adobe Photoshop and Illustrator. Mean linear intercept and Ashcroft scoring measurements of mouse lung morphometry were done as previously described^26,76^.

### Statistical analysis

Values are represented as indicated in figure legends. P values were calculated using two-tailed unpaired or paired Student’s t test. Nonparametric Rank test was used for Ashcroft Scoring. 1-way ANOVA with Sidak post-hoc test was used for the UCell enrichment analysis. A two-sided p value < 0.05 was statistically significant. Determination of sample size for animal experiments were based on pilot tests. Investigators were not blinded in the allocation of experiments. No data was excluded from analysis. Mice of both genders were randomly assigned into experimental or control groups for experiments. Statistical analyses were performed using GraphPad Prism (v9.5.1) except for the RNA-seq and CUT&RUN which were done as described above.

## ACKNOWLEDGEMENTS

We thank Drs. David Corry and Farrah Kheradmand for valuable scientific suggestions and for generous sharing of lab equipment. M. Sayeeduddin for histology at the Tissue Acquisition and Pathology Core (funded in part by P30 CA125123); and Joel M. Sederstrom for flow cytometry at the BCM and Cell Sorting Core (funded in part from NIH (CA125123 and RR024574); we thank the Texas Children’s Hospital William T Shearer Center for Human Immunobiology and Dr. Ronald Parchem for access to microscope equipment. Bioinformatics analysis was partially supported by The Cancer Prevention Institute of Texas (CPRIT) grants RP210227 and RP200504, NIEHS grants P30 ES030285 and P42 ES027725. This work was supported by grants from the NHLBI (R01HL140398, HL167814 to AR; HL155672 to KYK; F31 HL164287 to BTT), and NIGMS (T32 GM136554 to MS).

## AUTHOR CONTRIBUTIONS

M.J.S., M.S. and A.R. conceptualized experiments, interpreted data, and wrote the manuscript. M.J.S., M.S., P.A.E., B.T.T, S.L.L. carried out experiments. M.D.M. carried out the TEM imaging. A.F.C. provided microscope image acquisition expertise. M.E.R-E., R.S.K. L.M.S., S.C.W., S.A.O. performed bioinformatic data analysis. K.Y.K. provided expertise and supervision of epigenomics experiments. A.E., I.O.R., N.J.M., C.C., provided expertise and co-supervised bioinformatic data analysis.

## DATA AVAILABILITY

The authors declare that all data supporting the findings of this study are available within the article and its Supplementary information files or from the corresponding author upon reasonable request. AGO2-eCLIP+let-7, bulk RNA-seq, and CUT&RUN-seq data has been deposited in the NCBI Gene Expression Omnibus database under accession code GSExxxx.

## REFERENCES

1 Basil, M. C. et al. The Cellular and Physiological Basis for Lung Repair and Regeneration: Past, Present, and Future. Cell Stem Cell 26, 482–502 (2020). 10.1016/j.stem.2020.03.009

2 Hogan, B. & Tata, P. R. Cellular organization and biology of the respiratory system. Nat Cell Biol (2019). 10.1038/s41556-019-0357-7

3 Basil, M. C., Alysandratos, K. D., Kotton, D. N. & Morrisey, E. E. Lung repair and regeneration: Advanced models and insights into human disease. Cell Stem Cell 31, 439–454 (2024). 10.1016/j.stem.2024.02.009

4 Wuyts, W. A. et al. Idiopathic Pulmonary Fibrosis: Best Practice in Monitoring and Managing a Relentless Fibrotic Disease. Respiration 99, 73–82 (2020). 10.1159/000504763

5 Katzen, J. & Beers, M. F. Contributions of alveolar epithelial cell quality control to pulmonary fibrosis. J Clin Invest 130, 5088–5099 (2020). 10.1172/JCI139519

6 Chung, K. P. et al. Mitofusins regulate lipid metabolism to mediate the development of lung fibrosis. Nat Commun 10, 3390 (2019). 10.1038/s41467-019-11327-1

7 Barkauskas, C. E. et al. Type 2 alveolar cells are stem cells in adult lung. J Clin Invest 123, 3025–3036 (2013). 10.1172/JCI68782

8 Nabhan, A. N., Brownfield, D. G., Harbury, P. B., Krasnow, M. A. & Desai, T. J. Single-cell Wnt signaling niches maintain stemness of alveolar type 2 cells. Science 359, 1118–1123 (2018). 10.1126/science.aam6603

9 Zacharias, W. J. et al. Regeneration of the lung alveolus by an evolutionarily conserved epithelial progenitor. Nature 555, 251–255 (2018). 10.1038/nature25786

10 Choi, J. et al. Inflammatory Signals Induce AT2 Cell-Derived Damage-Associated Transient Progenitors that Mediate Alveolar Regeneration. Cell Stem Cell 27, 366–382 e367 (2020). 10.1016/j.stem.2020.06.020

11 Kobayashi, Y. et al. Persistence of a regeneration-associated, transitional alveolar epithelial cell state in pulmonary fibrosis. Nat Cell Biol 22, 934–946 (2020). 10.1038/s41556-020-0542-8

12 Strunz, M. et al. Alveolar regeneration through a Krt8+ transitional stem cell state that persists in human lung fibrosis. Nat Commun 11, 3559 (2020). 10.1038/s41467-020-17358-3

13 Wang, F. et al. Regulation of epithelial transitional states in murine and human pulmonary fibrosis. J Clin Invest 133 (2023). 10.1172/JCI165612

14 Adams, T. S. et al. Single-cell RNA-seq reveals ectopic and aberrant lung-resident cell populations in idiopathic pulmonary fibrosis. Sci Adv 6, eaba1983 (2020). 10.1126/sciadv.aba1983

15 Pandit, K. V. et al. Inhibition and role of let-7d in idiopathic pulmonary fibrosis. Am J Respir Crit Care Med 182, 220–229 (2010). 10.1164/rccm.200911-1698OC

16 Kim, S. et al. Integrative phenotyping framework (iPF): integrative clustering of multiple omics data identifies novel lung disease subphenotypes. BMC Genomics 16, 924 (2015). 10.1186/s12864-015-2170-4

17 Kim, H. H. et al. HuR recruits let-7/RISC to repress c-Myc expression. Genes Dev 23, 1743–1748 (2009). 10.1101/gad.1812509

18 Kong, D. et al. Loss of let-7 up-regulates EZH2 in prostate cancer consistent with the acquisition of cancer stem cell signatures that are attenuated by BR-DIM. PLoS One 7, e33729 (2012). 10.1371/journal.pone.0033729

19 Johnson, S. M. et al. RAS is regulated by the let-7 microRNA family. Cell 120, 635–647 (2005). 10.1016/j.cell.2005.01.014

20 Ma, Y., Shen, N., Wicha, M. S. & Luo, M. The Roles of the Let-7 Family of MicroRNAs in the Regulation of Cancer Stemness. Cells 10 (2021). 10.3390/cells10092415

21 Jin, B. et al. Let-7 inhibits self-renewal of hepatocellular cancer stem-like cells through regulating the epithelial-mesenchymal transition and the Wnt signaling pathway. BMC Cancer 16, 863 (2016). 10.1186/s12885-016-2904-y

22 Liu, Y. & Zheng, Y. Bach1 siRNA attenuates bleomycin-induced pulmonary fibrosis by modulating oxidative stress in mice. Int J Mol Med 39, 91–100 (2017). 10.3892/ijmm.2016.2823

23 Le, H. Q. et al. An EZH2-dependent transcriptional complex promotes aberrant epithelial remodelling after injury. EMBO Rep 22, e52785 (2021). 10.15252/embr.202152785

24 Qin, H. et al. C-MYC induces idiopathic pulmonary fibrosis via modulation of miR-9-5p-mediated TBPL1. Cell Signal 93, 110274 (2022). 10.1016/j.cellsig.2022.110274

25 Strobel, B. et al. Time and phenotype-dependent transcriptome analysis in AAV-TGFbeta1 and Bleomycin-induced lung fibrosis models. Sci Rep 12, 12190 (2022). 10.1038/s41598-022-16344-7

26 Erice, P. A. et al. Downregulation of Mirlet7 miRNA family promotes Tc17 differentiation and emphysema via de-repression of RORgammat. Elife 13 (2024). 10.7554/eLife.92879

27 Konishi, S., Tata, A. & Tata, P. R. Defined conditions for long-term expansion of murine and human alveolar epithelial stem cells in three-dimensional cultures. STAR Protoc 3, 101447 (2022). 10.1016/j.xpro.2022.101447

28 Van Nostrand, E. L. et al. Robust transcriptome-wide discovery of RNA-binding protein binding sites with enhanced CLIP (eCLIP). Nat Methods 13, 508–514 (2016). 10.1038/nmeth.3810

29 Rodriguez, A. et al. Requirement of bic/microRNA-155 for normal immune function. Science 316, 608–611 (2007). 10.1126/science.1139253

30 Batool, A., Jin, C. & Liu, Y. X. Role of EZH2 in cell lineage determination and relative signaling pathways. Front Biosci (Landmark Ed) 24, 947–960 (2019). 10.2741/4760

31 Comet, I., Riising, E. M., Leblanc, B. & Helin, K. Maintaining cell identity: PRC2-mediated regulation of transcription and cancer. Nat Rev Cancer 16, 803–810 (2016). 10.1038/nrc.2016.83

32 Dhanasekaran, R. et al. The MYC oncogene - the grand orchestrator of cancer growth and immune evasion. Nat Rev Clin Oncol 19, 23–36 (2022). 10.1038/s41571-021-00549-2

33 Zhang, X. et al. Bach1: Function, Regulation, and Involvement in Disease. Oxid Med Cell Longev 2018, 1347969 (2018). 10.1155/2018/1347969

34 Hou, W., Tian, Q., Steuerwald, N. M., Schrum, L. W. & Bonkovsky, H. L. The let-7 microRNA enhances heme oxygenase-1 by suppressing Bach1 and attenuates oxidant injury in human hepatocytes. Biochim Biophys Acta 1819, 1113–1122 (2012). 10.1016/j.bbagrm.2012.06.001

35 Yang, R. et al. E2F7-EZH2 axis regulates PTEN/AKT/mTOR signalling and glioblastoma progression. Br J Cancer 123, 1445–1455 (2020). 10.1038/s41416-020-01032-y

36 Igarashi, K., Nishizawa, H., Saiki, Y. & Matsumoto, M. The transcription factor BACH1 at the crossroads of cancer biology: From epithelial-mesenchymal transition to ferroptosis. J Biol Chem 297, 101032 (2021). 10.1016/j.jbc.2021.101032

37 Wang, X. et al. Bach1 Induces Endothelial Cell Apoptosis and Cell-Cycle Arrest through ROS Generation. Oxid Med Cell Longev 2016, 6234043 (2016). 10.1155/2016/6234043

38 Zeidler, M. et al. The Polycomb group protein EZH2 impairs DNA repair in breast epithelial cells. Neoplasia 7, 1011–1019 (2005). 10.1593/neo.05472

39 Zheng, W. & Yu, A. EZH2-mediated suppression of lncRNA-LET promotes cell apoptosis and inhibits the proliferation of post-burn skin fibroblasts. Int J Mol Med 41, 1949–1957 (2018). 10.3892/ijmm.2018.3425

40 Gonzalez, M. E. et al. Histone methyltransferase EZH2 induces Akt-dependent genomic instability and BRCA1 inhibition in breast cancer. Cancer Res 71, 2360–2370 (2011). 10.1158/0008-5472.CAN-10-1933

41 Kuzyk, A. & Mai, S. c-MYC-induced genomic instability. Cold Spring Harb Perspect Med 4, a014373 (2014). 10.1101/cshperspect.a014373

42 Yao, C. et al. Senescence of Alveolar Type 2 Cells Drives Progressive Pulmonary Fibrosis. Am J Respir Crit Care Med 203, 707–717 (2021). 10.1164/rccm.202004-1274OC

43 Sato, M. et al. BACH1 Promotes Pancreatic Cancer Metastasis by Repressing Epithelial Genes and Enhancing Epithelial-Mesenchymal Transition. Cancer Res 80, 1279–1292 (2020). 10.1158/0008-5472.CAN-18-4099

44 Wei, Z. et al. MYC reshapes CTCF-mediated chromatin architecture in prostate cancer. Nat Commun 14, 1787 (2023). 10.1038/s41467-023-37544-3

45 Kim, J. et al. Polycomb- and Methylation-Independent Roles of EZH2 as a Transcription Activator. Cell Rep 25, 2808–2820 e2804 (2018). 10.1016/j.celrep.2018.11.035

46 Niu, C. et al. BACH1 recruits NANOG and histone H3 lysine 4 methyltransferase MLL/SET1 complexes to regulate enhancer-promoter activity and maintains pluripotency. Nucleic Acids Res 49, 1972–1986 (2021). 10.1093/nar/gkab034

47 Wei, X. et al. Bach1 regulates self-renewal and impedes mesendodermal differentiation of human embryonic stem cells. Sci Adv 5, eaau7887 (2019). 10.1126/sciadv.aau7887

48 Weber, F., Treeck, O., Mester, P. & Buechler, C. Expression and Function of BMP and Activin Membrane-Bound Inhibitor (BAMBI) in Chronic Liver Diseases and Hepatocellular Carcinoma. Int J Mol Sci 24 (2023). 10.3390/ijms24043473

49 Jiang, Y. et al. Histone H3K27 methyltransferase EZH2 and demethylase JMJD3 regulate hepatic stellate cells activation and liver fibrosis. Theranostics 11, 361–378 (2021). 10.7150/thno.46360

50 Hu, Y. et al. Airway-derived emphysema-specific alveolar type II cells exhibit impaired regenerative potential in COPD. Eur Respir J 64 (2024). 10.1183/13993003.02071-2023

51 Lv, Z. et al. Alveolar regeneration by airway secretory-cell-derived p63(+) progenitors. Cell Stem Cell 31, 1685–1700 e1686 (2024). 10.1016/j.stem.2024.08.005

52 Liu, Q. et al. Lung regeneration by multipotent stem cells residing at the bronchioalveolar-duct junction. Nat Genet 51, 728–738 (2019). 10.1038/s41588-019-0346-6

53 Shen, Y. et al. c-Myc promotes renal fibrosis by inducing integrin alphav-mediated transforming growth factor-beta signaling. Kidney Int 92, 888–899 (2017). 10.1016/j.kint.2017.03.006

54 Luond, F. et al. Distinct contributions of partial and full EMT to breast cancer malignancy. Dev Cell 56, 3203–3221 e3211 (2021). 10.1016/j.devcel.2021.11.006

55 Yuan, Z. et al. Knockdown of Bach1 protects periodontal bone regeneration from inflammatory damage. J Cell Mol Med 27, 3465–3477 (2023). 10.1111/jcmm.17916

56 Byrd, A. L. et al. Dysregulated Polycomb Repressive Complex 2 contributes to chronic obstructive pulmonary disease by rewiring stem cell fate. Stem Cell Reports 18, 289–304 (2023). 10.1016/j.stemcr.2022.11.009

57 Sun, L. et al. Exosomal miRNA Let-7 from Menstrual Blood-Derived Endometrial Stem Cells Alleviates Pulmonary Fibrosis through Regulating Mitochondrial DNA Damage. Oxid Med Cell Longev 2019, 4506303 (2019). 10.1155/2019/4506303

58 Huleihel, L. et al. Let-7d microRNA affects mesenchymal phenotypic properties of lung fibroblasts. Am J Physiol Lung Cell Mol Physiol 306, L534–542 (2014). 10.1152/ajplung.00149.2013

59 Duan, R., Du, W. & Guo, W. EZH2: a novel target for cancer treatment. J Hematol Oncol 13, 104 (2020). 10.1186/s13045-020-00937-8

60 Bao, X. et al. Inhibition of EZH2 prevents acute respiratory distress syndrome (ARDS)-associated pulmonary fibrosis by regulating the macrophage polarization phenotype. Respir Res 22, 194 (2021). 10.1186/s12931-021-01785-x

61 McGovern, T. K., Robichaud, A., Fereydoonzad, L., Schuessler, T. F. & Martin, J. G. Evaluation of respiratory system mechanics in mice using the forced oscillation technique. J Vis Exp, e50172 (2013). 10.3791/50172

62 Chen, Q. & Liu, Y. Isolation and culture of mouse alveolar type II cells to study type II to type I cell differentiation. STAR Protoc 2, 100241 (2021). 10.1016/j.xpro.2020.100241

63 Katsura, H. et al. Human Lung Stem Cell-Based Alveolospheres Provide Insights into SARS-CoV-2-Mediated Interferon Responses and Pneumocyte Dysfunction. Cell Stem Cell 27, 890–904 e898 (2020). 10.1016/j.stem.2020.10.005

64 Kuleshov, M. V. et al. Enrichr: a comprehensive gene set enrichment analysis web server 2016 update. Nucleic Acids Res 44, W90–97 (2016). 10.1093/nar/gkw377

65 Zhou, Y. et al. Metascape provides a biologist-oriented resource for the analysis of systems-level datasets. Nat Commun 10, 1523 (2019). 10.1038/s41467-019-09234-6

66 Ochsner, S. A. et al. The Signaling Pathways Project, an integrated ’omics knowledgebase for mammalian cellular signaling pathways. Sci Data 6, 252 (2019). 10.1038/s41597-019-0193-4

67 Skene, P. J., Henikoff, J. G. & Henikoff, S. Targeted in situ genome-wide profiling with high efficiency for low cell numbers. Nat Protoc 13, 1006–1019 (2018). 10.1038/nprot.2018.015

68 Le, D. T. et al. BATF2 promotes HSC myeloid differentiation by amplifying IFN response mediators during chronic infection. iScience 26, 106059 (2023). 10.1016/j.isci.2023.106059

69 Trevino, L. S. et al. Epigenome environment interactions accelerate epigenomic aging and unlock metabolically restricted epigenetic reprogramming in adulthood. Nat Commun 11, 2316 (2020). 10.1038/s41467-020-15847-z

70 McLean, C. Y. et al. GREAT improves functional interpretation of cis-regulatory regions. Nat Biotechnol 28, 495–501 (2010). 10.1038/nbt.1630

71 Heinz, S. et al. Simple combinations of lineage-determining transcription factors prime cis-regulatory elements required for macrophage and B cell identities. Mol Cell 38, 576–589 (2010). 10.1016/j.molcel.2010.05.004

72 Krivdova, G. et al. Identification of the global miR-130a targetome reveals a role for TBL1XR1 in hematopoietic stem cell self-renewal and t(8;21) AML. Cell Rep 38, 110481 (2022). 10.1016/j.celrep.2022.110481

73 van Dongen, S., Abreu-Goodger, C. & Enright, A. J. Detecting microRNA binding and siRNA off-target effects from expression data. Nat Methods 5, 1023–1025 (2008). 10.1038/nmeth.1267

74 Huang, H. Y. et al. miRTarBase update 2022: an informative resource for experimentally validated miRNA-target interactions. Nucleic Acids Res 50, D222–D230 (2022). 10.1093/nar/gkab1079

75 Satija, R., Farrell, J. A., Gennert, D., Schier, A. F. & Regev, A. Spatial reconstruction of single-cell gene expression data. Nat Biotechnol 33, 495–502 (2015). 10.1038/nbt.3192

76 Hubner, R. H. et al. Standardized quantification of pulmonary fibrosis in histological samples. Biotechniques 44, 507–511, 514-507 (2008). 10.2144/000112729

