## Supplementary Fig. 1 for "*Let-7* restrains an oncogenic circuit in AT2 cells to prevent fibrogenic cell intermediates in pulmonary fibrosis"

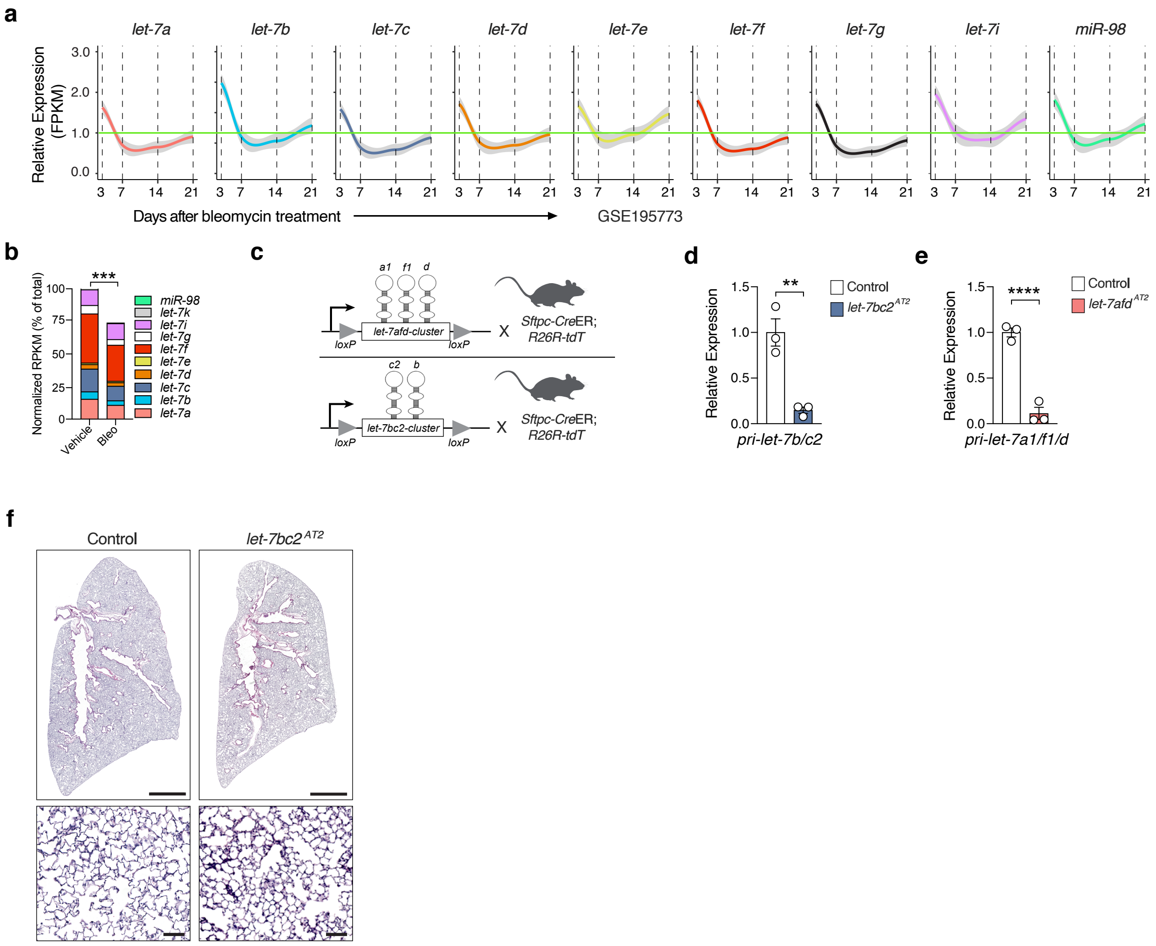


**Supplementary Fig 1** Temporal downregulation of *let-7* family after bleomycin induced lung injury. **a** Line plots show smoothed relative expression levels of *let-7-5p* family members in whole lungs of mice following 3, 7, 14, and 21 days of instillation with vehicle or bleomycin. Data was derived from published GSE195773 small RNA-seq dataset and is plotted relative to control lung samples at each time point. Data corresponds to individual lung samples of mice: day 3 (5 control; 7 bleomycin), day 7 (n=5 control; n=8 bleomycin), day 14 (6 control; 9 bleomycin), day 21 (6 control; 7 bleomycin), day 28 (7 control; 6 bleomycin). Bright green line corresponds to n = 1.0 fold-change relative to control samples. Gray area represents 95% confidence interval derived by smoothing fit. **b** Normalized RPKM values for each *let-7-5p* family member was tabulated as percentage of total lung *let-7* at the 7-day time point in vehicle treated (n = 5 mice) vs bleomycin (bleo) treated (n = 8) mice. Data are RPKM reads for individual *let-7* members adjusted as a percent of total reads. ***p < 0.001, by unpaired Student's t test. **c** Schematic representation for the *let-7afd* and *let-7bc2* cluster floxed alleles; also shown are mice used in this study: *let-7bc2*$^{f/f}$;*Sftpc*-CreERT2/+, *let-7afd*$^{f/f}$;*Sftpc*-CreERT2/+ mice with and without the R26R-LSL-tdTomato reporter (R26R-tdT). **d,e** The long noncoding RNA-like primary transcript for the *let-7bc2* cluster (*pri-let-7b/c2*) (**d**) or *let-7afd* cluster (*pri-let-7a1/f1/d*) (**e**) were detected by qPCR from experimental or control *Sftpc*-tdT$^{+}$ flow sorted AT2 cells following 6 days of iTAM (n = 3 mice per group). Data are mean±s.e.m. ****p < 0.0001, by unpaired Student's t test. **f** Representative H&E-stained lung lobe sections of *let-7bc2*$^{AT2}$ and control mice following 6-days of iTAM. Scale bars: 2 mm upper panels; 50 μm lower panels.
