## Supplementary Fig. 2 for "*Let-7* restrains an oncogenic circuit in AT2 cells to prevent fibrogenic cell intermediates in pulmonary fibrosis"

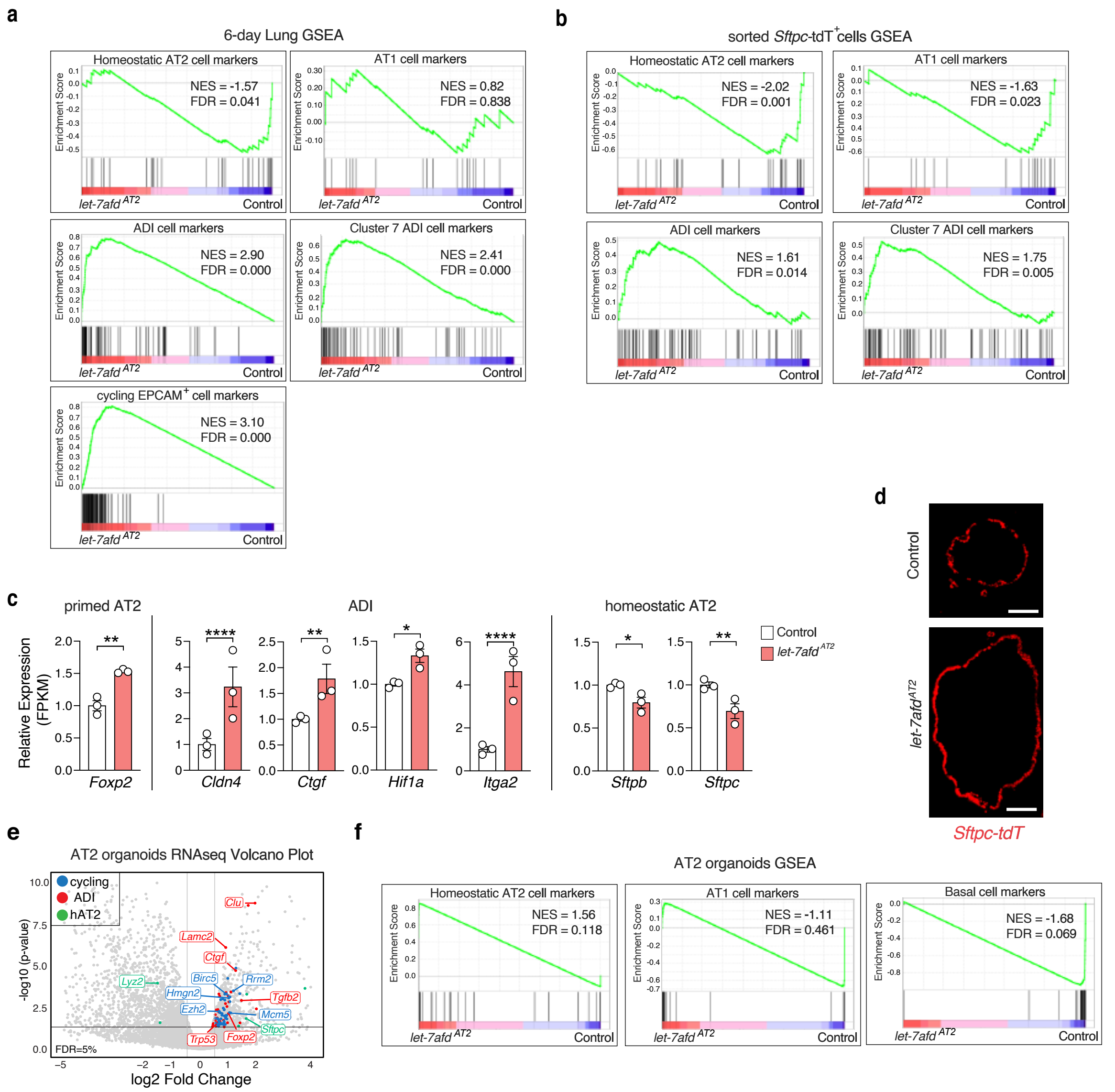


**Supplementary Fig 2** Transcriptomic analyses results from lungs and sorted AT2 cells of *let-7afd*$^{AT2}$ mice. **a** GSEA plots obtained by RNA-seq indicate induction of ADI and cycling EPCAM$^{+}$ cell gene sets but repression of hAT2 markers in lungs of *let-7afd*$^{AT2}$ mice compared to control mice following 6-days of iTAM (n = 3 mice per group). **b** GSEA plots indicate downregulation of hAT2 and AT1 cell markers and induction of ADI genes in *let-7afd*$^{-/-}$ vs control flow sorted *Sftpc*-tdT$^{+}$ cells following 6-days of iTAM (n = 3 samples per group from pools of 3 mice for RNA-seq). **c** Gene transcript levels detected by RNA-seq in *let-7afd*$^{-/-}$ vs control flow sorted *Sftpc*-tdT$^{+}$ cells. (n = 3 samples per group). Data are mean±s.e.m. ****p < 0.0001,**p < 0.01, *p < 0.05 by adjusted p value. **d** Representative IF images show the larger diameter of a *let-7afd*$^{-/-}$ compared to control AT2 alveolosphere on day 14 in AMM media. *Sftpc*-tdT (red); Scale bar 50μm. **e** Volcano plot of differentially expressed genes detected by RNA-seq in cultured *let-7afd*$^{-/-}$ vs control organoids highlights the induction of ADI (red), and cycling EPCAM$^{+}$ (blue) markers in AMM culture media (n = 2 mice per group). **f** GSEA plots derived by RNA-seq show subsignifant changes in hAT2, AT1 markers and lack of induction of basal cell markers in *let-7afd*$^{-/-}$ organoids in comparison to controls under AMM culture conditions.
