## Supplementary Fig. 3 for "*Let-7* restrains an oncogenic circuit in AT2 cells to prevent fibrogenic cell intermediates in pulmonary fibrosis"

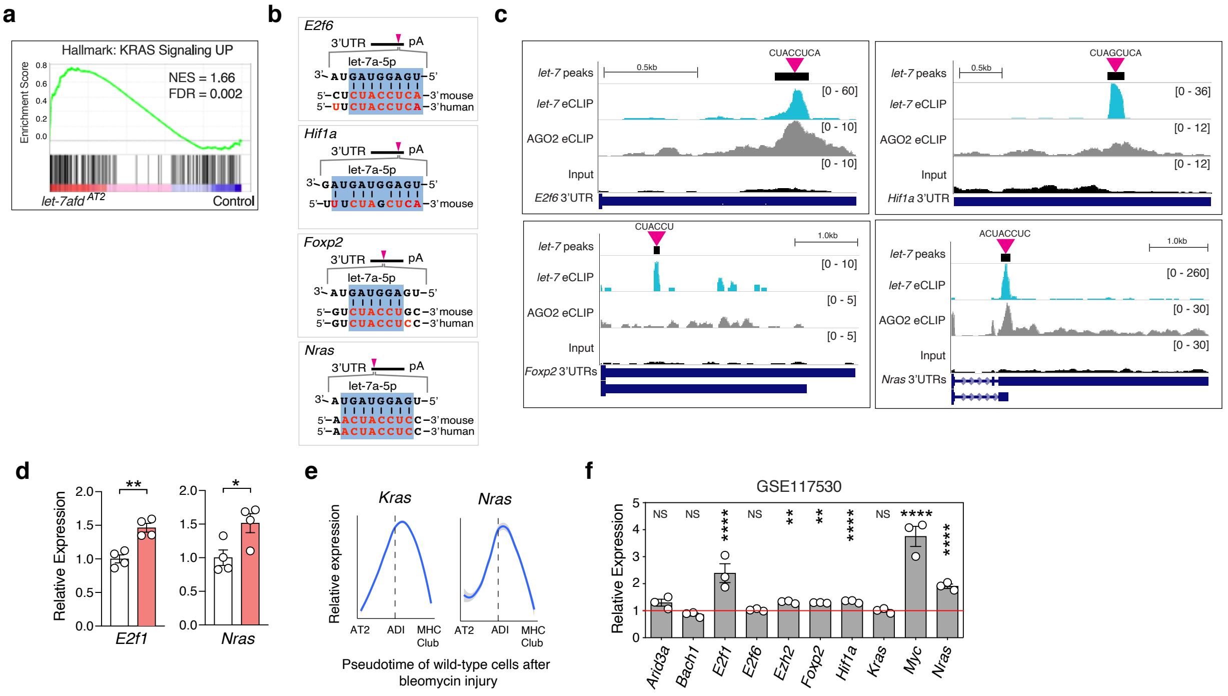


**Supplementary Fig 3** Characterization of selected *let-7* targets by AGO2-eCLIP. **a** GSEA plot obtained from RNA-seq shows induction of KRAS pathway genes in *let-7afd*$^{-/-}$ vs control *Sftpc*-tdT$^{+}$ cells after 6 days of iTAM (n = 3 samples per group from pools of 3 mice). **b** Schematic alignments of *let-7* "seed" region with target mRNA sequences in mice and humans. **c** The UCSC browser tracks for AGO2-eCLIP +let-7 shows the binding of *let-7* to the 3'UTRs of *E2f6*, *Foxp2*, *Hif1a*, and *Nras* respectively (n = 2 mice treated with bleomycin). Purple triangles indicate *let-7* binding motifs. **d** qPCR detection of indicated *let-7* targets from flow sorted *Sftpc*-tdT$^{+}$ cells on day 14 day after iTAM (n = 4 per group; from pools of 2 mice). Data are mean±s.e.m. **$p < 0.01$,*$p < 0.05$ by unpaired Student's t test. **e** Line plots show smoothed *Kras* and *Nras* relative expression levels across the ADI pseudotime trajectories after bleomycin injury (GSE141259). Dashed line corresponds to peak *Krt8* expression. **f** *Let-7* hub gene expression was determined from published RNA-seq (GSE117530) from flow sorted wild-type AT2 cells after 5-days of bleomycin-induced lung injury (n = 3 mice per group). Data are mean±s.e.m plotted relative to control samples = 1.0. **adjusted $p < 0.01$, ****adjusted $p < 0.0001$. Not significant (NS).
