## Supplementary Fig. 4 for "*Let-7* restrains an oncogenic circuit in AT2 cells to prevent fibrogenic cell intermediates in pulmonary fibrosis"

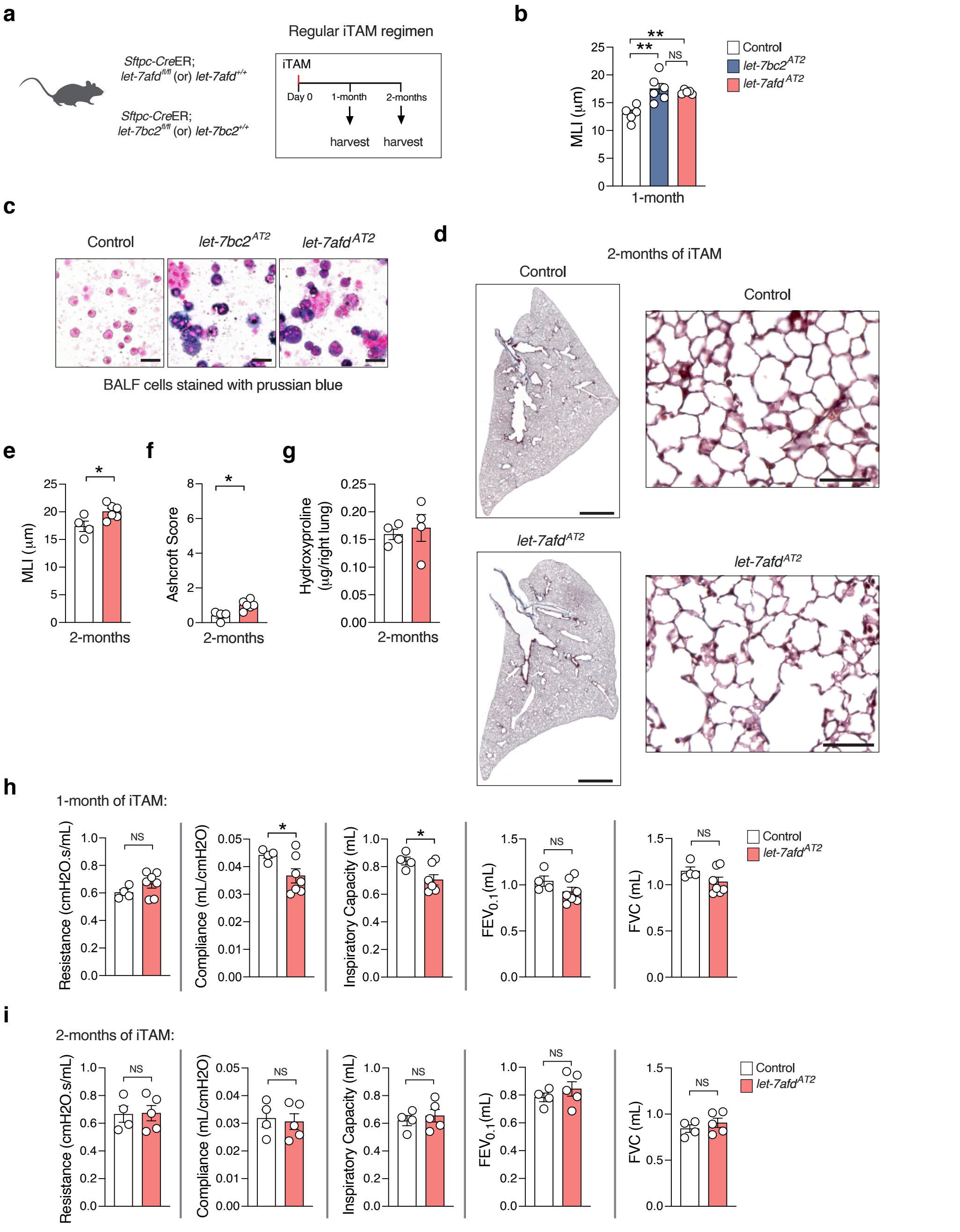


**Supplementary Fig 4** Phenotypic analysis of *let-7afd*$^{AT2}$ mice at 1-month and 2-months of iTAM. **a** Schematic representation for phenotypic analysis of mice at the 1- or 2-months time point after iTAM. **b** Mean linear intercept (MLI) measurements of lung morphometry from control (n = 5), *let-7bc2*$^{AT2}$ (n = 6), and *let-7afd*$^{AT2}$ (n = 5) mice harvested 1 month after iTAM. Data are mean±s.e.m. **p < 0.01, by one way ANOVA with Tukey's correction. **c** Representative brightfield images show prussian blue positive bronchoalveolar lavage fluid (BALF)-derived leukocytes in *let-7bc2*$^{AT2}$ and *let-7afd*$^{AT2}$ mice compared to controls after 1 month of iTAM (n = 6 mice per group). Scale bar: 50µm. **d** Representative Masson's trichrome-stained sections of mice lung lobes following 2-months of iTAM. Scale bars: 2 mm left panels; 50 µm right panels. **e,f** Mean linear intercept (MLI) and Ashcroft measurements of lung morphometry from control (n = 4) and *let-7afd*$^{AT2}$ (n = 6) mice at 2-months post-iTAM. Data are mean ±s.e.m. *p < 0.05, by unpaired Student's t test, (**e**); or Mann-Whitney U test, (**f**). **g** Hydroxyproline levels from lungs of indicated mice after 2-months of single iTAM regimen (n = 4 mice per group). Data are mean±s.e.m. Not significant (NS) by unpaired Student's t test. **h,i** Pulmonary biomechanics and spirometry quantitative measurements following 1-month (n = 4 and 7 mice per group) or 2-months (n = 4-5 mice per group) iTAM. Data are mean±s.e.m. *p < 0.05, by unpaired t test with Welch's correction. Not significant (NS). Each circle represents a mouse. Data is representative of two (**d,e,f,g,i**) or three (**b,c,h**) independent experiments.
