## Supplementary Fig. 5 for "*Let-7* restrains an oncogenic circuit in AT2 cells to prevent fibrogenic cell intermediates in pulmonary fibrosis"

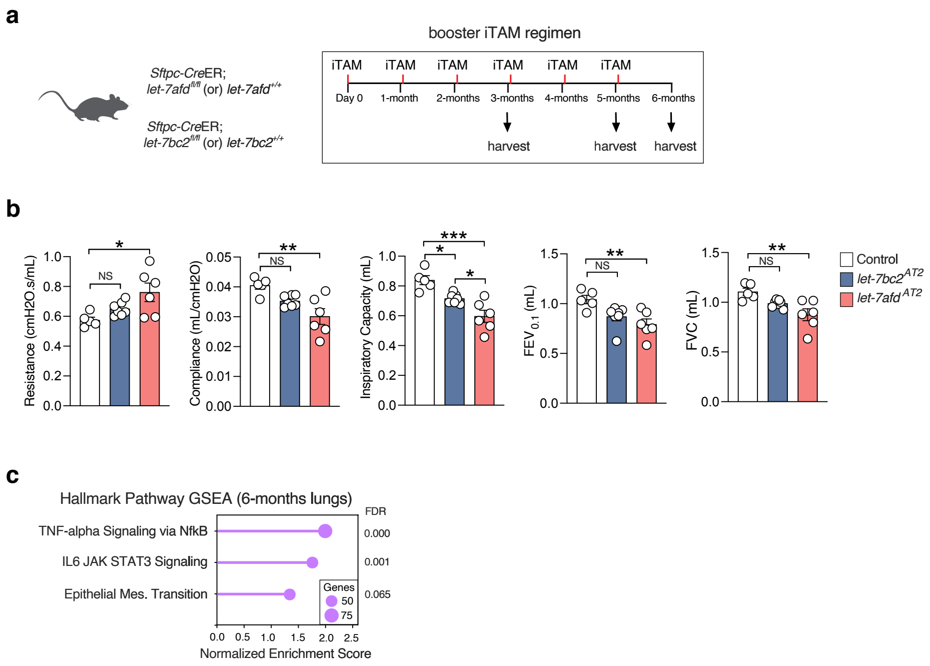


**Supplementary Fig 5** Phenotypic analysis of *let-7afd*$^{AT2}$ mice with booster iTAM regimen. **a** Schematic representation and strategy for booster iTAM regimen of mice. Mice were dosed once a month with iTAM and harvested 1-month following the last dose. **b** Pulmonary biomechanics and spirometry quantitative measurements following 3-months of booster iTAM (n = 5, 6, and 7 mice per group). Data are mean±s.e.m. *p < 0.05, **p < 0.01, ***p < 0.005 by one way ANOVA with Tukey's correction. Not significant (NS). **h** GSEA plots of bulk RNA-seq carried out from lungs of *let-7afd*$^{AT2}$ or control mice at 6 months of booster iTAM (n = 4 mice per group). Normalized enrichment scores and adjusted p values of for each gene set are indicated.
