## Supplementary Fig. 6 for "*Let-7* restrains an oncogenic circuit in AT2 cells to prevent fibrogenic cell intermediates in pulmonary fibrosis"

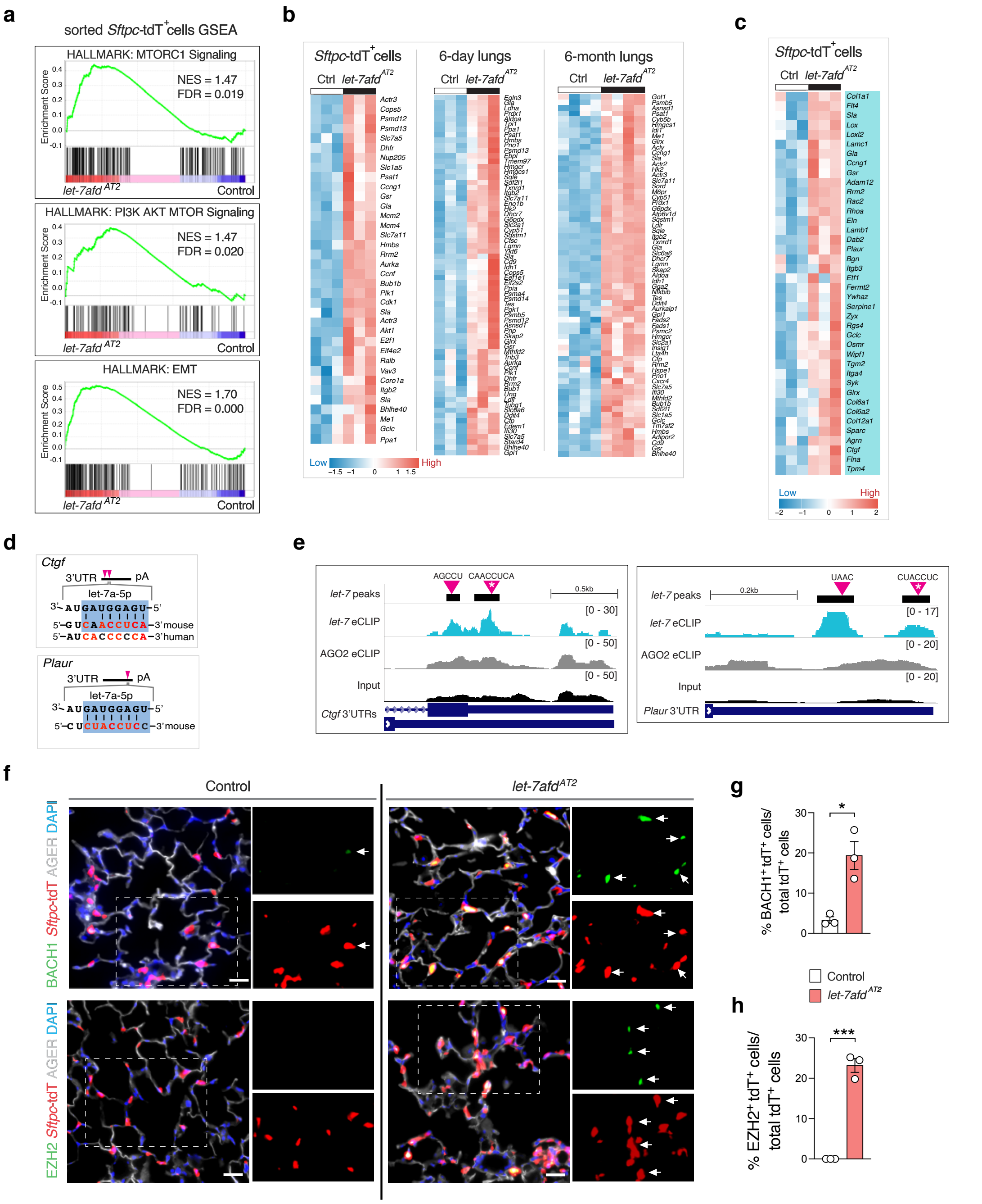

**Supplementary Fig 6** Delineation of targets of *let-7* associated with PI3K/AKT/MTOR. **a,b** The GSEA plots and heatmaps show selected PI3K/AKT/MTOR and EMT pathway genes which are upregulated upon loss of *let-7afd* in flow sorted AT2 cells or lungs. Adjusted p value < 0.05 vs control by RNA-seq. **c** RNA-seq based heat map of induced *let-7* targetome genes associated with PI3K/AKT/MTOR and/or EMT in purified AT2 cells after 6-days of iTAM. **d** Schematic sequence alignments of *let-7* "seed" region in *Ctgf* and *Plaur* mRNAs. **e** UCSC genome browser track indicates binding of *let-7* to the 3'UTRs of *Ctgf* and *Plaur*. Purple triangles are binding motifs of *let-7* and white asterisks are the chosen alignments in panel (**d**). **f** Representative IF images show increased numbers of BACH1⁺*Sftpc*-tdT⁺ (top panels) and EZH2⁺*Sftpc*-tdT⁺ (lower panels) cells (indicated by white arrows) in lungs of *let-7afd*$^{AT2}$ mice compared to control mice lungs after 5-months of booster iTAM. BACH1 (green); EZH2 (green); *Sftpc*-tdT (red); AGER (gray); DAPI (blue). Scale bars: 25 µm. **g**,**h** Quantification of BACH1⁺*Sftpc*-tdT⁺ cells (**h**) or EZH2⁺*Sftpc*-tdT⁺ cells (**i**) in total *Sftpc*-tdT⁺ cells (n = 3 mice per group). Data are mean±s.e.m. ***p < 0.001, *p < 0.05 by unpaired Student's t test.
