## Supplementary Fig. 7 for "*Let-7* restrains an oncogenic circuit in AT2 cells to prevent fibrogenic cell intermediates in pulmonary fibrosis"

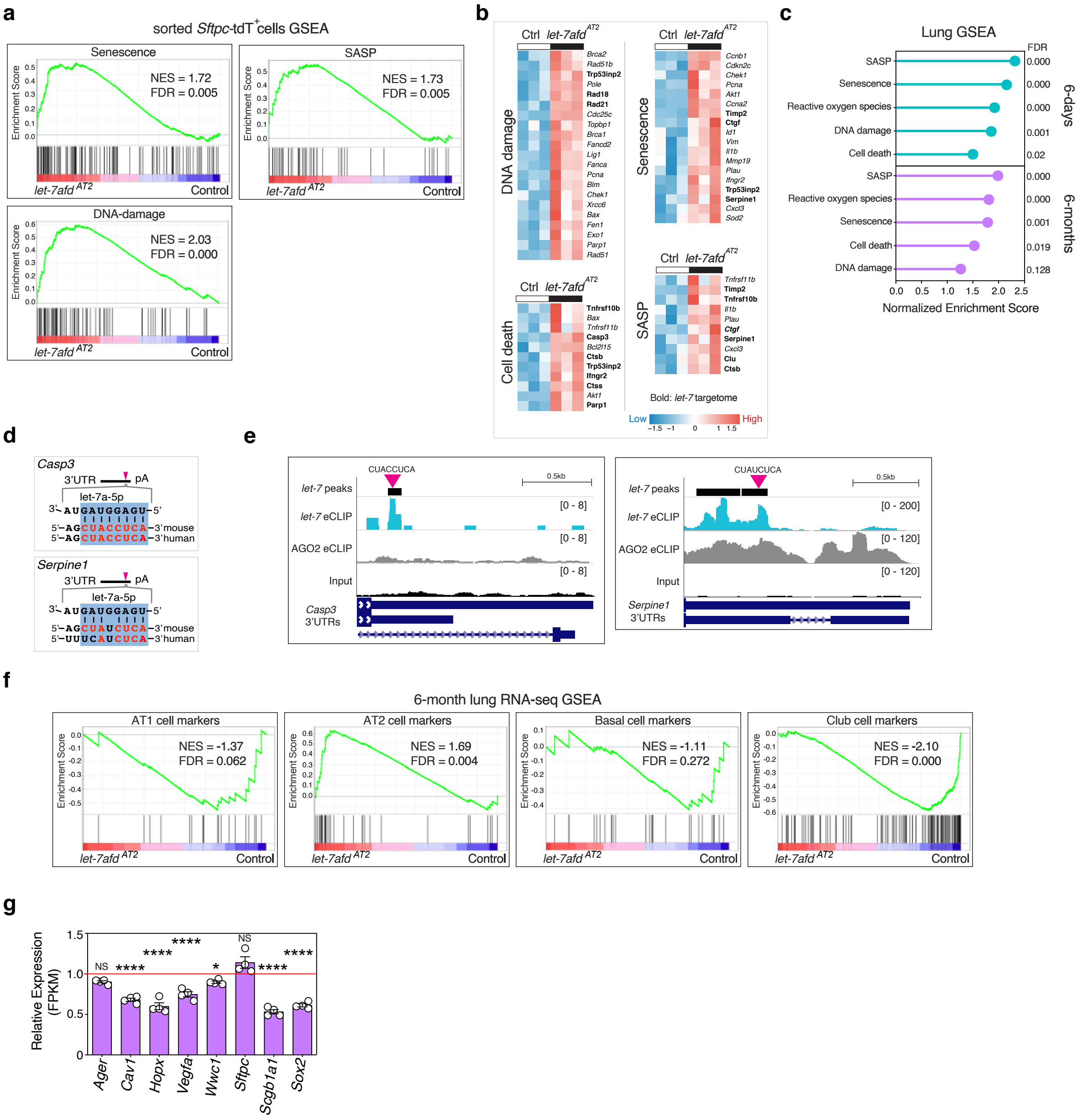


**Supplementary Fig 7** Delineation of *let-7* targets associated with cell death, DNA damage, and/or senescence. **a,b** GSEA (**a**) and heat maps (**b**) obtained by RNA-seq show induction of senescence, SASP, and DNA repair pathway genes upon deletion of *let-7afd* in purified AT2 cells after 6-days of iTAM (n = 3 samples per group from pools of 3 mice). **c** GSEA plots derived by RNA-seq analysis shows induction of indicated pathways in lungs of *let-7afd*$^{AT2}$ mice compared to control mice after 6-days of iTAM (n = 3 mice per group) or 6-months (n = 4 mice per group) of booster iTAM. **d** Schematic sequence alignments of *let-7* binding motifs in the *Casp3* and *Serpine* 3'UTRs. **e** AGO2-eCLIP UCSC genome browser tracks show binding of *let-7* to *Casp3* and *Serpine*. **f** GSEA of epithelial cell markers in whole lungs of *let-7afd*$^{AT2}$ compared to control mice with 6-months of booster induced iTAM deletion (n = 4 mice per group). **g** Selected AT1, AT2, club, and basal cell markers were detected by RNA-seq after 6-months of booster iTAM in *let-7afd*$^{AT2}$ compared to controls (n = 4 mice per group). Expression of *let-7afd*$^{AT2}$ samples were plotted relative to control sample average = 1.0. Data are mean±s.e.m. *adjusted $p < 0.05$, ****adjusted $p < 0.0001$. Not significant (NS).
