## Supplementary Fig. 8 for "*Let-7* restrains an oncogenic circuit in AT2 cells to prevent fibrogenic cell intermediates in pulmonary fibrosis"

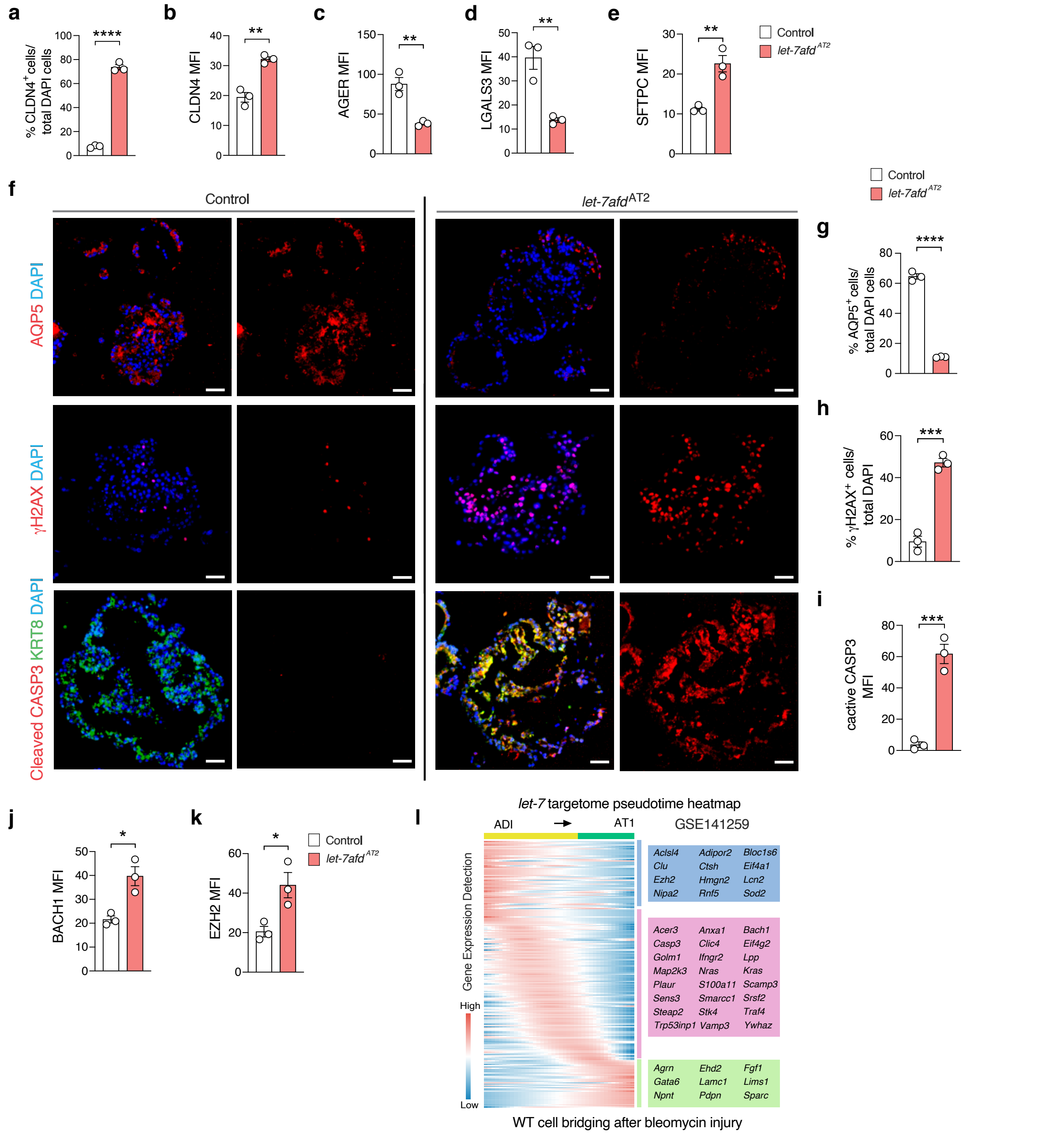


**Supplementary Fig 8** Requirement of *let-7afd* for effective AT2 to AT1 differentiation *ex vivo*. **a** Quantification of CLDN4$^+$ cells in total DAPI cells in *let-7afd*$^{-/-}$ vs control AT2 organoids. **b-e**,**j-k** Quantification of mean fluorescence intensity (MFI) for CLDN4 (**b**), AGER (**c**), LGALS3 (**d**), SFTPC (**e**), BACH1 (**j**) and EZH2 (**k**) in *let-7afd*$^{-/-}$ vs control AT2 organoids. **f** Representative immunostaining of AQP5$^+$ cells (upper panels), γH2AX$^+$ cells (middle panels) or active CASP3$^+$ cells (lower panels) in *let-7afd*$^{-/-}$ vs control alveolar organoids. Upper panels: AQP5 (red), DAPI (blue); middle panels γH2AX (red), DAPI (blue); bottom panels are active CASP3 (red), KRT8 (green), DAPI (blue). Scale bars: 25 μm. **g-i** Quantification of AQP5$^+$ cells in total DAPI cells (**g**), γH2AX$^+$ cells in total DAPI cells, (**h**) or active CASP3$^+$ MFI expression in total DAPI cells, (**i**). **a-k** AT2 organoids were cultured in AMM for 10 days followed by culture in ADM media for 7 days (n = 3 mice per group). Data are mean±s.e.m. ****$p < 0.0001$, ***$p < 0.001$, **$p < 0.01$, *$p < 0.05$, by unpaired Student's t test. Each dot represents a single mouse. Representative of three independent experiments. **l** The expression levels of 269 of 394 *let-7* targetome genes across the ADI to AT1 pseudotime trajectory is based on inferred likelihood of detection from scRNA-seq GSE141259 dataset.
