## Supplementary Fig. 9 for "*Let-7* restrains an oncogenic circuit in AT2 cells to prevent fibrogenic cell intermediates in pulmonary fibrosis"

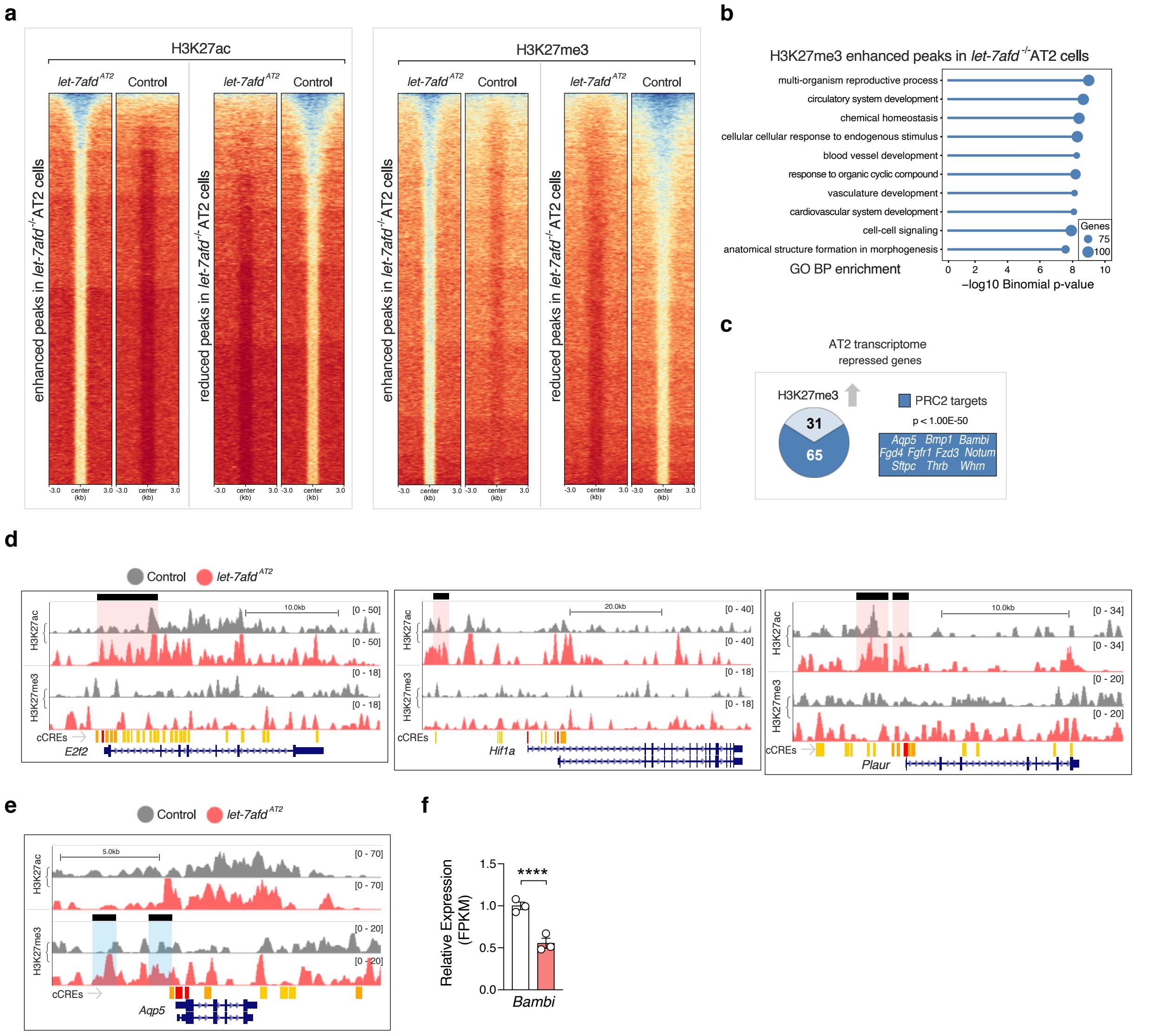

**Supplementary Fig 9** CUT&RUN data analyses. **a** CUT&RUN heatmaps of purified AT2 nuclei from *let-7afd*$^{-/-}$ vs control *Sftpc*-tdT$^{+}$ cells for H3K27ac and H3K27me3, grouped by differentiation enrichment between groups (3.0kb flanking peak center). **b** GO biological pathway (BP) enrichment of induced H3K27me3 peaks in *let-7afd*$^{-/-}$ compared to control *Sftpc*-tdT$^{+}$ cells. **c** Integration analysis of CUT&RUN with AT2 cell RNA-seq shows genes with congruent H3K27me3 enrichment and transcriptional repression in *let-7afd*$^{-/-}$ compared to control *Sftpc*-tdT$^{+}$ cells. ORA gene enrichment analysis shows that a significant fraction of downregulated genes are putative targets of the PRC2 complex. **d,e** UCSC tracks show H3K27ac and H3K27me3 peaks in selected genes in *let-7afd*$^{-/-}$ vs control *Sftpc*-tdT$^{+}$ cells. Induced H3K27ac peaks in *let-7afd*$^{-/-}$ *Sftpc*-tdT$^{+}$ cells are shaded in red. The induced H3K27me peaks are shaded in blue. ENCODE candidate cis-regulatory elements (cCREs). **f** *Bambi* transcript levels detected by RNA-seq in *let-7afd*$^{-/-}$ vs control flow sorted *Sftpc*-tdT$^{+}$ cells. (n = 3 samples per group). Data are mean±s.e.m. ****$p < 0.0001$, by adjusted p value.
