## Supplementary Fig. 10 for "*Let-7* restrains an oncogenic circuit in AT2 cells to prevent fibrogenic cell intermediates in pulmonary fibrosis"

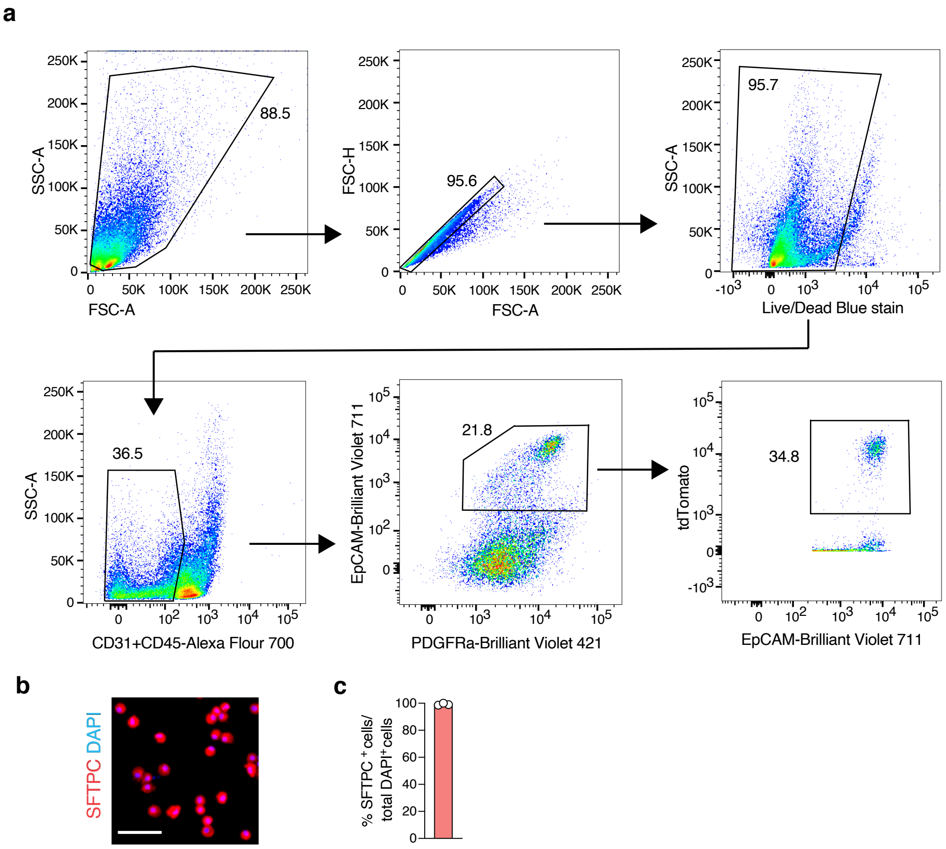


**Supplementary Fig 10** AT2 cell sorting strategy**. a** Representative flow cytometric gating and quantification strategy for isolation of lung Sftpc-tdT+ traced AT2 cells is shown. **b** Representative immunostaining of sorted cells with SFTPC (red) and DAPI (blue). Scale bars: 25 μm. **c** Quantification of SFTPC+ in total DAPI+ cells (n = 3 mice). Data are mean ±s.e.m.
