## Supplementary Data 1 for "*Let-7* restrains an oncogenic circuit in AT2 cells to prevent fibrogenic cell intermediates in pulmonary fibrosis"

### Supplementary Data 1: AG02-eCLIP+let-7 Results

| Gene | Peak Start | Peak End | miRNA | Chimeric reads |
|---|---|---|---|---|
| 0610030E20Rik | 72,352,688 | 72,352,789 | mmu-let-7a-5p | 2.5 |

AGO2-eCLIP+let-7 data has been deposited in the NCBI Gene Expression Omnibus data base.
