## Supplementary Data 2 for "*Let-7* restrains an oncogenic circuit in AT2 cells to prevent fibrogenic cell intermediates in pulmonary fibrosis"

Supplementary Data 2: *let-7* Targetome from fresh sorted *Sftpc*-tdT+ cells

| Symbol | AGO2-miR-eCLIP+let-7 | miRTarbase Experimentally Validated | TargetScan Predicted | Category | Class | Family | Base mean expression | Log Fold Change | p value | Adjusted p value |
|---|---|---|---|---|---|---|---|---|---|---|
| *Abcg1* | let-7a/b/c/d/f/i/k | | Targetscan | Co-nodes | Transporters and tra | ATP binding cassette | 206.239485 | 0.82 | 7.5E-04 | 1.2E-02 |
| *Abhd17c* | let-7a/b/c/k | miRTarbase | Targetscan | Enzymes | Esterases | Abhydrolases domain containing | 2329.9374 | 0.47 | 3.4E-03 | 3.5E-02 |
| *Acer3* | let-7a/b/c/d/f/k | | Targetscan | Enzymes | Ceramidases | Alkaline ceramidases (ACER) | 2953.81658 | 0.35 | 1.7E-03 | 2.2E-02 |
| *Acsl4* | let-7b/f/k | | | Enzymes | Long chain fatty aci | Long-chain acyl-CoA synthetases | 61437.3779 | 0.34 | 2.5E-03 | 2.9E-02 |
| *Acvr1b* | let-7a/b/c/d/e/f/k | miRTarbase | Targetscan | Receptors | Catalytic receptors | Transforming growth factor-? re | 3256.68587 | 0.61 | 1.6E-05 | 5.7E-04 |
| *Adam12* | let-7a/b/c/d/k | | Targetscan | Enzymes | Peptidases | ADAM metallopeptidase domain | 129.660315 | 2.70 | 4.3E-16 | 3.0E-13 |
| *Agrn* | let-7a/b/c/f | | | Receptors | Ligands | Agrin | 7272.1526 | 0.35 | 4.6E-03 | 4.4E-02 |
| *Ak2* | let-7a/d | | | Enzymes | Kinases | Adenylate kinases (AK) | 2582.08348 | 0.41 | 5.1E-04 | 8.7E-03 |
| *Alox5ap* | let-7b/c/k | | Targetscan | Enzymes | Enzyme regulators | Arachidonate 5-lipoxygenase act | 145.062122 | 1.62 | 1.8E-05 | 6.4E-04 |
| *Angptl4* | let-7a/b/c/d/f | | | Receptors | Ligands | Angiopoietin like | 510.858231 | 0.80 | 3.0E-03 | 3.3E-02 |
| *Anxa1* | let-7a/b/c/d/e/f/g | | | Receptors | Ligands | Annexin | 6471.44184 | 0.88 | 1.7E-05 | 6.0E-04 |
| *Ap1m1* | let-7a/b/k | | | Co-nodes | Adaptor, docking an | Adaptor related protein complex | 935.403915 | 0.45 | 8.1E-04 | 1.3E-02 |
| *Ap1s1* | let-7a/b/c/d/f/k | miRTarbase | Targetscan | Co-nodes | Adaptor, docking an | Adaptor related protein complex | 2857.42494 | 0.38 | 4.2E-04 | 7.6E-03 |
| *Arhgdib* | let-7a/b/c/f | | | Enzymes | Enzyme regulators | Rho GDP-dissociation inhibitors | 210.067991 | 1.25 | 4.9E-06 | 2.2E-04 |
| *Arid3a* | let-7a/b/c/d/e/f/k | miRTarbase | Targetscan | Transcription | ARID domain | ARID3 family | 660.398048 | 0.49 | 4.1E-03 | 4.1E-02 |
| *Arl5a* | let-7a/b/c/d/e/f/i | | Targetscan | Enzymes | GTPases | ADP-ribosylation factor like GTPa | 4077.31258 | 0.49 | 4.3E-05 | 1.3E-03 |
| *Asap1* | let-7d/f | | Targetscan | Enzymes | Enzyme regulators | ArfGAP with SH3 domain, ankyri | 351.038779 | 0.61 | 1.5E-03 | 1.9E-02 |
| *Aspm* | | miRTarbase | | Co-nodes | Cytoskeleton compo | Assembly factor for spindle micr | 1539.03295 | 0.74 | 1.4E-07 | 1.0E-05 |
| *Aurkb* | | miRTarbase | | Enzymes | Kinases | Aurora-related kinases (AURK) | 752.144142 | 0.85 | 2.5E-08 | 2.3E-06 |
| *Bend4* | | miRTarbase | Targetscan | Co-nodes | Other co-nodes | BEN domain containing | 245.889687 | 1.17 | 1.8E-06 | 9.5E-05 |
| *Bzw1* | let-7a/b/c/d/f/k | miRTarbase | Targetscan | Co-nodes | Leucine zipper | Basic leucine zipper and W2 dom | 8007.27718 | 0.45 | 1.0E-05 | 4.0E-04 |
| *Capn6* | let-7a/b/c/d/e/f/k | | | Enzymes | Peptidases | Calpains (CAPN) | 600.574487 | 0.92 | 1.1E-04 | 2.6E-03 |
| *Casp3* | let-7a/b/c | miRTarbase | Targetscan | Enzymes | Peptidases | Caspases (CASP) | 1614.54043 | 0.54 | 2.3E-06 | 1.2E-04 |
| *Ccdc25* | let-7a/b/c/d/f/k | | Targetscan | Co-nodes | Coiled coil domain | Coiled-coil domain containing | 2104.94891 | 0.38 | 1.3E-03 | 1.8E-02 |
| *Ccnb2* | | miRTarbase | | Enzymes | Enzyme regulators | Cyclins (CCN) | 1032.08187 | 0.98 | 1.3E-14 | 6.3E-12 |
| *Ccng1* | let-7a/b/c/d/e/f/k | miRTarbase | Targetscan | Enzymes | Enzyme regulators | Cyclins (CCN) | 6208.10145 | 0.62 | 8.8E-05 | 2.2E-03 |
| *Cd276* | let-7a/b/c/d/f | miRTarbase | Targetscan | Co-nodes | Cell surface proteins | Cluster of differentiation | 1709.13072 | 0.48 | 1.5E-04 | 3.4E-03 |
| *Cd38* | let-7a/b/c/d | | | Co-nodes | Cell surface proteins | Cluster of differentiation | 76.070217 | 1.49 | 2.8E-04 | 5.4E-03 |
| *Cd53* | let-7b/c/f | | | Co-nodes | Cell surface proteins | Cluster of differentiation | 170.274301 | 1.80 | 1.4E-06 | 7.5E-05 |
| *Cdc25B* | | miRTarbase | Targetscan | Enzymes | Phosphatases | Cell division cycle | 2960.92317 | 0.53 | 7.6E-07 | 4.5E-05 |
| *Cdc34* | let-7a/b/c/d/e/f/k | miRTarbase | Targetscan | Enzymes | Aminoacyltransfera | Cell division cycle | 1689.38422 | 0.65 | 3.1E-04 | 5.9E-03 |
| *Clcn5* | let-7a/b/c/d/f | | Targetscan | Ion channels | Other ion channels | Chloride voltage-gated channel | 1512.17297 | 0.50 | 4.4E-04 | 7.9E-03 |
| *Clu* | let-7a/b/c/d/e/f/k | | | Co-nodes | Other co-nodes | Clusterin | 17147.8768 | 0.75 | 1.6E-04 | 3.5E-03 |
| *Cnih1* | let-7a/b/c/d/e/f/k | | | Co-nodes | Receptor associated | Cornichon AMPA receptor auxilia | 2659.36676 | 0.35 | 1.8E-03 | 2.2E-02 |
| *Cntrl* | let-7a/b/c/d/e/f/k | miRTarbase | Targetscan | Co-nodes | Cell cycle, cell divisi | Centriolin | 4859.27342 | 0.76 | 1.0E-07 | 8.1E-06 |
| *Col12a1* | let-7a/b/c | | | Receptors | Ligands | Collagen chain | 2664.19228 | 0.58 | 2.2E-03 | 2.6E-02 |
| *Col18a1* | let-7c/e | | | Receptors | Ligands | Collagen chain | 1930.44299 | 0.58 | 1.9E-04 | 4.0E-03 |
| *Col5a2* | let-7a/b/c/d/e/f/k | | Targetscan | Receptors | Ligands | Collagen chain | 125.788593 | 1.31 | 6.6E-04 | 1.1E-02 |
| *Col6a1* | let-7a/b/c/d/f | | | Receptors | Ligands | Collagen chain | 27174.7027 | 0.50 | 5.6E-04 | 9.4E-03 |
| *Col6a2* | let-7a/b/c/d/f/k | | | Receptors | Ligands | Collagen chain | 6591.88884 | 0.75 | 5.9E-07 | 3.6E-05 |
| *Csf2rb* | let-7a/b/c/d/f/i/k | | Targetscan | Receptors | Catalytic receptors | GM-CSF/CSF2 receptor | 296.931183 | 1.58 | 4.9E-09 | 5.3E-07 |
| *Ctbs* | let-7a/b/c/d/e/f | | Targetscan | Co-nodes | Other co-nodes | Chitobiase | 752.763939 | 0.55 | 1.8E-03 | 2.3E-02 |
| *Ctgf* | let-7a/b/c/k | | | Receptors | Ligands | Cellular communication network | 621.083205 | 0.75 | 1.8E-04 | 3.9E-03 |
| *Ctsb* | let-7a/b/c/d/e/f/k | | Targetscan | Enzymes | Peptidases | Cathepsins (CTS) | 36832.4062 | 0.34 | 1.8E-03 | 2.2E-02 |
| *Ctss* | let-7a/b/d/k | | Targetscan | Enzymes | Peptidases | Cathepsins (CTS) | 506.598698 | 1.52 | 1.2E-09 | 1.7E-07 |
| *Dab2* | let-7a/b/c/d/f | | | Co-nodes | Adaptor, docking an | DAB adaptor protein | 179.881386 | 0.87 | 3.0E-04 | 5.8E-03 |
| *Dera* | let-7a/b/f/k | | Targetscan | Enzymes | Aldolases | Deoxyribose-phosphate aldolase | 832.474659 | 0.78 | 1.6E-07 | 1.2E-05 |
| *Dna2* | | miRTarbase | Targetscan | Enzymes | Helicases (DNA) | DNA2/NAM7 | 287.152318 | 0.67 | 2.6E-04 | 5.1E-03 |
| *Dot1l* | let-7b/d/k | | | Enzymes | Methyltransferases | Disruptor of telomeric silencing | 2448.37338 | 0.67 | 8.4E-06 | 3.4E-04 |
| *Dpp3* | let-7a/c/d | | Targetscan | Enzymes | Peptidases | Dipeptidyl-peptidases (DPP) | 2903.05874 | 0.43 | 2.3E-04 | 4.6E-03 |
| *E2f1* | | miRTarbase | | Transcription | E2F/FOX | E2F | 846.012936 | 0.50 | 6.2E-04 | 1.0E-02 |
| *Edn1* | let-7a/b/c/d/f/g/k | miRTarbase | Targetscan | Receptors | Ligands | Endothelin | 305.709657 | 1.32 | 3.3E-04 | 6.2E-03 |
| *Eif4a1* | let-7a/b/c/d/f/k | | | Enzymes | Helicases (RNA) | Eukaryotic initiation factors 4A (I | 19548.2914 | 0.43 | 5.6E-04 | 9.4E-03 |
| *Eif4e2* | let-7b | | | Co-nodes | Translation factors | Eukaryotic translation initiation f | 2226.81698 | 0.39 | 1.1E-03 | 1.6E-02 |
| *Elf4* | let-7a/b/c/d/f/i/k | | Targetscan | Transcription | Tryptophan cluster | Elf-1-like | 220.144854 | 0.68 | 1.9E-03 | 2.3E-02 |
| *Eln* | let-7a/b/c/d/e/f/g/i/k | | | Co-nodes | Other co-nodes | Elastin | 341.426621 | 2.28 | 2.7E-11 | 5.6E-09 |
| *Emilin2* | | miRTarbase | | Co-nodes | Other co-nodes | Elastin microfibril interfacer | 156.043914 | 1.50 | 7.5E-07 | 4.4E-05 |
| *Epb41l2* | let-7a/b/c/f | | | Co-nodes | Membrane proteins | Erythrocyte membrane protein b | 669.517623 | 0.54 | 4.8E-04 | 8.4E-03 |
| *Epha4* | | miRTarbase | Targetscan | Receptors | Catalytic receptors | Ephrin receptors | 948.679926 | 1.66 | 2.0E-16 | 1.7E-13 |
| *Espl1* | | miRTarbase | Targetscan | Enzymes | Peptidases | Extra spindle pole bodies like, se | 1087.30171 | 0.83 | 3.0E-09 | 3.6E-07 |
| *Ezh2* | | miRTarbase | | Co-nodes | Polycomb group (Pc | Enhancer of zeste polycomb repr | 1284.86944 | 0.60 | 1.8E-05 | 6.3E-04 |
| *Faf1* | let-7a/b/c | | | Co-nodes | Apoptosis and apop | Fas associated factor | 2561.89098 | 0.37 | 8.4E-04 | 1.3E-02 |
| *Fcer1g* | let-7c/d/e/f | | | Receptors | Other receptors | Fc epsilon receptors | 99.1166454 | 1.50 | 1.8E-06 | 9.5E-05 |
| *Fermt2* | let-7a/b/c/d/f/i | | | Co-nodes | Cytoskeleton compo | Fermitin | 9435.05186 | 0.57 | 1.4E-04 | 3.2E-03 |
| *Fermt3* | let-7b/k | | | Co-nodes | Cytoskeleton compo | Fermitin | 78.1377065 | 2.44 | 5.4E-07 | 3.3E-05 |

| | | | | | | | | | | |
|---|---|---|---|---|---|---|---|---|---|---|
| *Fgr* | let-7a/b/c/d/e/f/k | | Targetscan | Enzymes | Kinases | Src kinases | 94.2283061 | 1.67 | 1.2E-04 | 2.9E-03 |
| *Flna* | let-7a/b/k | | | Co-nodes | Cytoskeleton compo | Filamin | 22266.0562 | 0.48 | 1.6E-05 | 5.9E-04 |
| *Flt4* | let-7a/c/f | | | Receptors | Catalytic receptors | VEGF receptors | 104.164119 | 1.54 | 2.5E-04 | 4.9E-03 |
| *Fmnl3* | | miRTarbase | | Co-nodes | Cytoskeleton compo | Formin like | 582.456874 | 0.62 | 7.6E-05 | 2.0E-03 |
| *Foxp2* | let-7b/c/d/f | | Targetscan | Transcription | E2F/FOX | FOXP | 1242.01386 | 0.53 | 3.3E-05 | 1.0E-03 |
| *Galns* | let-7a/b/c/f | | Targetscan | Enzymes | Other enzymes | Galactosamine (N-acetyl)-6-sulf | 578.640794 | 0.52 | 4.3E-04 | 7.7E-03 |
| *Gclc* | let-7a/b/c/d/f | | | Enzymes | Other enzymes | Glutamate-cysteine ligase cataly | 47271.5231 | 0.61 | 6.8E-04 | 1.1E-02 |
| *Ggta1* | let-7a/b/c/k | | Targetscan | Enzymes | Glycosyltransferase | Glycoprotein galactosyltransfera | 78.2159546 | 1.42 | 2.9E-04 | 5.6E-03 |
| *Gla* | let-7a/b/c/f | | Targetscan | Enzymes | Glycosidases | Galactosidase alpha | 757.221906 | 0.53 | 8.9E-04 | 1.3E-02 |
| *Gnptab* | let-7a/b/c/d/e/f | | Targetscan | Enzymes | Transferases | N-acetylglucosamine-1-phospha | 14815.639 | 0.61 | 2.9E-06 | 1.4E-04 |
| *Golm1* | let-7a/b/c/d/f | | Targetscan | Co-nodes | Endosomal, lysosom | Golgi membrane protein | 6403.19967 | 0.76 | 4.2E-12 | 1.1E-09 |
| *Gsr* | let-7f/k | | | Enzymes | Oxidoreductases | Glutathione-disulfide reductases | 3120.18544 | 0.56 | 4.6E-04 | 8.2E-03 |
| *Guf1* | let-7a/b/c/d | | Targetscan | Enzymes | GTPases | GUF1 homolog, GTPases | 390.930901 | 0.56 | 1.9E-03 | 2.4E-02 |
| *Haspin* | | miRTarbase | | Enzymes | Kinases | Histone H3 associated protein ki | 245.656677 | 0.78 | 1.2E-04 | 2.9E-03 |
| *Hif1a* | let-7a/b/c/d/e/f/i | | | Transcription | BHLH factors | Ahr-like family | 7681.10229 | 0.33 | 1.9E-03 | 2.4E-02 |
| *Hist1H1D* | | miRTarbase | | Co-nodes | Histones and histon | H1 linker histone | 15.3176421 | 3.20 | 4.9E-04 | 8.4E-03 |
| *Hmgn2* | | miRTarbase | Targetscan | Co-nodes | Transcriptional core | High mobility group nucleosoma | 6415.32359 | 0.60 | 7.5E-09 | 7.8E-07 |
| *Hspb6* | let-7a/b/c/d/f/k | | | Co-nodes | Stress response fact | Heat shock protein family B (sma | 540.481263 | 0.65 | 5.9E-05 | 1.6E-03 |
| *Ifngr2* | let-7c/k | | | Receptors | Catalytic receptors | Interferon gamma receptor | 1612.347 | 0.61 | 9.2E-05 | 2.3E-03 |
| *Igf2bp2* | let-7a/b/c/d/f/i/k | | Targetscan | Co-nodes | RNA binding and RB | Insulin like growth factor mRNA | 450.159221 | 0.86 | 7.5E-06 | 3.1E-04 |
| *Il1rap* | let-7a/b/c | | | Receptors | Catalytic receptors | Interleukin 1 receptor accessory | 520.012627 | 0.63 | 3.7E-04 | 6.8E-03 |
| *Ints2* | let-7a/b/c/f/k | | Targetscan | Co-nodes | Transcriptional core | Integrator complex subunit | 905.063721 | 0.59 | 5.9E-05 | 1.6E-03 |
| *Ints6l* | let-7a/b/c/d/e/f/k | | | Co-nodes | Transcriptional core | Integrator complex subunit like | 1983.17411 | 0.52 | 6.9E-04 | 1.1E-02 |
| *Ipo9* | | miRTarbase | Targetscan | Co-nodes | Karyopherins | Importin | 3650.2905 | 0.30 | 1.6E-03 | 2.1E-02 |
| *Iqgap3* | let-7b/c/k | | | Enzymes | Enzyme regulators | IQ motif containing GTPase activ | 2387.36157 | 0.82 | 4.3E-09 | 4.8E-07 |
| *Isca2* | let-7a/b/c/d/f/k | | Targetscan | Co-nodes | Vitamin and trace e | Iron-sulfur cluster assembly | 428.7059 | 0.54 | 1.1E-03 | 1.6E-02 |
| *Itga4* | let-7a/b/c/d/f | | Targetscan | Receptors | Catalytic receptors | Integrins | 277.35357 | 1.46 | 1.6E-05 | 6.0E-04 |
| *Itgal* | let-7a/b/c/d/e/f/k | | Targetscan | Receptors | Catalytic receptors | Integrins | 521.689643 | 1.31 | 7.2E-06 | 3.0E-04 |
| *Kazn* | let-7b/c | | | Co-nodes | Interacting proteins | Kazrin, periplakin interacting pro | 300.930391 | 0.90 | 2.3E-04 | 4.6E-03 |
| *Knstrn* | let-7a/b | | | Co-nodes | Cell cycle, cell divisi | Kinetochore localized astrin (SPA | 1356.27459 | 1.10 | 3.0E-20 | 4.0E-17 |
| *Lamb1* | let-7b | | Targetscan | Receptors | Ligands | Laminin subunit | 273.806388 | 0.81 | 5.5E-04 | 9.3E-03 |
| *Lamc1* | let-7a/b/c/d/e/f/k | | | Receptors | Ligands | Laminin subunit | 8947.39691 | 0.38 | 1.8E-03 | 2.2E-02 |
| *Lcn2* | let-7b/c | | | Co-nodes | Transporters and tra | Lipocalin | 59346.9482 | 1.30 | 4.6E-11 | 9.1E-09 |
| *Lims1* | let-7b/c/f/k | | Targetscan | Co-nodes | Zinc finger proteins | LIM zinc finger domain containin | 3023.22051 | 0.39 | 4.8E-04 | 8.4E-03 |
| *Lipa* | let-7a/b/c/d/f/k | | | Enzymes | Lipases | Lipases (LIP) | 349.09635 | 1.88 | 1.6E-15 | 9.0E-13 |
| *Lox* | let-7a/b/c/e/f/g/k | | | Enzymes | Oxidases | Protein-lysine 6-oxidases (LOX) | 68.7293954 | 1.29 | 2.6E-03 | 2.9E-02 |
| *Loxl2* | let-7a/b/c/d/e/f/k | | | Enzymes | Oxidases | Protein-lysine 6-oxidases (LOX) | 48.1523269 | 1.36 | 2.6E-03 | 2.9E-02 |
| *Lpgat1* | let-7a/b/c/d/f/g/i | | Targetscan | Enzymes | Acyltransferases | Lysophosphatidylglycerol acyltra | 3007.81646 | 0.39 | 2.0E-03 | 2.4E-02 |
| *Lpp* | let-7a/b/c/d/e/f/k | | | Co-nodes | LIM domain protein | LIM domain containing preferred | 5955.26254 | 0.49 | 3.6E-03 | 3.7E-02 |
| *Lrg1* | let-7a/b/c/d/e/f/i/k | | | Co-nodes | Glycoproteins | Leucine rich alpha-2-glycoprotei | 39219.9018 | 0.74 | 1.4E-04 | 3.3E-03 |
| *Lsp1* | let-7b/c/d/e/k | | | Co-nodes | Immune system cor | Lymphocyte specific protein | 182.4613 | 1.26 | 4.2E-03 | 4.2E-02 |
| *Ly6c1* | let-7a/b/c/d/f/k | | | Co-nodes | Immune system cor | Lymphocyte antigen | 3919.53312 | 0.54 | 8.9E-04 | 1.3E-02 |
| *Man2a2* | let-7a/b/c/k | | | Enzymes | Glycosidases | Mannosidases (MAN) | 5032.41588 | 0.33 | 5.2E-04 | 8.8E-03 |
| *Marcks* | let-7a/b/c/f | | | Co-nodes | Substrate proteins | Myristoylated alanine rich protei | 237.193994 | 0.66 | 4.2E-03 | 4.2E-02 |
| *Mcam* | let-7b/c/f/k | | | Co-nodes | Adhesion molecules | Melanoma cell adhesion molecu | 315.436362 | 0.93 | 1.5E-03 | 2.0E-02 |
| *Mpeg1* | let-7a/b/c/d/e/k | | | Co-nodes | Other co-nodes | Macrophage expressed | 2423.61609 | 0.54 | 9.4E-04 | 1.4E-02 |
| *Mpzl1* | let-7b/c/d/f/k | | | Co-nodes | CNS proteins | Myelin protein zero like | 1160.90949 | 0.39 | 3.3E-03 | 3.4E-02 |
| *Mrps2* | | miRTarbase | | Co-nodes | Ribosomes and ribo | Mitochondrial ribosomal protein | 616.948406 | 0.45 | 4.7E-03 | 4.5E-02 |
| *Msn* | let-7a/b/c/d/f/k | | Targetscan | Co-nodes | Cytoskeleton compo | Moesin | 17743.2095 | 0.41 | 3.9E-04 | 7.2E-03 |
| *Msr1* | let-7a/b/c/d/f/k | | Targetscan | Receptors | Other receptors | Scavenger receptors | 29.4635855 | 2.63 | 1.2E-04 | 2.8E-03 |
| *Mtch2* | let-7a/b/c/d/f/k | | Targetscan | Co-nodes | Transporters and tra | Mitochondrial carrier | 5303.9993 | 0.30 | 3.0E-03 | 3.2E-02 |
| *Myc* | let-7a/b/c/d/f/k | miRTarbase | | Transcription | BHLH factors | Myc/Max factor | 933.797096 | 0.64 | 3.7E-01 | 6.6E-01 |
| *Myo1f* | let-7a/b/c/d/f/k | | Targetscan | Co-nodes | Cytoskeleton compo | Myosin | 167.281924 | 1.60 | 6.7E-06 | 2.8E-04 |
| *Nemp1* | let-7b/k | | | Co-nodes | Membrane proteins | Nuclear envelope integral memb | 572.532377 | 0.53 | 4.0E-03 | 4.0E-02 |
| *Nfam1* | let-7a/b/c/d/e | | Targetscan | Co-nodes | Other co-nodes | NFAT activating protein with ITA | 107.129008 | 1.78 | 5.7E-04 | 9.5E-03 |
| *Nhlrc3* | let-7a/b/c/d/f/k | miRTarbase | Targetscan | Co-nodes | Other co-nodes | NHL repeat containing E3 ubiqui | 697.461967 | 0.50 | 4.3E-03 | 4.2E-02 |
| *Nipa2* | let-7a/b/c/f | | Targetscan | Co-nodes | Transporters and tra | NIPA magnesium transporter | 1899.0818 | 0.40 | 1.6E-04 | 3.6E-03 |
| *Nras* | let-7a/b/c/d/e/f/g/k | miRTarbase | Targetscan | Enzymes | GTPases | Ras Type GTPases | 2226.94796 | 0.56 | 9.1E-06 | 3.6E-04 |
| *Nrp2* | let-7b | | Targetscan | Co-nodes | CNS Proteins | Neuropilin | 952.125907 | 0.49 | 1.5E-03 | 2.0E-02 |
| *Nudt4* | | miRTarbase | | Enzymes | Phosphatases | Nudix (NUDT) | 8162.74269 | 0.66 | 3.8E-09 | 4.3E-07 |
| *Olr1* | let-7a/b/c/d/f/k | | Targetscan | Receptors | Other receptors | Scavenger receptors | 37.6018089 | 2.58 | 2.9E-05 | 9.4E-04 |
| *Osmr* | let-7a/b/c/d/e/f/k | | Targetscan | Receptors | Catalytic receptors | Oncostatin M receptor | 5622.88061 | 0.56 | 1.4E-03 | 1.9E-02 |
| *Parp1* | let-7a/b/c/f/k | miRTarbase | Targetscan | Enzymes | ADP ribosyltransfera | Poly [ADP-ribose] polymerases (I | 4423.29318 | 0.31 | 2.4E-03 | 2.8E-02 |
| *Pcx* | let-7a/b/c/d/e/f/g/k | | Targetscan | Enzymes | Other enzymes | Pyruvate carboxylases | 18292.6018 | 1.08 | 1.0E-15 | 6.1E-13 |
| *Pdpn* | let-7b | | Targetscan | Co-nodes | Other co-nodes | Podoplanin | 837.750866 | 0.71 | 2.5E-06 | 1.2E-04 |
| *Picalm* | let-7a/b/c/e/f/k | | | Co-nodes | Endosomal, lysosom | Phosphatidylinositol binding clat | 10872.11 | 0.32 | 1.3E-03 | 1.8E-02 |
| *Pign* | | miRTarbase | | Co-nodes | Other co-nodes | Phosphatidylinositol glycan subu | 4002.05101 | 0.29 | 5.3E-03 | 4.9E-02 |
| *Pigs* | let-7a/c/d/e/f/g | | Targetscan | Co-nodes | Other co-nodes | Phosphatidylinositol glycan subu | 2733.63149 | 0.28 | 5.1E-03 | 4.7E-02 |
| *Pim1* | let-7a/b/c/d/e/f/k | | | Enzymes | Kinases | Pim-1 kinases (PIM) | 3300.96116 | 0.80 | 2.5E-04 | 5.0E-03 |
| *Plagl2* | let-7a/b/c/d/f/g/k | miRTarbase | Targetscan | Transcription | C2H2 Zn finger facto | PLAG Zinc Finger | 1827.19399 | 0.50 | 1.8E-03 | 2.2E-02 |
| *Plaur* | let-7a/b/c/d/e/f/g | | Targetscan | Receptors | Other receptors | Plasminogen activator, urokinase | 342.066655 | 1.54 | 3.1E-09 | 3.7E-07 |
| *Plxnb2* | let-7a/c/d/f/g | | | Receptors | Other receptors | Plexin | 37213.3947 | 0.39 | 3.1E-04 | 5.9E-03 |
| *Pmpca* | | miRTarbase | Targetscan | Enzymes | Peptidases | Mitochondrial processing peptida | 3120.90643 | 0.36 | 1.1E-03 | 1.6E-02 |
| *Pqlc2* | let-7b/c/d/f/g/k | | Targetscan | Co-nodes | Transporters and tra | Solute carrier superfamily memb | 611.240566 | 0.59 | 1.3E-03 | 1.7E-02 |
| *Prim2* | | miRTarbase | | Enzymes | Other enzymes | DNA primase subunit | 692.152659 | 0.42 | 3.7E-03 | 3.7E-02 |
| *Prr5l* | let-7a/b/c/d/f/g/k | | Targetscan | Co-nodes | Proline rich proteins | Proline rich like | 454.832409 | 0.85 | 9.2E-05 | 2.3E-03 |
| *Psmd1* | let-7b/k | | | Co-nodes | Proteasome | Proteasome 26S subunit, non-AT | 6494.46754 | 0.28 | 4.8E-03 | 4.5E-02 |

| | | | | | | | | | | |
|---|---|---|---|---|---|---|---|---|---|---|
| *Ptafr* | let-7a/b/c/d/e/f/k | | Targetscan | Receptors | G protein coupled re | Platelet-activating factor recepto | 168.364742 | 1.29 | 5.4E-03 | 4.9E-02 |
| *Rab8b* | let-7a/c/d/e/f | | Targetscan | Enzymes | GTPases | RAB, member RAS oncogene | 324.539683 | 0.98 | 5.8E-05 | 1.6E-03 |
| *Rac2* | let-7a/b/c/d/f/k | | | Enzymes | GTPases | Rho GTPases | 113.934109 | 1.25 | 5.3E-05 | 1.5E-03 |
| *Rad18* | | miRTarbase | Targetscan | Enzymes | E3 ubiquitin ligases | Cell cycle checkpoint proteins (R/ | 268.314306 | 0.69 | 4.6E-04 | 8.2E-03 |
| *Rad21* | let-7d | miRTarbase | | Co-nodes | Cohesin complex | Cell cycle checkpoint proteins (R/ | 10194.2133 | 0.29 | 1.2E-03 | 1.7E-02 |
| *Ralb* | let-7b | | Targetscan | Enzymes | GTPases | Ras Type GTPases | 1827.08035 | 0.41 | 1.1E-03 | 1.6E-02 |
| *Rbm3* | let-7c/f | | | Co-nodes | RNA binding and RB | RNA binding motif protein | 10499.2059 | 1.26 | 1.1E-17 | 1.2E-14 |
| *Rrm1* | | miRTarbase | Targetscan | Enzymes | Reductases | Ribonucleoside-diphosphate redu | 3854.88476 | 0.61 | 1.2E-06 | 6.8E-05 |
| *Rrm2* | | miRTarbase | Targetscan | Enzymes | Enzyme regulators | Ribonucleoside-diphosphate redu | 2607.10892 | 0.92 | 7.8E-15 | 3.9E-12 |
| *S100a11* | let-7a/b/c/d/e/f/k | | Targetscan | Co-nodes | Calcium associated | S100 calcium binding protein | 10863.0006 | 0.33 | 1.0E-03 | 1.5E-02 |
| *Sdc3* | let-7b/c/f/k | | | Co-nodes | Cell surface proteins | Syndecan | 240.832121 | 0.76 | 4.8E-04 | 8.4E-03 |
| *Serpina3n* | let-7a/b/c/d/e/f/g/i/k | | | Enzymes | Enzyme regulators | Serpin A | 124.185328 | 1.48 | 7.9E-05 | 2.0E-03 |
| *Serpine1* | let-7a/b/c/d/e/f/g/i/k | | | Receptors | Ligands | Serpins | 470.869772 | 0.83 | 4.9E-04 | 8.5E-03 |
| *Ska3* | | miRTarbase | | Co-nodes | Cell cycle, cell divisi | Spindle and kinetochore associat | 215.580403 | 0.82 | 2.1E-03 | 2.5E-02 |
| *Sla* | let-7b/c | | Targetscan | Co-nodes | SH2 domain contain | Src like adaptor | 90.359727 | 1.44 | 2.1E-04 | 4.3E-03 |
| *Slc16a6* | let-7b/c | | | Co-nodes | Transporters and tra | Solute carrier superfamily memb | 1259.32187 | 1.13 | 4.6E-07 | 2.9E-05 |
| *Slc19a2* | let-7a/c/d/f/k | | | Co-nodes | Transporters and tra | Solute carrier superfamily memb | 622.925861 | 0.69 | 4.0E-04 | 7.3E-03 |
| *Slc20A1* | | miRTarbase | Targetscan | Co-nodes | Transporters and tra | Solute carrier superfamily memb | 1918.10738 | 0.38 | 4.9E-03 | 4.6E-02 |
| *Slc25a24* | let-7a/b/c/d/k | | Targetscan | Co-nodes | Transporters and tra | Solute carrier superfamily memb | 1245.74345 | 0.49 | 1.0E-04 | 2.5E-03 |
| *Slc26a2* | let-7a/b/k | | | Co-nodes | Transporters and tra | Solute carrier superfamily memb | 2380.25985 | 0.36 | 1.4E-03 | 1.9E-02 |
| *Slc4a8* | let-7a/b/c | | Targetscan | Co-nodes | Transporters and tra | Solute carrier superfamily memb | 222.913264 | 1.52 | 4.3E-09 | 4.8E-07 |
| *Smpdl3B* | | miRTarbase | | Co-nodes | CNS proteins | Sphingomyelin phosphodiesteras | 93.6446063 | 1.41 | 1.9E-04 | 4.0E-03 |
| *Sod2* | | miRTarbase | Targetscan | Enzymes | Oxidoreductases | Superoxide dismutases (SOD) | 7162.25447 | 0.60 | 4.6E-04 | 8.2E-03 |
| *Sp100* | let-7b | | | Co-nodes | Nuclear proteins | SP100 nuclear antigen | 186.080396 | 0.71 | 5.4E-03 | 4.9E-02 |
| *Spcs3* | let-7a/d/f | | Targetscan | Co-nodes | Endosomal, lysosom | Signal peptidase complex subuni | 4405.84141 | 0.36 | 1.8E-03 | 2.3E-02 |
| *Spint1* | let-7a/b/c/f/k | | Targetscan | Receptors | Ligands | Serine peptidase inhibitor, Kunitz | 4136.90373 | 0.27 | 2.7E-03 | 3.0E-02 |
| *Spns2* | let-7b/c/k | | | Co-nodes | Lipid metabolism | Sphingolipid transporter | 289.313593 | 0.77 | 2.0E-03 | 2.4E-02 |
| *Srgap1* | let-7a/b/c/d/e | | Targetscan | Enzymes | Enzyme regulators | SLIT-ROBO Rho GTPase activatin | 76.2617512 | 1.01 | 2.0E-03 | 2.5E-02 |
| *Steap2* | let-7a/b/c/d/f | | | Enzymes | Other enzymes | STEAP | 4277.93806 | 0.60 | 4.6E-03 | 4.4E-02 |
| *Stx17* | let-7a/b/c/d/f | | Targetscan | Co-nodes | Endosomal, lysosom | Syntaxin | 3818.78496 | 0.32 | 2.4E-03 | 2.8E-02 |
| *Syk* | let-7a/b/c/d/f | | Targetscan | Enzymes | Kinases | Spleen associated tyrosine kinas | 333.632863 | 1.70 | 4.3E-07 | 2.7E-05 |
| *Tgm2* | let-7a/b/c/d/e/f/k | | Targetscan | Receptors | Ligands | Transglutaminases (TGM) | 4310.37955 | 0.53 | 5.6E-04 | 9.4E-03 |
| *Thyn1* | | miRTarbase | Targetscan | Co-nodes | Nuclear proteins | Thymocyte nuclear protein | 414.631052 | 0.74 | 4.0E-05 | 1.2E-03 |
| *Timp2* | let-7a/b/c/d/f/k | | | Receptors | Ligands | TIMP metallopeptidase inhibitor | 3686.93612 | 0.78 | 3.1E-11 | 6.3E-09 |
| *Tk1* | let-7a/c/f | | | Enzymes | Kinases | Thymidine kinases (TK) | 1035.9233 | 0.88 | 4.4E-08 | 3.7E-06 |
| *Tmem165* | | miRTarbase | | Co-nodes | Membrane proteins | Transmembrane protein | 2397.70557 | 0.40 | 8.2E-04 | 1.3E-02 |
| *Tmem30a* | let-7a/b/c/f | | Targetscan | Co-nodes | Membrane proteins | Transmembrane protein | 27127.872 | 0.40 | 7.9E-04 | 1.2E-02 |
| *Tmem37* | let-7a/b/c/d/k | | | Co-nodes | Membrane proteins | Transmembrane protein | 1373.80299 | 0.66 | 8.2E-05 | 2.1E-03 |
| *Tnfrsf10b* | let-7a/b/c/f | miRTarbase | | Receptors | Catalytic receptors | TRAIL receptors | 293.686687 | 0.92 | 8.5E-04 | 1.3E-02 |
| *Tnfrsf1b* | let-7a/b/c/d/e/f/k | | Targetscan | Receptors | Catalytic receptors | Tumor necrosis factor receptors | 1202.71534 | 0.58 | 4.2E-03 | 4.1E-02 |
| *Tpm4* | let-7a/b/c/d/e/f/k | | | Co-nodes | Cytoskeleton compo | Tropomyosin | 19221.4537 | 0.43 | 7.5E-05 | 2.0E-03 |
| *Trp53inp2* | let-7a/b/c/d/e/f | | | Co-nodes | Nuclear proteins | Tumor protein p53 inducible nuc | 11093.0241 | 0.49 | 2.4E-05 | 7.9E-04 |
| *Ttc22* | | miRTarbase | Targetscan | Co-nodes | Other co-nodes | Tetratricopeptide repeat domain | 261.037503 | 0.91 | 6.2E-05 | 1.7E-03 |
| *Tubb5* | let-7a/b/c/d/e/f/i/k | | | Co-nodes | Cytoskeleton compo | Tubulin | 18116.3059 | 0.40 | 5.4E-05 | 1.5E-03 |
| *Uhrf1* | | miRTarbase | | Enzymes | E3 ubiquitin ligases | Ubiquitin-like PHD and RING fing | 2255.82274 | 0.88 | 1.2E-08 | 1.2E-06 |
| *Vamp3* | let-7a/b/c/d/e/f/k | | Targetscan | Co-nodes | Endosomal, lysosom | Vesicle associated membrane pr | 3214.33735 | 0.44 | 2.0E-05 | 7.1E-04 |
| *Zeb2* | let-7a/b/c/d/f/g | | | Transcription | Homeo domain | Zn finger E-box binding homeobo | 184.802007 | 0.83 | 1.4E-03 | 1.8E-02 |
| *Zfp169* | let-7b/f/k | | Targetscan | Co-nodes | Zinc finger proteins | ZNF zinc finger protein | 558.627215 | 4.01 | 3.8E-62 | 5.6E-58 |
| *Zfp512b* | let-7a/b/c/d/e/f/k | | | Transcription | C2H2 Zn finger facto | ZNF512-factor | 1259.61116 | 0.42 | 4.7E-03 | 4.5E-02 |
| *Zmat3* | let-7a/b/d | | | Co-nodes | Zinc finger proteins | Zinc finger matrin-type | 1315.65888 | 0.70 | 3.4E-03 | 3.5E-02 |
| *Zyx* | let-7a/b/c/e/f/g/k | | | Co-nodes | Cytoskeleton compo | Zyxin | 1222.15819 | 0.49 | 2.6E-03 | 2.9E-02 |

Supplementary Data 2: *let-7* Targetome from *Sftpc*-tdT+ alveolar organoids

| Symbol | AGO2-miR-eCLIP+let-7 | miRTarbase (Experimentally Validated) | TargetScan Predicted | Category | Class | Family | Base mean expression | Log Fold Change | p value | Adjusted p value |
|---|---|---|---|---|---|---|---|---|---|---|
| *Abhd2* | let-7a/b/c/d/f/k | | | Enzymes | Esterases | Abhydrolase: | 4402.75092 | 0.89 | 1.5E-03 | 1.1E-02 |
| *Acsl1* | let-7a/c/f/ | | | Enzymes | Long chain fa | Long-chain a | 1015.15978 | 0.97 | 2.3E-06 | 5.7E-05 |
| *Acta2* | let-7a/b/e/f/k | | | Co-nodes | Cytoskeleton | Actin | 8.29573045 | 6.46 | 1.8E-03 | 1.3E-02 |
| *Actb* | | miRTarbase | | Co-nodes | Cytoskeleton | Actin | 44762.0343 | 0.95 | 2.6E-04 | 3.0E-03 |
| *Adamtsl3* | let-7b/d/k | | Targetscan | Enzymes | Peptidases | ADAMTS like | 105.72997 | 3.06 | 8.5E-11 | 6.9E-09 |
| *Adipor2* | let-7a/b/c/e/f | | Targetscan | Receptors | Other recept | Adiponectin | 6404.95912 | | | 8.7E-01 |
| *Adk* | let-7b/c/k | | Targetscan | Enzymes | Kinases | Adenosine ki | 2668.10713 | 1.33 | 2.4E-04 | 2.9E-03 |
| *Ak2* | let-7a/d | | | Enzymes | Kinases | Adenylate ki | 2430.47369 | 0.80 | 1.7E-04 | 2.2E-03 |
| *Akap5* | let-7a/b/c/d/f | | | Enzymes | Enzyme regu | A-kinase anc | 6158.36422 | 1.68 | 1.3E-05 | 2.4E-04 |
| *Angptl4* | let-7a/b/c/d/f | | | Receptors | Ligands | Angiopoietin | 114.833005 | 1.81 | 7.7E-04 | 7.0E-03 |
| *Ap1ar* | let-7b/k | | | Co-nodes | Adaptor, doc | Adaptor rela | 1263.56398 | 0.56 | 2.9E-03 | 1.9E-02 |
| *Ap1m1* | let-7a/b/k | | | Co-nodes | Adaptor, doc | Adaptor rela | 803.579857 | 0.75 | 2.2E-03 | 1.5E-02 |
| *Apbb2* | let-7b/f | | | Co-nodes | Amyloid prot | Amyloid beta | 924.23488 | 0.75 | 8.7E-03 | 4.4E-02 |
| *Arf4* | let-7a/b/c/d/e/f | | Targetscan | Enzymes | GTPases | ADP-ribosyla | 6421.64575 | 0.43 | 8.6E-03 | 4.3E-02 |
| *Arhgap1* | let-7a/b/c/d/f/k | | | Enzymes | Enzyme regu | Rho GTPase | 3574.08244 | 0.79 | 2.4E-04 | 2.8E-03 |
| *Arhgdia* | let-7a/b/c/d/e/f/g/i/k | | | Enzymes | Enzyme regu | Rho GDP-dis | 9038.05695 | 0.88 | 6.7E-06 | 1.4E-04 |
| *Arl2* | let-7a/c/f/k | | | Enzymes | GTPases | ADP-ribosyla | 634.998508 | 0.77 | 5.8E-04 | 5.7E-03 |
| *Atl3* | let-7c/d/e/f | | Targetscan | Enzymes | GTPases | Atlastin GTP | 5575.88948 | 0.45 | 9.3E-03 | 4.5E-02 |
| *Atxn7l3b* | let-7a/b/c/d/f/k | | Targetscan | Co-nodes | Other co-noc | Ataxin | 3142.75642 | 0.48 | 4.7E-03 | 2.8E-02 |
| *Aurkb* | | miRTarbase | | Enzymes | Kinases | Aurora-relat | 1078.16546 | 0.84 | 4.2E-04 | 4.5E-03 |
| *Aven* | let-7a/b/c/d/f/g/i/k | | Targetscan | Enzymes | Enzyme regu | Apoptosis ar | 256.689017 | 1.33 | 9.9E-05 | 1.4E-03 |
| *Bcap29* | let-7b/k | | Targetscan | Co-nodes | Receptor ass | B cell recept | 1005.79444 | 0.95 | 1.6E-03 | 1.2E-02 |
| *Bcl2l1* | let-7b/d/g | | Targetscan | Enzymes | Enzyme regu | Protein phos | 3761.54234 | 0.79 | 7.0E-04 | 6.6E-03 |
| *Bgn* | let-7a/b/c/d/e/f/g/i/k | | | Receptors | Ligands | Biglycan | 64.60061 | 3.64 | 1.8E-03 | 1.3E-02 |
| *Bloc1s6* | let-7a/b/c/f | | Targetscan | Co-nodes | Endosomal, l | Biogenesis o | 595.610284 | | | 8.8E-01 |
| *Bysl* | let-7a/b/c/d/e | | Targetscan | Co-nodes | Other co-noc | Bystin like | 768.297379 | 1.23 | 2.9E-04 | 3.3E-03 |
| *Cables2* | let-7b/k | | | Co-nodes | Substrate pr | Cdk5 and Ab | 561.40218 | 0.82 | 3.4E-03 | 2.2E-02 |
| *Capn6* | let-7a/b/c/d/e/f/k | | | Enzymes | Peptidases | Calpains (CA | 626.614299 | 1.93 | 1.1E-08 | 5.5E-07 |
| *Car2* | let-7a/b/c/d/f/k | | Targetscan | Enzymes | Dehydratase | Carbonic anh | 1658.13751 | 1.81 | 9.2E-09 | 4.5E-07 |
| *Cbx5* | let-7a/b/c/d/e/f/g/i/k | miRTarbase | Targetscan | Co-nodes | Transcription | Chromobox | 7818.83058 | 0.63 | 8.0E-03 | 4.1E-02 |
| *Ccdc25* | let-7a/b/c/d/f/k | | Targetscan | Co-nodes | Coiled coil do | Coiled-coil do | 1330.66665 | 0.83 | 1.5E-05 | 2.8E-04 |
| *Ccnb2* | | miRTarbase | | Enzymes | Enzyme regu | Cyclins (CCN | 1284.19192 | 0.53 | 5.2E-03 | 3.0E-02 |
| *Ccnyl1* | let-7a/b/c/d/e/k | | Targetscan | Enzymes | Enzyme regu | Cyclin Y like | 484.897456 | 0.64 | 9.6E-03 | 4.7E-02 |
| *Cd276* | let-7a/b/c/d/f | | Targetscan | Co-nodes | Cell surface | Cluster of dif | 1381.14894 | 0.59 | 2.5E-03 | 1.7E-02 |
| *Cdc34* | let-7a/b/c/d/e/f/k | miRTarbase | Targetscan | Enzymes | Aminoacyltra | Cell division | 1731.61577 | 0.86 | 6.0E-06 | 1.3E-04 |
| *Cfl1* | let-7a/b/c/d/e/f/i | | | Co-nodes | Other co-noc | Cofilin | 13877.9958 | 0.65 | 4.8E-05 | 7.5E-04 |
| *Clcn5* | let-7a/b/c/d/f | | Targetscan | Ion channels | Other ion cha | Chloride volt | 1203.16649 | 1.03 | 1.1E-04 | 1.5E-03 |
| *Clic4* | let-7a/b/c/d/e/f/k | | | Ion channels | Other ion cha | Chloride intra | 3252.01055 | 0.58 | 6.2E-04 | 6.0E-03 |
| *Clic5* | let-7a/b/c/d/e/f/k | | | Ion channels | Other ion cha | Chloride intra | 3701.26957 | 1.78 | 2.5E-07 | 8.5E-06 |
| *Clip1* | let-7a/b/c/d/e/f | | | Co-nodes | Other co-noc | CAP-Gly dom | 2180.13319 | 0.65 | 1.4E-03 | 1.1E-02 |
| *Cnih1* | let-7a/b/c/d/e/f/k | | | Co-nodes | Receptor ass | Cornichon AM | 2518.95544 | 0.58 | 4.7E-03 | 2.8E-02 |
| *Col1a1* | let-7a/b/c/d/e/f/g/i/k | | Targetscan | Receptors | Ligands | Collagen cha | 63.5055923 | 2.72 | 4.9E-07 | 1.5E-05 |
| *Col4a3* | let-7a/b/c/f/k | | | Receptors | Ligands | Collagen cha | 444.187894 | 2.38 | 9.7E-08 | 3.6E-06 |
| *Copz1* | let-7b/c/e/f/k | miRTarbase | | Co-nodes | Proteasome | COPI coat co | 3257.58481 | 0.57 | 6.8E-03 | 3.6E-02 |
| *Crip2* | let-7a/b/c/d/e/f/i/k | | | Co-nodes | LIM domain | Cysteine rich | 4528.79245 | 1.21 | 7.0E-08 | 2.7E-06 |
| *Ctgf* | let-7a/b/c/k | | | Receptors | Ligands | Cellular com | 540.042269 | 1.26 | 7.6E-07 | 2.2E-05 |
| *Ctsh* | let-7a/b/c/k | | | Enzymes | Peptidases | Cathepsins (( | 43207.3507 | 1.38 | 2.7E-05 | 4.7E-04 |
| *Cttnbp2nl* | let-7b | | | Co-nodes | Other co-noc | CTTNBP2 N-t | 1645.38835 | 0.96 | 1.0E-04 | 1.4E-03 |
| *Cux1* | let-7a/b/c/d | | Targetscan | Transcription | Homeo doma | CUX | 4791.24359 | 0.49 | 4.4E-03 | 2.6E-02 |
| *Cyb5r3* | let-7a/b/c/d/f/k | | | Enzymes | Reductases | Cytochrome- | 5695.24959 | 0.69 | 2.3E-04 | 2.7E-03 |
| *Dab2* | let-7a/b/c/d/f | | | Co-nodes | Adaptor, doc | DAB adaptor | 18.9883197 | 2.71 | 5.8E-03 | 3.2E-02 |
| *Dcakd* | let-7a/b/d/k | | Targetscan | Co-nodes | Other co-noc | Dephospho-C | 1176.58517 | 1.11 | 6.4E-06 | 1.4E-04 |
| *Dennd3* | let-7a/b/c/d/f | | Targetscan | Enzymes | Enzyme regu | DENN domai | 1931.6713 | 1.15 | 5.3E-04 | 5.4E-03 |
| *Dmd* | let-7a/b/c/d/f/k | | Targetscan | Co-nodes | Cytoskeleton | Dystrophin | 565.407297 | 0.62 | 6.3E-03 | 3.5E-02 |
| *Dna2* | | miRTarbase | Targetscan | Enzymes | Helicases (DI | DNA2/NAM7 | 554.144639 | 1.13 | 6.5E-03 | 3.5E-02 |

| | | | | | | | | | | |
|---|---|---|---|---|---|---|---|---|---|---|
| *Dpy19l1* | let-7b/k | | | Enzymes | Glycosyltrans | Dpy-19 like C | 639.624126 | 0.87 | 1.1E-04 | 1.5E-03 |
| *E2f1* | | miRTarbase | | Transcription | E2F/FOX | E2F | 533.778086 | 0.81 | 3.9E-04 | 4.2E-03 |
| *E2f2* | | miRTarbase | Targetscan | Transcription | E2F/FOX | E2F | 1122.90677 | 1.42 | 9.0E-04 | 8.0E-03 |
| *E2f6* | let-7a/b/c/d/e/f/i/k | miRTarbase | Targetscan | Transcription | E2F/FOX | E2F | 771.174126 | 0.61 | 3.0E-03 | 2.0E-02 |
| *Edil3* | let-7a/b | | | Receptors | Ligands | EGF like repe | 48.0115771 | 2.58 | 1.3E-04 | 1.7E-03 |
| *Ehd1* | let-7b/c | | | Co-nodes | Calcium asso | EH domain c | 1977.30381 | 0.70 | 1.9E-03 | 1.4E-02 |
| *Ehd2* | let-7a/b/c/d/f/i/k | | | Co-nodes | Calcium asso | EH domain c | 2468.57943 | 1.92 | 4.1E-27 | 3.8E-24 |
| *Eif1ad* | let-7a/b/c/d/f/k | | Targetscan | Co-nodes | Translation f | Eukaryotic tr | 698.313925 | 0.68 | 3.2E-03 | 2.0E-02 |
| *Eif4a1* | let-7a/b/c/d/f/k | | | Enzymes | Helicases (R | Eukaryotic in | 27870.714 | 0.47 | 3.5E-03 | 2.2E-02 |
| *Eif4e2* | let-7b | | | Co-nodes | Translation f | Eukaryotic tr | 1950.16909 | 0.63 | 3.2E-04 | 3.6E-03 |
| *Eif4g2* | let-7a/b/c/d/e/f/k | miRTarbase | Targetscan | Co-nodes | Translation f | Eukaryotic tr | 21989.5247 | 0.57 | 3.5E-04 | 3.8E-03 |
| *Eif5a2* | let-7e/f | | | Co-nodes | Translation f | Eukaryotic tr | 145.47285 | 1.14 | 3.3E-03 | 2.1E-02 |
| *Epb41l5* | let-7b/c/f/k | | | Co-nodes | Membrane p | Erythrocyte m | 2394.57203 | 0.86 | 1.9E-04 | 2.4E-03 |
| *Etf1* | let-7a/b/c/e/g/k | | | Co-nodes | Translation f | Eukaryotic tr | 5653.58653 | 0.46 | 7.5E-03 | 3.9E-02 |
| *Ezh2* | | miRTarbase | | Co-nodes | Polycomb gr | Enhancer of | 1148.65854 | 0.67 | 5.9E-04 | 5.8E-03 |
| *Fam102b* | let-7b | | | Co-nodes | Family with | Family with | 7985.21527 | 0.72 | 2.4E-03 | 1.6E-02 |
| *Fam98a* | let-7b | | | Co-nodes | Family with | Family with | 1585.7669 | 0.72 | 7.0E-04 | 6.6E-03 |
| *Fgf1* | let-7a/b/c/d/f/k | | | Receptors | Ligands | Fibroblast gr | 2358.36849 | 1.05 | 9.5E-04 | 8.3E-03 |
| *Fkbp1a* | let-7a/b/c/d/e/f/k | | | Enzymes | Peptidylproly | FK506-bindir | 3304.1449 | 1.23 | 6.2E-05 | 9.3E-04 |
| *Fkrp* | let-7a/b/c/f | | Targetscan | Co-nodes | Other co-noc | Fukutin relat | 897.713148 | 0.64 | 3.2E-03 | 2.0E-02 |
| *Foxp2* | let-7b/c/d/f | | Targetscan | Transcription | E2F/FOX | FOXP | 2684.70083 | 1.01 | 8.7E-04 | 7.8E-03 |
| *Fxn* | | miRTarbase | Targetscan | Enzymes | Oxidases | Frataxin | 258.570292 | 0.79 | 7.1E-03 | 3.8E-02 |
| *Gata6* | let-7b | | | Transcription | Other transc | Two zinc-fing | 1291.6989 | 0.85 | 1.0E-02 | 4.9E-02 |
| *Glrx* | let-7a/b/c/d/f/g/k | | Targetscan | Enzymes | Other enzym | Glutaredoxin | 3635.86294 | 1.57 | 1.0E-06 | 2.8E-05 |
| *Gnas* | let-7a/b/c/d/f/k | | | Receptors | Ligands | GNAS compl | 16116.0439 | 0.55 | 5.9E-04 | 5.8E-03 |
| *Gng5* | | miRTarbase | Targetscan | Receptors | G protein co | G protein sul | 2493.1104 | 0.51 | 7.6E-03 | 4.0E-02 |
| *Golt1b* | let-7a/b/c/f | | Targetscan | Co-nodes | Endosomal, | Golgi transpo | 927.47509 | 0.73 | 8.4E-04 | 7.5E-03 |
| *Gorasp2* | | miRTarbase | | Co-nodes | Endosomal, | Golgi reasse | 3353.65513 | 0.72 | 4.3E-04 | 4.5E-03 |
| *Gpat4* | | miRTarbase | | Enzymes | Acyltransfera | Glycerol-3-pl | 1993.39092 | 0.68 | 2.4E-03 | 1.6E-02 |
| *Gpd2* | let-7a/b/d | | | Enzymes | Dehydrogena | Glycerol-3-pl | 1847.1703 | 0.59 | 1.2E-03 | 1.0E-02 |
| *Gpn1* | | miRTarbase | | Enzymes | GTPases | GPN-loop GT | 779.206134 | 0.74 | 4.3E-03 | 2.6E-02 |
| *Gpr155* | let-7b/c | | Targetscan | Receptors | G protein co | G protein-co | 350.804706 | 1.31 | 6.6E-04 | 6.3E-03 |
| *Gprc5a* | let-7a/b/c/d/e/f/k | | | Receptors | G protein co | G protein-co | 18469.7539 | 0.92 | 3.6E-03 | 2.3E-02 |
| *Grb10* | let-7a/c/f/g/k | | Targetscan | Co-nodes | Receptor ass | Growth facto | 2272.36488 | 1.39 | 2.0E-03 | 1.4E-02 |
| *Heatr3* | let-7a/b/c/d/f/k | | | Co-nodes | Other co-noc | HEAT repeat | 1359.99555 | 0.63 | 4.0E-03 | 2.5E-02 |
| *Hif1an* | let-7a/b/c/d/e/f/k | | Targetscan | Enzymes | Dioxygenase | Hypoxia indu | 2788.58784 | 0.50 | 6.2E-03 | 3.4E-02 |
| *Hipk2* | let-7a/b/c/d/e/f/k | | Targetscan | Enzymes | Kinases | Homeodoma | 564.637267 | 1.01 | 8.4E-05 | 1.2E-03 |
| *Hmgn2* | | miRTarbase | Targetscan | Co-nodes | Transcription | High mobility | 4818.83207 | 1.01 | 6.2E-05 | 9.3E-04 |
| *Hs2st1* | let-7b/f | | Targetscan | Enzymes | Sulfotransfer | Heparan sulf | 4110.75025 | 1.00 | 6.7E-05 | 9.9E-04 |
| *Hsd17b11* | let-7a/b/c/k | | | Enzymes | Dehydrogena | 17 beta hydr | 1213.6599 | 0.78 | 9.0E-03 | 4.5E-02 |
| *Il17re* | let-7b/f/k | | | Receptors | Catalytic rec | Interleukin 1 | 555.605955 | 0.82 | 8.8E-04 | 7.8E-03 |
| *Ilk* | let-7a/b/c/f | | | Enzymes | Kinases | Integrin linke | 2214.98663 | 0.75 | 4.5E-05 | 7.2E-04 |
| *Imp4* | let-7a/b/c/d/f | | Targetscan | Co-nodes | Ribosomes a | IMP U3 smal | 988.631307 | 0.62 | 1.5E-03 | 1.2E-02 |
| *Ino80c* | let-7b/k | | | Co-nodes | Chromatin as | INO80 comp | 840.185326 | 0.70 | 1.5E-03 | 1.2E-02 |
| *Iqgap3* | let-7b/c/k | | | Enzymes | Enzyme regu | IQ motif con | 2895.9385 | 1.28 | 8.0E-04 | 7.2E-03 |
| *Itgb3* | let-7a/b/c/d/e/f/k | miRTarbase | Targetscan | Receptors | Catalytic rec | Integrins | 21.4563158 | 3.24 | 6.8E-04 | 6.4E-03 |
| *Itgb6* | let-7a/b/c | | | Receptors | Catalytic rec | Integrins | 1163.10172 | 1.09 | 2.3E-05 | 4.1E-04 |
| *Itpripl2* | let-7a/b/c/e/k | | | Co-nodes | Receptor ass | ITPRIP like | 2797.44932 | 0.55 | 6.8E-03 | 3.6E-02 |
| *Kctd10* | let-7a/b/c/d/f/g/k | | | Ion channels | Voltage gate | Potassium ch | 5618.90195 | 0.79 | 4.9E-04 | 5.0E-03 |
| *Knstrn* | let-7a/b | | | Co-nodes | Cell cycle, ce | Kinetochore | 1561.3531 | 0.61 | 2.2E-03 | 1.6E-02 |
| *Kras* | let-7b/k | miRTarbase | | Enzymes | GTPases | Ras Type GT | 1736.1385 | 0.62 | 4.3E-04 | 4.6E-03 |
| *Krt80* | let-7a/b/c/d/f/i/k | | | Co-nodes | Cytoskeleton | Keratin | 892.154701 | 1.50 | 6.2E-12 | 6.3E-10 |
| *Lasp1* | let-7a/b/c/k | | | Co-nodes | LIM domain | LIM and SH3 | 14459.238 | 0.86 | 6.6E-04 | 6.3E-03 |
| *Lhfp* | let-7b/c/d/f/g/k | | | Co-nodes | Other co-noc | Lipoma HMG | 207.03559 | 1.70 | 4.6E-03 | 2.7E-02 |
| *Lrrc20* | | miRTarbase | Targetscan | Co-nodes | Leucine rich | Leucine rich | 275.945719 | 0.83 | 9.9E-03 | 4.8E-02 |
| *Lrrc59* | let-7a/b/c/d/k | | Targetscan | Co-nodes | Leucine rich | Leucine rich | 7588.96821 | 0.96 | 1.8E-04 | 2.2E-03 |
| *Mal2* | let-7a/b/c/d/e/f/k | | | Co-nodes | Differentiatio | Mal, T cell di | 11915.5676 | 0.82 | 1.1E-03 | 9.3E-03 |
| *Map2k3* | let-7b/c | | | Enzymes | Kinases | Mitogen-acti | 2215.69458 | 0.68 | 4.8E-04 | 4.9E-03 |
| *Mapre2* | let-7a/b/c/f | | | Co-nodes | Cytoskeleton | Microtubule | 1851.71582 | 1.27 | 1.2E-05 | 2.4E-04 |
| *Marveld2* | let-7a/b/c/d/f | | Targetscan | Co-nodes | Other co-noc | MARVEL don | 765.173491 | 0.72 | 3.8E-04 | 4.1E-03 |
| *Mbnl1* | let-7a/b/c/e/f/i/k | | | Co-nodes | Splicing facto | Muscleblind | 7150.7418 | 1.08 | 1.4E-06 | 3.7E-05 |
| *Mbnl3* | let-7a/c/d/f | | Targetscan | Co-nodes | Splicing facto | Muscleblind | 1126.784 | 0.62 | 3.3E-03 | 2.1E-02 |
| *Mical1* | let-7a/b/c/d/k | | | Enzymes | Monooxygen | Molecule int | 1206.43957 | 2.64 | 1.2E-08 | 5.9E-07 |
| *Mlf2* | let-7a/b/c/d/e/f/k | | | Co-nodes | Other co-noc | Myeloid leuk | 4895.32857 | 0.60 | 2.6E-04 | 3.0E-03 |
| *Msn* | let-7a/b/c/d/f/k | | Targetscan | Co-nodes | Cytoskeleton | Moesin | 9995.31011 | 0.82 | 3.1E-03 | 2.0E-02 |
| *Mtfr1l* | let-7a/b/c/d/k | miRTarbase | | Co-nodes | Mitochondria | Mitochondria | 1959.10398 | 0.81 | 5.1E-03 | 2.9E-02 |

| | | | | | | | | | | |
|---|---|---|---|---|---|---|---|---|---|---|
| *Myh10* | let-7a/c/d/f | | | Co-nodes | Cytoskeleton | Myosin heav | 722.016156 | 1.50 | 1.3E-03 | 1.1E-02 |
| *Myo1c* | let-7a/b/c/d/e/f/k | | | Co-nodes | Cytoskeleton | Myosin | 6259.148 | 0.65 | 1.4E-03 | 1.1E-02 |
| *Naaa* | let-7b/c/d/f | | | Enzymes | Other enzym | N-acylethano | 471.353358 | 0.80 | 5.0E-04 | 5.1E-03 |
| *Nap1l1* | let-7b | miRTarbase | Targetscan | Co-nodes | Cell cycle, ce | Nucleosome | 7824.89419 | 0.51 | 2.4E-03 | 1.7E-02 |
| *Ncbp1* | | miRTarbase | Targetscan | Co-nodes | Nuclear prot | Nuclear cap | 2069.65556 | 0.64 | 7.9E-03 | 4.1E-02 |
| *Nceh1* | let-7a/c/d/f | | Targetscan | Enzymes | Other enzym | Neutral chole | 1356.61316 | 0.75 | 1.7E-03 | 1.3E-02 |
| *Nectin3* | let-7a/b/c/d/e/f/g/i | | | Co-nodes | Adhesion mo | Nectin cell a | 1616.5274 | 1.98 | 1.3E-05 | 2.5E-04 |
| *Nek6* | let-7a/b/c/e | | | Enzymes | Kinases | NimA related | 3179.96754 | 0.80 | 6.7E-03 | 3.6E-02 |
| *Nme6* | let-7b/d/e/f | | Targetscan | Enzymes | Kinases | NM23 kinase | 251.327975 | 0.84 | 7.8E-03 | 4.0E-02 |
| *Nol4l* | let-7a/b/c/d/e/f/i/k | | Targetscan | Co-nodes | Ribosomes a | Nucleolar pr | 1685.36989 | 1.70 | 1.1E-06 | 3.0E-05 |
| *Nolc1* | | miRTarbase | | Co-nodes | Ribosomes a | Nucleolar an | 1723.83252 | 0.96 | 7.0E-04 | 6.6E-03 |
| *Npnt* | let-7a/b/e/f/k | | | Receptors | Ligands | Nephronecti | 2808.03702 | 1.65 | 1.9E-05 | 3.5E-04 |
| *Nrep* | let-7a/b/c/e/f/g | | | Co-nodes | CNS Proteins | Neuronal reg | 2956.57583 | 1.66 | 2.2E-05 | 4.0E-04 |
| *Ntn1* | let-7a/b/c/d/e/f/g | | Targetscan | Receptors | Ligands | Netrin | 139.872032 | 1.66 | 1.0E-03 | 8.6E-03 |
| *Nucks1* | let-7a/b/c/d/f/k | | | Co-nodes | Cell cycle, ce | Nuclear case | 6794.76018 | 0.69 | 5.0E-05 | 7.7E-04 |
| *Nup62* | let-7a/b/c/d/f/k | | Targetscan | Co-nodes | Nuclear pore | Nucleoporin | 1864.35317 | 0.59 | 9.2E-03 | 4.5E-02 |
| *Opa3* | | miRTarbase | Targetscan | Co-nodes | Mitochondria | Outer mitoch | 890.963297 | 0.71 | 2.8E-03 | 1.9E-02 |
| *Oxct1* | let-7b/c/f | | | Enzymes | Transferases | 3-oxoacid Co | 6011.7711 | 0.51 | 3.7E-03 | 2.3E-02 |
| *Pafah1b2* | let-7a/b/c/d/k | | | Enzymes | Other enzym | Platelet-activ | 3536.44913 | 0.45 | 7.8E-03 | 4.0E-02 |
| *Paqr4* | let-7b/k | | | Receptors | Other recept | Progesterone | 256.44058 | 1.26 | 7.8E-06 | 1.6E-04 |
| *Pctp* | let-7a/b/c/d/f/k | | Targetscan | Co-nodes | Other co-noc | Phosphatidyl | 426.107507 | 0.85 | 3.3E-04 | 3.7E-03 |
| *Pcyt1a* | let-7a/b/c/d | | | Enzymes | Nucleotidyltr | Phosphate cy | 2989.15056 | 1.16 | 6.8E-07 | 2.0E-05 |
| *Pepd* | let-7b/c/f | | | Enzymes | Peptidases | Peptidase D | 1485.88703 | 0.95 | 1.7E-04 | 2.2E-03 |
| *Pfn1* | let-7a/b/c/f | | | Co-nodes | Cytoskeleton | Profilin | 7361.61061 | 0.82 | 3.0E-06 | 7.0E-05 |
| *Plaur* | let-7a/b/c/d/e/f/g | | Targetscan | Receptors | Other recept | Plasminogen | 1008.05936 | 0.89 | 1.6E-04 | 2.0E-03 |
| *Plxnd1* | let-7a/b/c/d/e/f/i/k | miRTarbase | Targetscan | Receptors | Other recept | Plexin | 3278.35747 | 0.99 | 1.0E-02 | 4.9E-02 |
| *Polr3d* | let-7a/c/f/i/k | miRTarbase | Targetscan | Enzymes | Nucleotidyltr | DNA-directed | 701.409485 | 0.65 | 6.3E-03 | 3.5E-02 |
| *Pphln1* | let-7a/b/f/k | | Targetscan | Co-nodes | Other co-noc | Periphilin | 1041.99606 | 0.75 | 1.3E-04 | 1.8E-03 |
| *Ppic* | let-7a/b/c/d/e/f/g/i/k | | | Enzymes | Peptidylproly | Peptidyl-prol | 9363.70431 | 0.60 | 2.4E-03 | 1.6E-02 |
| *Ppp1r16b* | let-7a/b/c/d/f/k | | Targetscan | Enzymes | Enzyme regu | Protein phos | 9.55601406 | 5.18 | 6.3E-03 | 3.5E-02 |
| *Pptc7* | let-7b/c/k | | Targetscan | Enzymes | Phosphatase | Protein phos | 1573.28395 | 0.66 | 1.0E-03 | 8.6E-03 |
| *Prkar2a* | let-7a/b/c/d/e/f | | Targetscan | Enzymes | Enzyme regu | Protein kinas | 3120.1113 | 1.47 | 8.4E-10 | 5.4E-08 |
| *Psma2* | | miRTarbase | | Enzymes | Peptidases | Proteasome | 4858.66886 | 0.46 | 9.0E-03 | 4.5E-02 |
| *Psmd1* | let-7b/k | | | Co-nodes | Proteasome | Proteasome | 5027.85733 | 0.66 | 4.5E-03 | 2.7E-02 |
| *Psme3* | let-7d | | | Co-nodes | Proteasome | Proteasome | 2539.5724 | 0.69 | 2.7E-04 | 3.1E-03 |
| *Qk* | let-7a/b/c/d/f/k | | | Co-nodes | Other co-noc | Quaking | 3728.65281 | 1.12 | 2.6E-05 | 4.5E-04 |
| *Rab12* | let-7a/b/c/f/k | | | Enzymes | GTPases | RAB, membe | 1366.73836 | 0.71 | 9.0E-04 | 8.0E-03 |
| *Rab1b* | let-7a/b/c/d/f/k | | | Enzymes | GTPases | RAB, membe | 4771.30371 | 0.46 | 4.2E-03 | 2.6E-02 |
| *Rabl3* | let-7b/c/d/f/k | | Targetscan | Enzymes | GTPases | RAB, membe | 303.920044 | 0.74 | 5.5E-03 | 3.1E-02 |
| *Rad21* | let-7d | miRTarbase | | Co-nodes | Cohesin com | Cell cycle che | 6687.66816 | 0.43 | 7.1E-03 | 3.8E-02 |
| *Rap1b* | let-7a/b/c/d/f | | Targetscan | Enzymes | GTPases | Ras Type GT | 4646.78631 | 0.49 | 3.1E-03 | 2.0E-02 |
| *Rap2a* | let-7a/b/d | | | Enzymes | GTPases | Ras Type GT | 914.836714 | 0.72 | 3.0E-03 | 2.0E-02 |
| *Rbm38* | let-7a/b/d/f | | Targetscan | Co-nodes | RNA binding | RNA binding | 281.181084 | 1.89 | 3.2E-04 | 3.6E-03 |
| *Rbpms* | let-7a/b/c/d/e/k | | | Co-nodes | RNA binding | RNA binding | 1432.14782 | 0.64 | 7.8E-03 | 4.0E-02 |
| *Rdh10* | let-7a/b/c/d/f | | Targetscan | Enzymes | Dehydrogena | Retinol (RDH | 1163.19829 | 0.82 | 1.3E-04 | 1.8E-03 |
| *Rgs4* | let-7b/c | | | Enzymes | Enzyme regu | Regulator of | 952.673203 | 1.57 | 6.1E-03 | 3.4E-02 |
| *Rhoa* | let-7a/b/f | | | Enzymes | GTPases | Rho GTPases | 7580.21245 | 0.45 | 5.7E-03 | 3.2E-02 |
| *Rnf122* | let-7a/b/c | | | Co-nodes | Ring finger p | Ring finger p | 491.749981 | 0.84 | 6.4E-04 | 6.2E-03 |
| *Rnf5* | let-7a/b/c/d/e/f/i/k | | Targetscan | Enzymes | E3 ubiquitin | RING finger | 1167.18041 | 0.78 | 7.5E-04 | 6.9E-03 |
| *Rrm1* | | miRTarbase | Targetscan | Enzymes | Reductases | Ribonucleosi | 5766.44933 | 0.63 | 5.9E-03 | 3.3E-02 |
| *Rrm2* | | miRTarbase | Targetscan | Enzymes | Enzyme regu | Ribonucleosi | 1991.3756 | 1.18 | 1.9E-05 | 3.5E-04 |
| *Scamp3* | let-7a/b/c/d/f/k | | | Co-nodes | Membrane p | Secretory car | 1172.67349 | 0.99 | 1.6E-04 | 2.1E-03 |
| *Sec14l1* | let-7a/b/c/d/k | | Targetscan | Co-nodes | Lipid metabo | SEC14 like lip | 7919.92089 | 0.86 | 1.8E-06 | 4.5E-05 |
| *Sema5a* | let-7a/b/c/k | | Targetscan | Receptors | Ligands | Semaphorin | 564.733667 | 1.23 | 4.7E-03 | 2.8E-02 |
| *Sema7a* | let-7a/b/c/d/f/k | | | Receptors | Ligands | Semaphorin | 73.3903182 | 1.55 | 1.2E-03 | 9.9E-03 |
| *Senp2* | let-7a/b/c/f | | Targetscan | Enzymes | Peptidases | Sentrin-spec | 1717.27906 | 0.62 | 3.8E-03 | 2.4E-02 |
| *Serpinb6b* | let-7b/c/f/k | | Targetscan | NA | NA | NA | 3698.15345 | 1.49 | 3.6E-06 | 8.3E-05 |
| *Serpinb9* | let-7a/b/c/d | | | Enzymes | Enzyme regu | Serpin B | 4076.5413 | 1.54 | 1.0E-06 | 2.8E-05 |
| *Sesn3* | let-7a/b/c/d/f/k | | Targetscan | Enzymes | Peroxidases | Sestrin | 1608.7034 | 0.56 | 7.4E-03 | 3.9E-02 |
| *Sf3a1* | | miRTarbase | Targetscan | Co-nodes | Splicing facto | Splicing facto | 2238.06587 | 0.51 | 9.0E-03 | 4.5E-02 |
| *Ska3* | | miRTarbase | | Co-nodes | Cell cycle, ce | Spindle and k | 397.389668 | 1.08 | 3.6E-04 | 4.0E-03 |
| *Slc26a9* | let-7a/b/c/f/k | | Targetscan | Co-nodes | Transporters | Solute carrie | 4356.39066 | 0.93 | 1.0E-02 | 5.0E-02 |
| *Slc35a4* | | miRTarbase | | Co-nodes | Transporters | Solute carrie | 2046.60637 | 0.72 | 1.5E-04 | 2.0E-03 |
| *Slc35b4* | let-7a/c/d/f | | | Co-nodes | Transporters | Solute carrie | 819.958761 | 0.80 | 9.1E-04 | 8.1E-03 |
| *Slc6a14* | let-7b/c/d | | | Co-nodes | Transporters | Solute carrie | 3078.01929 | 1.23 | 6.7E-03 | 3.6E-02 |
| *Slco2a1* | let-7a/b/c/d/e/f/g/i/k | | Targetscan | Co-nodes | Transporters | Solute carrie | 5928.07258 | 1.47 | 7.2E-03 | 3.8E-02 |

| | | | | | | | | | | |
|---|---|---|---|---|---|---|---|---|---|---|
| *Smarcc1* | let-7a/b/c/f/i | | Targetscan | Co-nodes | Chromatin as | SWI/SNF rel | 3238.29527 | 0.66 | 2.7E-03 | 1.8E-02 |
| *Snx18* | let-7a/d/e/f/k | | | Co-nodes | Membrane p | Sorting nexin | 2367.17972 | 0.84 | 4.3E-03 | 2.6E-02 |
| *Sparc* | let-7a/b/c/d/e/f/g/i/k | | | Co-nodes | Cysteine rich | Secreted pro | 9746.45927 | 2.11 | 3.3E-26 | 3.0E-23 |
| *Spock2* | let-7b/c/k | | | Co-nodes | Extracellular | SPARC (osteo | 55.4412191 | 1.93 | 6.6E-04 | 6.3E-03 |
| *Srsf2* | | miRTarbase | | Co-nodes | Splicing facto | Serine and a | 5753.70311 | 0.64 | 1.8E-03 | 1.4E-02 |
| *St13* | let-7a/b/c/d/e/f/k | | Targetscan | Co-nodes | Interacting p | ST13 Hsp70 i | 3622.46649 | 0.44 | 9.8E-03 | 4.7E-02 |
| *Stard3nl* | let-7a/b/c/d/f | | Targetscan | Co-nodes | Other co-nod | STARD3 N-te | 895.398345 | 0.65 | 7.3E-03 | 3.8E-02 |
| *Stk4* | let-7a/b/c/f | miRTarbase | | Enzymes | Kinases | Hippo/MST k | 2097.33733 | 0.63 | 1.1E-03 | 9.5E-03 |
| *Surf4* | let-7a/b/c/d/e/f/k | miRTarbase | Targetscan | Co-nodes | Other co-nod | Surfeit | 8888.62272 | 0.57 | 7.5E-03 | 3.9E-02 |
| *Taf9b* | let-7a/b/c/d/k | | Targetscan | Co-nodes | General tran | TATA-box bin | 281.776215 | 1.17 | 2.5E-03 | 1.7E-02 |
| *Tagln2* | let-7a/b/c/d/e/f/k | | | Co-nodes | Cytoskeleton | Transgelin | 6809.75502 | 0.47 | 2.9E-03 | 1.9E-02 |
| *Tcf7l1* | let-7a/b/c/d/f/k | | Targetscan | Transcription | HMG domain | TCF-7-relate | 960.613413 | 1.32 | 3.8E-05 | 6.3E-04 |
| *Tgfb3* | let-7b/d/f | | | Receptors | Ligands | Transforming | 3889.35103 | 0.70 | 3.3E-05 | 5.5E-04 |
| *Tgfbrap1* | let-7a/b/c/d/f/k | | | Co-nodes | Receptor ass | Transforming | 948.752134 | 0.73 | 1.9E-03 | 1.4E-02 |
| *Them6* | | miRTarbase | | Enzymes | Other enzym | Thioesterase | 318.444458 | 0.77 | 3.6E-03 | 2.3E-02 |
| *Thsd4* | let-7b/k | | Targetscan | Co-nodes | Other co-nod | Thrombospo | 2118.71231 | 0.76 | 6.2E-03 | 3.4E-02 |
| *Tma7* | let-7a/b/c/d/f | | Targetscan | Co-nodes | Membrane p | Translation m | 2431.87207 | 0.57 | 1.7E-03 | 1.3E-02 |
| *Tmed5* | let-7a/b/c/d/e/f/g/k | miRTarbase | Targetscan | Co-nodes | Endosomal, l | Transmembr | 2158.77026 | 0.70 | 5.4E-04 | 5.5E-03 |
| *Tmem37* | let-7a/b/c/d/k | | | Co-nodes | Membrane p | Transmembr | 736.689845 | 1.22 | 5.5E-04 | 5.5E-03 |
| *Tmem41a* | let-7a/b/c/d/f | | Targetscan | Co-nodes | Membrane p | Transmembr | 272.674642 | 0.88 | 2.4E-03 | 1.6E-02 |
| *Tmem50b* | let-7a/b/c/d/f | | | Co-nodes | Membrane p | Transmembr | 2856.1835 | 0.83 | 4.4E-03 | 2.6E-02 |
| *Tmsb10* | let-7a/b/c/f/g/i/k | | | Co-nodes | Cytoskeleton | Thymosin be | 1747.41111 | 0.54 | 9.6E-03 | 4.7E-02 |
| *Tmsb4x* | let-7a/b/c/e/f/g/k | | | Co-nodes | Cytoskeleton | Thymosin be | 58895.7144 | 0.87 | 6.5E-04 | 6.2E-03 |
| *Tpcn1* | let-7a/b/c/d/f/k | | | Ion channels | Voltage gate | Two pore seg | 702.546931 | 0.68 | 2.9E-03 | 1.9E-02 |
| *Tpm1* | let-7a/b/c/d/f/k | | | Co-nodes | Cytoskeleton | Tropomyosin | 11974.3513 | 0.65 | 3.2E-05 | 5.4E-04 |
| *Tpm4* | let-7a/b/c/d/e/f/k | | | Co-nodes | Cytoskeleton | Tropomyosin | 13140.9284 | 0.51 | 1.5E-03 | 1.2E-02 |
| *Trabd2b* | let-7b/c/k | | | Co-nodes | Other co-nod | TraB domain | 37.0146277 | 2.82 | 8.0E-04 | 7.3E-03 |
| *Traf4* | let-7b/d | | Targetscan | Enzymes | E3 ubiquitin | TNF receptor | 3025.79617 | 0.85 | 2.6E-05 | 4.5E-04 |
| *Trim2* | let-7a/b/c/k | | | Enzymes | E3 ubiquitin | Tripartite mo | 2768.28073 | 0.62 | 1.1E-03 | 9.3E-03 |
| *Ttc28* | let-7b/c/d//k | | | Co-nodes | Other co-nod | Tetratricoper | 75.5267655 | 1.81 | 4.8E-04 | 4.9E-03 |
| *Tubb5* | let-7a/b/c/d/e/f/i/k | | | Co-nodes | Cytoskeleton | Tubulin | 22395.4671 | 0.72 | 8.2E-05 | 1.2E-03 |
| *Txn2* | let-7a/b/c/d/f | | Targetscan | Co-nodes | Redox protei | Thioredoxin | 1936.90202 | 0.54 | 3.8E-03 | 2.4E-02 |
| *Ube2g2* | let-7a/b/c | | Targetscan | Enzymes | E2 ubiquitin | Ubiquitin con | 1440.97513 | 0.56 | 3.5E-03 | 2.2E-02 |
| *Ube2j1* | let-7a/b | | Targetscan | Enzymes | E2 ubiquitin | Ubiquitin con | 1654.89381 | 0.76 | 2.4E-05 | 4.2E-04 |
| *Ubxn8* | let-7a/b/c/d/e/k | | | Co-nodes | UBX domain | UBX domain | 429.232946 | 0.87 | 5.8E-04 | 5.7E-03 |
| *Uhrf1* | | miRTarbase | | Enzymes | E3 ubiquitin | Ubiquitin-like | 3372.55519 | 0.87 | 1.1E-03 | 8.9E-03 |
| *Vapa* | let-7c/d/f | | | Co-nodes | Interacting p | VAMP associ | 3283.31866 | 0.50 | 4.4E-03 | 2.6E-02 |
| *Vcam1* | let-7a/c | | | Receptors | Ligands | Vascular cell | 440.124293 | 1.89 | 1.9E-09 | 1.1E-07 |
| *Vps25* | let-7a/b/c/d/f/i/k | | Targetscan | Co-nodes | Endosomal, l | Vacuolar pro | 1769.21856 | 0.59 | 1.3E-03 | 1.0E-02 |
| *Wdr46* | | miRTarbase | | Co-nodes | WD repeat p | WD repeat d | 614.235096 | 0.79 | 4.9E-03 | 2.9E-02 |
| *Wipf1* | let-7a/b/d/k | | | Co-nodes | Interacting p | WAS/WASL | 102.78755 | 1.84 | 2.3E-03 | 1.6E-02 |
| *Yeats4* | let-7a/b/c/d/e/f/k | | Targetscan | Co-nodes | Other co-nod | YEATS doma | 1097.72132 | 0.66 | 1.5E-03 | 1.2E-02 |
| *Ywhaz* | let-7a/b/c/d/e/f/k | miRTarbase | | Enzymes | Enzyme regu | Tyrosine-mo | 15442.8688 | 0.58 | 2.6E-04 | 3.0E-03 |
| *Zfp169* | let-7b/f/k | | Targetscan | Co-nodes | Zinc finger p | ZNF zinc fing | 1186.26611 | 2.24 | 1.3E-23 | 8.2E-21 |
| *Zfp568* | let-7a/b/c/f | | Targetscan | NA | NA | NA | 1332.61479 | 1.18 | 1.2E-04 | 1.6E-03 |
| *Zfp697* | let-7a/b/c/d/e/f/k | | Targetscan | NA | NA | NA | 144.700561 | 2.06 | 1.3E-08 | 6.1E-07 |
