## Supplementary Data 3 for "*Let-7* restrains an oncogenic circuit in AT2 cells to prevent fibrogenic cell intermediates in pulmonary fibrosis"

### Supplementary Data 3: *let-7* AT2 Cell Targetome Gene Pathway Annotation

| Grouped Terms | Term Names | ORA FDR p value | Gene Symbols |
|---|---|---|---|
| Stemness, Cell Cycle, Growth, Proliferation | HALLMARK_G2-M CHECKPOINT | 1.13E-05 | Prim2, Smarcc1, Nolc1, Hif1A, Aurkb, Cdc25B, Ccnb2, Marcks, Espl1, Myc, Rad21, Srsf2, E2F1, E2F2, Dmd, Hmgn2, Ezh2 |
| | HALLMARK_E2F TARGETS | 1.23E-04 | Prim2, Cbx5, Rrm2, Ak2, Nolc1, Nap1L1, Aurkb, Cdc25B, Ccnb2, Espl1, Myc, Rad21, Srsf2, Tk1, Ezh2 |
| | WIKIPATHWAY_MECHANISMS ASSOCIATED PLURIPOTENCY | 3.46E-02 | Smarcc1, Uhrf1, Gata6, Tcf7l1, Casp3, Ezh2, Acvr1b, Hif1a, Parp1, Myc, Pim1, Ipo9 |
| | GOBP_MTOTIC CELL CYCLE PROCESS | 2.65E-06 | Aven, Ccng1, Aurkb, Rad21, Anxa1, Cdc25b, Knstrn, E2f1, Ccnb2, Nme6, Psme3, E2f2, Actb, Smarcc1, Acvr1b, Rhoa, Rrm2, Myh10, Edn1, Bach1, Nek6, Rrm1, Flna, Snx18, Haspin, Espl1, Nup62, E2f6, Senp2, Iqgap3, Pdpn, Ezh2, Dna2, Cfl1, Myc, Rbm38 |
| | GOBP_POS. REG. CELL POPULATION PROLIFERATION | 1.95E-09 | Syk, Anxa1, Cd38, Fgf1, Actb, Smarcc1, Col18a1, Crip2, Bcl2l1, Edn1, Myc, Fam98a, Gata6, Nolc1, Flt4, Rrm2, Itgb3, Tgfb3, Osmr, Pim1, Epha4, Itga4, E2f1, Nras, Vcam1, Ezh2, Kras, Acer3, Cd276, Lrg1, Fermt2, Ppp1r16b, Dmd, Ptafr, Fxn, Dot1l, Hif1a, Foxp2, Rac2, Aspm, Hipk2, Ntn1, Iqgap3, Itgal, Ilk, Ctsh, Nap1l1, Gnq5, Clu, Gnas, Flna, Adk, Csf2rb |
| | GOBP_EPITHELIAL CELL PROLIFERATION | 0.009746511 | Fgf1, Col18a1, Bcl2l1, Uhrf1, Sparc, Flt4, Itgb3, Dab2, Itga4, Nras, Loxl2, Lrg1, Ppp1r16b, Col4a3, Stk4, Myc, Foxp2, Lipa, Iqgap3, Adk |
| | GOBP_GROWTH | 7.17E-07 | Anxa1, Nme6, Sema7a, Ntn1, Edn1, Epb41l5, Acvr1b, Eif4g2, Yeats4, Dab2, Sema5a, Pim1, Itga4, Ncbp1, Cd38, Adipor2, Fgf1, Rgs4, Dmd, Taf9b, Fxn, Gata6, Stk4, Gpd2, Csf2rb, Bcl2l1, Myh10, Hif1a, Rdh10, Nrp2, Zeb2, Gnas, Ezh2, Gpat4, Ccnb2, Aspm, Clic4, Agrn, Opa3, Zfp568, Rhoa, Flt4, Ilk, Acsl4, Picalm, Cfl1 |
| Nutrient Sensing & Metabolism | KEGG_PI3K-AKT SIGNALING PATHWAY | 2.38E-04 | Bcl2l1, Col4a3, Col6a1, Col6a2, Col1a1, Fgf1, Flt4, Gng5, Itga4, Itgb3, Itgb6, Kras, Lamb1, Myc, Nras, Osmr, Syk, Ywhaz, Lamc1, Eif4e2 |
| | WIKIPATHWAY_FOCAL ADHESION PI3K-AKT-MTOR PATHWAY | 0.001740586 | Col6a2, Col5a2, Itga4, Itgal, Col1a1, Eif4e2, Osmr, Lamc1, Hif1a, Fgf1, Nras, Flt4, Gng5, Kras, Lamb1, Itgb3, Itgb6 |
| | HALLMARK_PI3K AKT MTORC SIGNALING | 0.003954066 | Map2K3, Ralb, Arhgdia, Prkar2A, Cfl1, E2F1, Sla, Pfn1 |
| | GOBP_CARBODYDRATE DERIVATIVE BIO. PROCESS | 0.00438566 | Acsl1, Nudt4, Rrm2, Dcakd, Tk1, Ctbs, Dera, Rrm1, Gla, Acsl4, Gnptab, Man2a2, Hs2st1, Pigs, Dpy19l1, Fkrp, Bgn, Adk, Spock2, Hif1a, Myc, Rab1b, Arl2, Parp1, Gpd2, Ube2j1, Ak2, Tmem165, Gpat4, Nme6, Ggta1, Pign, Mtch2, Bcl2l1, Rhoa |
| | KEGG_LYSOSOME | 0.006655361 | Gla, Ap1m1, Ap1s1, Ctsb, Ctsh, Ctss, Lipa, Gnptab, Galns |
| | HALLMARK_MTORC1 SIGNALING | 0.006108741 | Map2K3, Gclc, Rrm2, Psme3, Gsr, Ccng1, Sla, Etf1, Glrx, Adipor2, Gla |
| Extracellular Matrix, Cytoskeleton, Motility, Cell Polarity | HALLMARK_EPITHELIAL MESENCHYMAL TRANSITION | 9.64E-12 | Sparc, Wipf1, Eln, Itgb3, Serpine1, Col12A1, Lamc1, Loxl2, Rgs4, Flna, Edil3, Tgm2, Vcam1, Tpm4, Tpm1, Bgn, Plaur, Col1A1, Acta2, Dab2, Lox, Col6A2, Adam12, Col5A2, Fermt2 |
| | GOBP_CELL ADHESION | 1.47E-08 | Lamb1, Itgb3, Syk, Nectin3, Anxa1, Fermt3, Itgb6, Vcam1, Pdpn, Plxnd1, Olr1, Itgal, Ilk, Lpp, Plxnb2, Fermt2, Actb, Smarcc1, Col1a1, Rhoa, Il1rap, Vamp3, Edil3, Cd276, Npnt, Spock2, Col6a1, Col18a1, Col6a2, Myh10, Dab2, Emilin2, Epha4, Lamc1, Itga4, Zyx, Msn, Flna, Mcam, Col12a1, Rac2, Lrg1, Serpine1, Plaur, Adam12, Ptafr, Ap1ar, Col4a3, Ntn1, Faf1, Stk4, Epb41l5, Casp3, Tgm2, Dmd, Sdc3, Gnas, Tpm1, Adk, Cfl1, Marcks, Arl2 |
| | KEGG_FOCAL ADHESION | 2.60E-05 | Actb, Rhoa, Col4a3, Col6a1, Col6a2, Col1a1, Flt4, Ilk, Itga4, Itgb3, Itgb6, Lamb1, Flna, Rac2, Rap1b, Lamc1, Zyx |
| | KEGG_ECM-RECEPTOR INTERACTION | 3.70E-05 | Npnt, Agrn, Col4a3, Col6a1, Col6a2, Col1a1, Itga4, Itgb3, Itgb6, Lamb1, Lamc1 |
| | HALLMARK_TGF-beta SIGNALING | 2.51E-03 | Fkbp1A, Slc20A1, Ifngr2, Serpine1, Rhoa, Hipk2 |
| Cell Survival, Senescence, & Death | GOBP REGULATION CELL DEATH | 2.20E-08 | Bcl2l1, Faf1, Clu, Tnfrsf10b, Anxa1, Mtch2, Tnfrsf1b, Casp3, Hspb6, Yeats4, Actb, Col18a1, Angptl4, Myc, Apbb2, Serpinb9, Plagl2, Fcer1g, Aven, Gata6, Sod2, Traf4, Stk4, Cdc34, Ccng1, Flt4, Itgb3, Aurkb, Hif1a, Tgfb3, Edn1, Syk, Arf4, Dab2, Pim1, Emilin2, Parp1, Itga4, E2f1, Cd38, Rad18, Acer3, Flna, Gclc, Serpine1, Fermt2, Tgm2, Rnf122, Picalm, Slc25a24, Plaur, Taf9b, Fxn, Psme3, Nup62, Ywhaz, Kras, Ctsh, Rhoa, Mical1, Lox, Trim2, Pdpn, Plxnd1, Agrn, Hipk2, Glrx, Ctsb, Sp100, Lcn2, Ilk |
| | GOBP_APOPTOTIC PROCESS | 5.01E-07 | Bcl2l1, Clu, Tnfrsf10b, Anxa1, Mtch2, Tnfrsf1b, Casp3, Hspb6, Hipk2, Yeats4, Actb, Col18a1, Angptl4, Faf1, E2f2, Myc, Lcn2, E2f1, Apbb2, Serpinb9, Plagl2, Fcer1g, Acvr1b, Aven, Ralb, Gata6, Sod2, Traf4, Stk4, Cdc34, Ccng1, Flt4, Bcap29, Aurkb, Ntn1, Hif1a, Tgfb3, Edn1, Arf4, Dab2, Rad21, Pim1, Emilin2, Parp1, Nek6, Itga4, Cd38, Flna, Gclc, Ctsh, Serpine1, Tgm2, Rnf122, Plaur, Taf9b, Fxn, Psme3, Col4a3, Nup62, Kras, Rhoa, Mical1, Lox, Trim2, Pdpn, Plxnd1, Agrn, Marcks, Sp100, Ilk, Lsp1 |
| | KEGG_P53 SIGNALING PATHWAY | 0.000208358 | Bcl2l1, Casp3, Ccnb2, Ccng1, Serpine1, Rrm2, Tnfrsf10b, Zmat3, Sesn3 |
| | KEGG_CELLULAR SENESCENCE | 0.009155925 | Ccnb2, E2f1, Hipk2, Kras, Myc, Nras, Serpine1, Tgfb3, E2f2, Map2k3 |
| Inflammation, Immunity | HALLMARK_INFLAMMATORY RESPONSE | 4.14E-04 | Msr1, Edn1, Ifngr2, Itgb3, Serpine1, Ptafr, Plaur, Osmr, Tnfrsf1B, Acvr1B, Hif1A, Myc, Pdpn, Olr1 |
| | HALLMARK_IL-6/JAK/STAT3 SIGNALING | 4.14E-04 | Itga4, Itgb3, Ifngr2, Pim1, Cd38, Csf2Rb, Osmr, Tnfrsf1B, Acvr1B |
| | HALLMARK_TNFalpha SIGNALING VIA NFKB | 0.006108741 | Ehd1, Map2K3, Edn1, Marcks, Myc, Ifngr2, Serpine1, Slc16A6, Plaur, Olr1, Sod2 |
| | REACTOME_IMMUNE SYSTEM | 8.15E-05 | Col1a1, Tubb5, Ap1m1, Pafah1b2, Ap1s1, Eif4g2, Bcl2l1, Rhoa, Ube2g2, Surf4, Galns, Psma2, Timp2, Myo1c, Cyb5r3, Map2k3, Grb10, Cdc34, Syk, Ctsb, Clu, Osmr, Ywhaz, Sla, Il1rap, Ifngr2, Vapa, Psmd1, Eif4e2, Lcn2, Itga4, S100a11, Vcam1, Ube2j1, Tnfrsf1b, Fgr, Vamp3, Actb, Olr1, Dera, Kras, Itgal, Gla, Casp3, Tmem30a, Ctsh, Fkbp1a, Rac2, Ctss, Cd53, Nhlrc3, Plaur, Rap1b, Ptafr, Nfam1, Fcer1g, Eif4a1, Csf2rb, Wipf1, Psme3 |
| Stress Responses | GOBP_CELLULAR RESPONSE STRESS | 1.34E-05 | Bcl2l1, Ube2g2, Faf1, Rnf5, Traf4, Hif1a, Rad21, Anxa1, Nucks1, Ube2j1, Tnfrsf1b, Rad18, Gsr, Sesn3, Hif1an, Nrep, Hipk2, Dot1l, Yeats4, Actb, Rrm1, Smarcc1, Myc, Ezh2, Casp3, Ino80c, Marcks, Nup62, Uhrf1, Ralb, Gata6, Sod2, Cbx5, Gorasp2, Map2k3, Ccng1, Flt4, Edn1, Clu, Bach1, Epha4, Parp1, E2f1, Tpm1, Prr5l, Fgf1, Dna2, Slc25a24, Rap2a, Ubxn8, Fxn, Mtch2, Flna, Rhoa, Glrx, Sp100, Lcn2, Rbm38, Kras, Ilk, Slc35a4, Serpinb6b, Cfl1, Tnfrsf10b, Angptl4 |
| | GOBP_RESPONSE OXIDATIVE STRESS | 0.001204245 | Anxa1, Gsr, Ezh2, Col1a1, Txn2, Sod2, Hif1a, Edn1, Parp1, Cd38, Rbpms, Casp3, Sesn3, Gclc, Tpm1, Prr5l, Slc25a24, Fxn, Glrx, Lcn2, Cfl1 |
| Cell Differentiation & Development | GOBP_EPITHELIUM_DEVELOPMENT | 5.58E-09 | Col18a1, Gata6, Anxa1, Casp3, Krt80, Ppp1r16b, Epb41l5, Plxnd1, Kras, Ilk, Tubb5, Rhoa, Marveld2, Ctsb, Dab2, Tagln2, Actb, Msn, Flna, Acta2, Fgf1, Lrg1, Clic4, Serpine1, Fermt2, Kazn, Rap1b, Stk4, Pfn1, Ntn1, Epha4, Spint1, Foxp2, Acvr1b, Hif1a, Edn1, Myc, Clic5, Rdh10, Zeb2, Car2, Iqgap3, Pdpn, Ezh2, Gpat4, Ctsh, Plxnb2, Tgm2, Hs2st1, Npnt, Dmd, Cfl1, Marcks, Zfp568, Lcn2, Gnas, Plaur, Vcam1 |
| | GOBP_STEM_CELL_DIFFERENTIATION | 0.045017855 | Sema7a, Edn1, Nolc1, Gata6, Hif1a, Rdh10, Nrp2, Zeb2, Mtch2, Ezh2, Cfl1 |
| | GO BP_EPITHELIAL CELL DIFFERENTIATION | 0.000771462 | Col18a1, Gata6, Anxa1, Casp3, Krt80, Ppp1r16b, Epb41l5, Kras, Tubb5, Rhoa, Marveld2, Ctsb, Dab2, Tagln2, Msn, Acta2, Clic4, Serpine1, Kazn, Rap1b, Stk4, Hif1a, Clic5, Zeb2, Pdpn, Ezh2, Gpat4, Dmd, Flna, Plaur |
| Lung Cancer | KEGG_MICRORNAS IN CANCER | 2.60E-05 | Rhoa, Casp3, Ccng1, Cdc25b, E2f1, Ezh2, Itgb3, Kras, Marcks, Myc, Nras, Pim1, Tpm1, Zeb2, E2f2 |
| | KEGG_PATHWAYS IN CANCER | 0.000208358 | Rhoa, Bcl2l1, Casp3, Col4a3, Csf2rb, E2f1, Edn1, Fgf1, Flt4, Gnas, Gng5, Hif1a, Ifngr2, Kras, Lamb1, Myc, Nras, Pim1, Rac2, Tcf7l1, Tgfb3, Traf4, Lamc1, E2f2, Stk4, Ralb |
| | KEGG_SMALL CELL LUNG CANCER | 7.84E-04 | Bcl2l1, Casp3, Col4a3, E2f1, Lamb1, Myc, Traf4, Lamc1, E2f2 |
