## Supplementary Data 4 for "*Let-7* restrains an oncogenic circuit in AT2 cells to prevent fibrogenic cell intermediates in pulmonary fibrosis"

### Supplementary Data 4: AT2 Cell ChIP Seq Regulatory Network Analysis

| NODE | Category | Class | Family | Symbol | Aliases | PVAL sorted_AT2_UP | PVAL organoids_UP | PVAL sorted_AT2_DOWN | PVAL organoids_DOWN |
|---|---|---|---|---|---|---|---|---|---|
| E2f4 | Transcriptio | E2F/FOX | E2F | E2f4 | E2F-4 | 1.08E-135 | 1.7162E-88 | | |
| Sirt1 | Enzymes | Deacetylas | Sirtuins (S | Sirt1 | SIR2L1 | 6.107E-80 | 2.3546E-55 | | |
| Lin9 | Co-nodes | Transcripti | Lin-9 DRE/ | Lin9 | TGS | 1.591E-56 | 9.0776E-30 | | |
| E2f3 | Transcriptio | E2F/FOX | E2F | E2f3 | | 6.184E-39 | 7.2724E-51 | | |
| Hdac1 | Enzymes | Deacetylas | Histone de | Hdac1 | HD1, GON | 4.91E-33 | 1.3297E-48 | | |
| Sap130 | Co-nodes | Transcripti | Sin3A asso | Sap130 | FLJ12761 | 3.234E-30 | 8.8924E-83 | | |
| Stat1 | Transcriptio | STAT dom | STAT facto | Stat1 | STAT91, IS | 2.369E-28 | 9.4215E-05 | 0.00832 | 2.4E-18 |
| Spi1 | Transcriptio | Tryptophar | Spi-like | Spi1 | PU.1, SPI-/ | 3.059E-26 | 4.1095E-15 | 0.00342 | 1E-11 |
| Sin3a | Co-nodes | Transcripti | SIN3 trans | Sin3a | KIAA0700, | 2.103E-23 | 2.5689E-35 | | |
| Stat5a | Transcriptio | STAT dom | STAT facto | Stat5a | MGF | 2.139E-22 | 7.4735E-13 | 1.5E-06 | 2.8E-18 |
| Irf8 | Transcriptio | Tryptophar | Interferon | Irf8 | IRF-8, ICSE | 1.33E-20 | 3.0839E-07 | 0.02411 | 9.5E-10 |
| Atf3 | Transcriptio | BZIP facto | ATF-2-like | Atf3 | | 6.56E-19 | 0.0001446 | 0.01646 | 2.9E-10 |
| Tal1 | Transcriptio | BHLH facto | Tal/HEN-li | Tal1 | SCL, bHLHa | 1.884E-18 | 7.005E-09 | 1.6E-06 | 1E-13 |
| Irf4 | Transcriptio | Tryptophar | Interferon | Irf4 | LSIRF | 1.805E-16 | 0.0002233 | 0.001 | 2.1E-09 |
| Stat3 | Transcriptio | STAT dom | STAT facto | Stat3 | APRF | 8.475E-16 | 1.0354E-07 | 3.2E-05 | 1E-15 |
| Jun | Transcriptio | BZIP facto | Jun factor | Jun | c-Jun, AP-1 | 1.998E-15 | 2.2928E-14 | 0.00101 | 1.7E-12 |
| Stat6 | Transcriptio | STAT dom | STAT facto | Stat6 | D12S1644, | 3.491E-15 | 4.1288E-07 | | 0.00177 |
| Kat5 | Enzymes | Acetyltran | Lysine ace | Kat5 | TIP60, PLIF | 6.681E-15 | 1.8454E-44 | | |
| Ep300 | Enzymes | Acetyltran | CBP/p300 | Ep300 | p300, KAT | 9.019E-15 | 4.4782E-09 | 6.3E-09 | 4.3E-19 |
| Rad21 | Co-nodes | Cohesin co | Cell cycle ( | Rad21 | KIAA0078, | 1.137E-14 | 7.7806E-09 | 0.00079 | 1.5E-10 |
| Cebpd | Transcriptio | BZIP facto | C/EBP | Cebpd | CRP3, CELF | 2.016E-14 | 0.01442195 | | 2.7E-08 |
| Trp53 | Transcriptio | p53 domai | p53-relate | Trp53 | | 1.729E-13 | 0.0002418 | | 1.4E-06 |
| Cebpb | Transcriptio | BZIP facto | C/EBP | Cebpb | LAP, CRP2, | 2.528E-13 | 6.9812E-09 | 0.00639 | 2.9E-10 |
| Rxra | Receptors | Nuclear re | Retinoid X | Rxra | NR2B1 | 2.545E-13 | 6.2009E-17 | 4.1E-14 | 6.4E-18 |
| Nfkb1 | Transcriptio | Rel Homol | NF-kappaE | Nfkb1 | KBF1, p10 | 2.61E-13 | 0.00181776 | 0.0015 | 2.9E-11 |
| Nr1h2 | Receptors | Nuclear re | Liver X rec | Nr1h2 | NER, NER- | 4.191E-13 | 1.0513E-07 | 0.00095 | 0.00031 |
| Gata3 | Transcriptio | Other tran | Two zinc-f | Gata3 | HDR | 5.079E-13 | 2.1379E-08 | 0.00967 | 3.2E-07 |
| Irf9 | Transcriptio | Tryptophar | Interferon | Irf9 | | 7.421E-13 | | 0.0314 | 6.8E-13 |
| Stat2 | Transcriptio | STAT dom | STAT facto | Stat2 | STAT113 | 7.59E-13 | 1.1395E-05 | | 1.3E-12 |
| Ctcf | Transcriptio | C2H2 Zn fi | CTCF-like | Ctcf | FAP108, C | 7.76E-13 | 5.6753E-15 | | 0.00063 |
| Stag1 | Co-nodes | Cohesin co | Stromal ar | Stag1 | SA-1, SCC3 | 9.932E-13 | 1.2111E-05 | 0.0011 | 2.8E-10 |
| Stag2 | Co-nodes | Cohesin co | Stromal ar | Stag2 | SA-2, SCC3 | 1.145E-12 | 1.6534E-11 | 0.00046 | 6.6E-09 |
| Sin3b | Co-nodes | Transcripti | SIN3 trans | Sin3b | KIAA0700 | 1.394E-12 | 0.00045036 | | |
| Eomes | Transcriptio | T Box facto | TBrain-rela | Eomes | TBR2 | 2.104E-12 | 4.0243E-09 | 0.00113 | 1.6E-07 |
| Nfe2 | Transcriptio | BZIP facto | NF-E2-like | Nfe2 | NF-E2 | 3.66E-12 | 0.00117723 | | 0.00105 |
| Batf | Transcriptio | BZIP facto | B-ATF-rela | Batf | B-ATF, SF/ | 4.747E-12 | 7.7118E-05 | 0.00011 | 3.3E-11 |
| Nr4a1 | Receptors | Nuclear re | NGFBI-like | Nr4a1 | TR3, N10, | 4.75E-12 | 3.9058E-06 | 7.9E-05 | 6.7E-06 |
| Smc3 | Co-nodes | Cohesin co | Structural | Smc3 | HCAP, BAM | 5.934E-12 | 1.5235E-12 | 1.9E-07 | 1.4E-12 |
| Cebpe | Transcriptio | BZIP facto | C/EBP | Cebpe | CRP1 | 5.975E-12 | 0.02686498 | | 0.00074 |

| | | | | | | | | | |
|---|---|---|---|---|---|---|---|---|---|
| Dr1 | Co-nodes | Other co-r | Down-regu | Dr1 | NC2-BETA | 6.758E-12 | 2.9304E-32 | 5.3E-05 | 2.4E-06 |
| Runx1 | Transcriptio | Runt doma | Core-bindi | Runx1 | PEBP2A2, | 1.403E-11 | 5.3723E-22 | 0.0269 | 0.02658 |
| Smc1a | Co-nodes | Cohesin cc | Structural | Smc1a | DXS423E, | 1.701E-11 | 3.3655E-12 | 9.3E-09 | 2.1E-14 |
| Mafk | Transcriptio | BZIP facto | Small Maf | Mafk | P18, NFE2 | 2.819E-11 | 8.1182E-08 | 0.00461 | 1.9E-08 |
| Stat4 | Transcriptio | STAT dom | STAT facto | Stat4 | | 5.155E-11 | 0.00144905 | 0.00328 | 9.9E-10 |
| Hdac2 | Enzymes | Deacetylas | Histone de | Hdac2 | RPD3, YAF | 5.82E-11 | 1.3571E-28 | | |
| Junb | Transcriptio | BZIP facto | Jun factor | Junb | | 6.07E-11 | 9.2863E-06 | 0.01715 | 1.6E-09 |
| Batf3 | Transcriptio | BZIP facto | B-ATF-rela | Batf3 | JUNDM1, ! | 1.099E-10 | 0.00048396 | 0.03051 | 2.6E-05 |
| Runx2 | Transcriptio | Runt doma | Core-bindi | Runx2 | AML3, PEE | 1.579E-10 | 1.9863E-15 | 8E-05 | 4.2E-07 |
| Cebpa | Transcriptio | BZIP facto | C/EBP | Cebpa | C/EBP-alp | 1.634E-10 | 5.7265E-12 | 7.6E-05 | 3E-10 |
| Fosl1 | Transcriptio | BZIP facto | Fos factor | Fosl1 | fra-1 | 2.056E-10 | 3.6918E-13 | 0.00016 | 1E-22 |
| Nr3c1 | Receptors | Nuclear re | Glucocorti | Nr3c1 | GR | 2.425E-10 | 5.6974E-07 | 2E-10 | 3.8E-28 |
| Maff | Transcriptio | BZIP facto | Small Maf | Maff | hMafF | 3.595E-10 | 2.6977E-08 | 0.01301 | 4.8E-08 |
| Lmo2 | Co-nodes | LIM domai | LIM domai | Lmo2 | TTG2, RHO | 3.774E-10 | 8.2407E-09 | 0.01866 | 1E-12 |
| Mafb | Transcriptio | BZIP facto | Large Maf | Mafb | | 6.152E-10 | 2.6836E-05 | | 0.00189 |
| Gata4 | Transcriptio | Other tran | Two zinc-f | Gata4 | | 9.305E-10 | 2.1568E-27 | 0.00044 | 0.00343 |
| Tbx21 | Transcriptio | T Box facto | TBrain-rela | Tbx21 | TBLYM, T- | 1.374E-09 | 3.3848E-13 | 5.9E-06 | 0.00021 |
| Myc | Transcriptio | BHLH facto | Myc/Max | Myc | c-Myc, bHI | 1.548E-09 | 4.2513E-70 | | |
| Irf3 | Transcriptio | Tryptophai | Interferon | Irf3 | | 1.669E-09 | | 0.00187 | 1.7E-06 |
| Prdm16 | Co-nodes | PR domair | PR/SET do | Prdm16 | MEL1, PFN | 1.96E-09 | 2.2716E-12 | 1.1E-05 | 2.5E-12 |
| Srf | Transcriptio | MADS box | Respondei | Srf | MCM1 | 2.114E-09 | 4.4027E-18 | 0.0056 | 7.2E-09 |
| Fosl2 | Transcriptio | BZIP facto | Fos factor | Fosl2 | FRA2, FLJ2 | 2.428E-09 | 2.1644E-07 | 0.00038 | 1.7E-13 |
| Max | Transcriptio | BHLH facto | Myc/Max | Max | bHLHd4, b | 2.466E-09 | 3.8088E-52 | | |
| Bach2 | Transcriptio | BZIP facto | NF-E2-like | Bach2 | BTBD25 | 2.589E-09 | 1.1263E-08 | | 5.3E-06 |
| Gata1 | Transcriptio | Other tran | Two zinc-f | Gata1 | ERYF1, NF | 2.672E-09 | 1.1026E-05 | 2.3E-07 | 2.4E-06 |
| Klf5 | Transcriptio | C2H2 Zn fi | KrÂ▯uppel | Klf5 | IKLF, CKLF | 5.641E-09 | 4.0269E-27 | 0.00034 | 0.00013 |
| Meis1 | Transcriptio | Homeo do | MEIS | Meis1 | | 7.996E-09 | 6.0566E-11 | 4.4E-08 | 2.3E-14 |
| Rcor2 | Transcriptio | Tryptophai | REST core | Rcor2 | | 8.304E-09 | 4.0588E-13 | 0.00016 | 2.1E-07 |
| Mxi1 | Transcriptio | BHLH facto | Mad-like f | Mxi1 | MXD2, MA | 1.058E-08 | 9.4278E-34 | | |
| Hnf4a | Receptors | Nuclear re | Hepatocyt | Hnf4a | NR2A1, HN | 1.133E-08 | 5.0478E-05 | 2E-06 | 1.9E-11 |
| Gfi1b | Transcriptio | C2H2 Zn fi | GFI1 facto | Gfi1b | ZNF163B | 1.165E-08 | 3.9791E-08 | 0.0018 | 0.00078 |
| Gata2 | Transcriptio | Other tran | Two zinc-f | Gata2 | NFE1B | 1.338E-08 | 4.0968E-14 | 0.01756 | 2.7E-07 |
| Dmap1 | Transcriptio | Tryptophai | MIER-like | Dmap1 | DNMAP1, | 1.468E-08 | 1.3956E-29 | | |
| Stat5b | Transcriptio | STAT dom | STAT facto | Stat5b | | 1.593E-08 | 0.00133754 | 0.02443 | 1.6E-07 |
| Pparg | Receptors | Nuclear re | Peroxisom | Pparg | PPARG1, F | 1.641E-08 | 8.3799E-15 | 2E-05 | 1.1E-11 |
| Tead4 | Transcriptio | TEA doma | TEF-1-rela | Tead4 | TEF-3, TEF | 2.177E-08 | 1.593E-34 | 4.7E-08 | 0.00019 |
| Sox2 | Transcriptio | HMG dom | Group B | Sox2 | | 2.363E-08 | 3.6608E-17 | 0.00125 | 1.3E-10 |
| Jund | Transcriptio | BZIP facto | Jun factor | Jund | AP-1 | 2.371E-08 | 1.0003E-09 | 6.9E-06 | 5.9E-15 |
| Irf5 | Transcriptio | Tryptophai | Interferon | Irf5 | | 2.53E-08 | 0.03213732 | 0.01776 | 1E-06 |
| Trim33 | Enzymes | E3 ubiquiti | Tripartite | Trim33 | TIF1GAMN | 3.156E-08 | 6.5221E-07 | | 5E-07 |
| Lyl1 | Transcriptio | BHLH facto | Tal/HEN-li | Lyl1 | bHLHa18 | 3.911E-08 | 0.00092491 | 0.04903 | 4.2E-08 |
| Srebf1 | Transcriptio | BHLH facto | SREBP fac | Srebf1 | SREBP1, b | 4.056E-08 | | 2.3E-06 | 6.4E-15 |
| Tfap4 | Transcriptio | BHLH facto | AP-4 famil | Tfap4 | AP-4, bHLI | 4.249E-08 | 2.1109E-16 | 0.03721 | |
| Gfi1 | Transcriptio | C2H2 Zn fi | GFI1 facto | Gfi1 | GFI1A, GF | 4.369E-08 | 2.2631E-12 | | |
| Egr2 | Transcriptio | C2H2 Zn fi | Early grow | Egr2 | | 5.808E-08 | 3.619E-16 | | 2.8E-05 |
| Tcf4 | Transcriptio | BHLH facto | E2A-relate | Tcf4 | SEF2-1B, I | 6.363E-08 | 7.9932E-08 | 1.7E-07 | 1E-11 |
| Tcf3 | Transcriptio | BHLH facto | E2A-relate | Tcf3 | E2A, ITF1, | 7.82E-08 | 2.9401E-15 | 8.4E-07 | 4.9E-10 |
| Fos | Transcriptio | BZIP facto | Fos factor | Fos | c-fos, AP-1 | 7.826E-08 | 1.0564E-22 | 0.00014 | 4.9E-05 |
| Elf5 | Transcriptio | Tryptophai | EHF-like | Elf5 | | 1.609E-07 | 9.8186E-06 | 0.01735 | 6E-12 |

| | | | | | | | | | |
|---|---|---|---|---|---|---|---|---|---|
| Sp9 | Transcriptio | C2H2 Zn fi | Sp1-like | Sp9 | ZNF990 | 1.615E-07 | 1.8876E-10 | | 0.00057 |
| Ebf1 | Transcriptio | Rel Homol | Early B-Ce | Ebf1 | OLF1, COE | 2.202E-07 | 2.189E-07 | 0.01914 | 8E-05 |
| Cbfb | Transcriptio | Runt doma | Core-bindi | Cbfb | PEBP2B | 3.408E-07 | 5.4318E-10 | 0.00591 | 0.00414 |
| Rbbp4 | Co-nodes | Chromatin | RB binding | Rbbp4 | RbAp48, N | 5.037E-07 | 0.01380279 | | |
| Nkx2-1 | Transcriptio | Homeo do | NK-2.1 | Nkx2-1 | TTF-1, TTF | 5.318E-07 | 1.939E-14 | 1.3E-07 | 0.00076 |
| Med24 | Co-nodes | Transcripti | Mediator c | Med24 | TRAP100, | 7.928E-07 | 1.79E-06 | 7.9E-06 | 0.00027 |
| Ikzf1 | Transcriptio | C2H2 Zn fi | Ikaros | Ikzf1 | hIk-1, LyF- | 9.982E-07 | 6.0976E-10 | | 0.00148 |
| Trp63 | Transcriptio | p53 domai | p53-relate | Trp63 | | 1.225E-06 | 0.0001641 | 8.9E-08 | 2E-45 |
| E2f1 | Transcriptio | E2F/FOX | E2F | E2f1 | RBP3 | 1.229E-06 | 9.0767E-20 | | |
| Ncor1 | Transcriptio | Tryptopha | Nuclear re | Ncor1 | N-CoR, hC | 1.388E-06 | 1.0401E-06 | 3.4E-07 | 4.4E-15 |
| Sox9 | Transcriptio | HMG dom | Group E | Sox9 | SRA1 | 1.707E-06 | 1.8186E-17 | 3.5E-05 | 1.7E-10 |
| Foxf1 | Transcriptio | E2F/FOX | FOXF | Foxf1 | FREAC1 | 1.793E-06 | 5.6534E-07 | 4.1E-06 | 1.6E-09 |
| Tle3 | Co-nodes | Transcripti | Groucho/T | Tle3 | ESG, ESG3 | 1.833E-06 | 1.8504E-05 | 0.00184 | 1E-12 |
| Nanog | Transcriptio | Homeo do | Nanog hor | Nanog | FLJ12581, | 1.873E-06 | 2.8352E-14 | 0.00033 | 2.7E-07 |
| Zc3h11a | Co-nodes | Zinc finger | Zinc finger | Zc3h11a | KIAA0663 | 1.935E-06 | 0.02780929 | 0.00201 | 4.5E-07 |
| Hnf1b | Transcriptio | Homeo do | HNF1-like | Hnf1b | LFB3, VHN | 2.039E-06 | 7.6766E-08 | 5.3E-05 | 1.7E-05 |
| Pgr | Receptors | Nuclear re | Progester | Pgr | PR, NR3C3 | 2.372E-06 | 4.1589E-06 | 5.3E-07 | 8.2E-19 |
| Kmt2c | Enzymes | Methyltrai | Histone-ly | Kmt2c | MLL3, HAL | 2.417E-06 | 3.6591E-12 | 7.7E-06 | 3.3E-08 |
| Utf1 | Transcriptio | BZIP facto | UTF | Utf1 | | 2.453E-06 | 1.9566E-09 | 3.7E-05 | 4.2E-10 |
| Ncoa2 | Enzymes | Acetyltran | Nuclear re | Ncoa2 | TIF2, GRIP | 2.515E-06 | 9.6253E-14 | | |
| Mettl3 | Enzymes | Methyltrai | Methyltrai | Mettl3 | Spo8, M6A | 2.553E-06 | 5.2201E-15 | | 0.00129 |
| Mitf | Transcriptio | BHLH facto | TFE3-like t | Mitf | MI, bHLHe | 3.018E-06 | 4.9972E-07 | | |
| Kmt2d | Enzymes | Methyltrai | Histone-ly | Kmt2d | ALR, MLL4 | 3.034E-06 | 2.1557E-21 | 9.4E-05 | 0.0007 |
| Nipbl | Co-nodes | Other co-r | NIPBL coh | Nipbl | IDN3, DKF | 3.034E-06 | 5.7376E-27 | | |
| Snai1 | Transcriptio | C2H2 Zn fi | Snail-like | Snai1 | SNA, SLUG | 3.204E-06 | 4.7116E-16 | 4.3E-06 | 0.00348 |
| Esr1 | Receptors | Nuclear re | Estrogen r | Esr1 | NR3A1, Er | 3.707E-06 | 1.5186E-13 | 4.7E-06 | 3.8E-11 |
| Klf3 | Transcriptio | C2H2 Zn fi | KrÂ�uppel | Klf3 | BKLF | 3.877E-06 | 1.5293E-14 | 0.00173 | 0.01236 |
| Atf4 | Transcriptio | BZIP facto | ATF-4-rela | Atf4 | TAXREB67 | 3.881E-06 | 0.01739131 | | 1.6E-07 |
| Runx3 | Transcriptio | Runt doma | Core-bindi | Runx3 | AML2, PEB | 4.641E-06 | 3.3435E-16 | 0.00015 | |
| Esrrb | Receptors | Nuclear re | Estrogen-r | Esrrb | ERR2, ERR | 4.681E-06 | 6.3335E-11 | 0.00076 | 8.6E-08 |
| Gtf3c1 | Co-nodes | General tr | General tr | Gtf3c1 | TFIIIC220 | 6.485E-06 | 2.8348E-23 | | |
| Irf1 | Transcriptio | Tryptopha | Interferon | Irf1 | MAR | 6.917E-06 | | 0.00106 | 1.3E-07 |
| Tgif1 | Transcriptio | Homeo do | TGIF | Tgif1 | | 7.073E-06 | 5.2257E-20 | 0.00147 | 0.00052 |
| Hdac3 | Enzymes | Deacetylas | Histone de | Hdac3 | RPD3, HD3 | 7.09E-06 | 8.4159E-06 | 0.00017 | 3.2E-12 |
| Smarca4 | Co-nodes | Chromatin | SWI/SNF r | Smarca4 | hSNF2b, B | 7.336E-06 | 1.8954E-14 | 3.1E-08 | 1.4E-18 |
| Klf4 | Transcriptio | C2H2 Zn fi | KrÂ�uppel | Klf4 | EZF, GKLF | 8.281E-06 | 2.783E-22 | 0.00519 | 1.8E-08 |
| Arntl | Transcriptio | BHLH facto | Arnt-like fa | Arntl | MOP3, JAP | 9.5E-06 | 3.5354E-12 | 0.01785 | 7.3E-15 |
| Mef2c | Transcriptio | MADS box | Myocyte e | Mef2c | | 1.033E-05 | 8.0266E-11 | 8.9E-06 | 1.1E-09 |
| Tet2 | Enzymes | Dioxygena | Ten-eleven | Tet2 | FLJ20032 | 1.281E-05 | 7.2045E-08 | | 3.4E-07 |
| Nfib | Transcriptio | Other tran | Nuclear fa | Nfib | NFI-RED, N | 1.382E-05 | 1.302E-11 | 0.00056 | 2.9E-15 |
| Maf | Transcriptio | BZIP facto | Large Maf | Maf | c-MAF | 1.439E-05 | 0.02262513 | 0.03496 | 6.3E-08 |
| Pou5f1 | Transcriptio | Homeo do | Oct-3/4-lil | Pou5f1 | OCT3, Oct | 1.646E-05 | 9.9539E-13 | 1.2E-05 | 1.2E-12 |
| Prdm13 | Co-nodes | PR domair | PR/SET do | Prdm13 | PFM10 | 1.731E-05 | 8.5125E-16 | | 0.00591 |
| Ahr | Transcriptio | BHLH facto | Ahr-like fa | Ahr | bHLHe76 | 1.826E-05 | 0.00122279 | 1.6E-05 | 8.2E-08 |
| Aire | Transcriptio | SAND dom | AIRE | Aire | PGA1, APS | 1.908E-05 | 2.6201E-17 | | 0.04387 |
| Rbpj | Transcriptio | Rel Homol | M | Rbpj | SUH, IGKJI | 1.957E-05 | 4.6352E-12 | | 9.2E-05 |
| Foxa2 | Transcriptio | E2F/FOX | FOXA | Foxa2 | | 1.987E-05 | 5.8194E-14 | 5.8E-20 | 3.3E-20 |
| Foxa1 | Transcriptio | E2F/FOX | FOXA | Foxa1 | | 2.127E-05 | 2.044E-10 | 1.3E-23 | 1.8E-16 |
| Sox6 | Transcriptio | HMG dom | Group D | Sox6 | | 2.156E-05 | 1.9236E-12 | | 6.1E-14 |

| | | | | | | | | | |
|---|---|---|---|---|---|---|---|---|---|
| Bach1 | Transcriptio | BZIP facto | NF-E2-like | Bach1 | BACH-1, B | 2.207E-05 | 2.5881E-05 | | 0.0185 |
| Tead1 | Transcriptio | TEA doma | TEF-1-rela | Tead1 | TEF-1 | 2.3E-05 | 5.0756E-18 | 7.2E-07 | 7.5E-07 |
| Satb1 | Transcriptio | Homeo do | SATB | Satb1 | | 2.318E-05 | 1.099E-10 | 0.03594 | 5.2E-05 |
| Paxip1 | Co-nodes | Interacting | PAX intera | Paxip1 | CAGF29, C | 2.83E-05 | 2.9417E-08 | | |
| Nr1d1 | Receptors | Nuclear re | Rev-Erb re | Nr1d1 | ear-1, hRe | 2.957E-05 | 1.3847E-12 | 3.1E-06 | 3.1E-14 |
| Ttf1 | Co-nodes | Nucleopro | Transcripti | Ttf1 | | 3.026E-05 | 3.6445E-20 | 4.9E-21 | 3.9E-06 |
| Nfia | Transcriptio | Other tran | Nuclear fa | Nfia | NFI-L, KIA | 3.186E-05 | 0.0039399 | 0.03974 | 3E-08 |
| Gata6 | Transcriptio | Other tran | Two zinc-f | Gata6 | | 4.722E-05 | 0.00145181 | | 0.03049 |
| Yap1 | Co-nodes | Transcripti | Yes1 asso | Yap1 | YAP65 | 4.741E-05 | 1.1746E-16 | 7.3E-05 | 0.0015 |
| Crebbp | Enzymes | Acetyltran | CBP/p300 | Crebbp | RTS, CBP, | 5.162E-05 | 8.1114E-13 | 6.2E-05 | 1.5E-12 |
| Myb | Transcriptio | Tryptopha | Myb-like | Myb | c-myb | 5.341E-05 | 2.9685E-07 | | 0.02457 |
| Wiz | Transcriptio | C2H2 Zn fi | ZNF37A-li | Wiz | ZNF803 | 5.363E-05 | 7.6022E-20 | 0.00238 | 0.03195 |
| Clock | Transcriptio | BHLH facto | Arnt-like f | Clock | KIAA0334, | 5.813E-05 | 1.9965E-09 | 0.00043 | 7.4E-14 |
| Bcl11b | Transcriptio | C2H2 Zn fi | B-cell lym | Bcl11b | CTIP-2, CT | 6.025E-05 | 3.5857E-18 | 0.0424 | |
| Ctcfl | Transcriptio | C2H2 Zn fi | CTCF-like | Ctcfl | dJ579F20. | 6.079E-05 | 5.9437E-22 | 7.3E-05 | 0.00033 |
| Kdm4c | Enzymes | Demethyla | Histone-H3 | Kdm4c | GASC1, KI | 6.249E-05 | 1.2827E-17 | 0.00124 | 0.03551 |
| Nfyb | Transcriptio | Heteromer | Heteromer | Nfyb | CBF-A, HA | 6.371E-05 | 5.9907E-12 | | |
| Olig2 | Transcriptio | BHLH facto | Neurogeni | Olig2 | RACK17, C | 6.411E-05 | 5.508E-15 | 3E-10 | 2E-11 |
| Mta2 | Transcriptio | Other tran | Single GA | Mta2 | MTA1-L1 | 6.591E-05 | 1.9125E-15 | | 0.03755 |
| Erf | Transcriptio | Tryptopha | Ets-like | Erf | PE-2, PE2 | 6.933E-05 | 2.7117E-32 | | |
| Foxp3 | Transcriptio | E2F/FOX | FOXP | Foxp3 | JM2, XPID, | 7.248E-05 | 2.3801E-06 | 1E-05 | 3E-09 |
| Ppara | Receptors | Nuclear re | Peroxisom | Ppara | hPPAR, NF | 7.293E-05 | 2.416E-06 | 8.7E-05 | 8.8E-11 |
| Insm1 | Transcriptio | C2H2 Zn fi | Insulinoma | Insm1 | IA-1, IA1 | 7.981E-05 | 1.6382E-09 | | |
| Nr1h3 | Receptors | Nuclear re | Liver X rec | Nr1h3 | LXR-a, RLD | 8.032E-05 | 4.4701E-11 | 8.9E-06 | 0.00066 |
| Qk | Co-nodes | Other co-r | Quaking | Qk | | 8.55E-05 | 2.1038E-20 | | |
| Mybl1 | Transcriptio | Tryptopha | Myb-like | Mybl1 | AMYB, A-r | 8.64E-05 | 0.00056344 | | |
| Ppard | Receptors | Nuclear re | Peroxisom | Ppard | NUC1, NU | 8.736E-05 | 5.7941E-09 | | |
| Sumo2 | Co-nodes | Ubiquitin & | Small ubic | Sumo2 | SMT3B | 8.944E-05 | 6.7146E-07 | 0.00269 | 5.4E-05 |
| Irf2bp2 | Co-nodes | Transcripti | Interferon | Irf2bp2 | IRF-2BP2 | 9.2E-05 | 0.00342276 | 0.00027 | 0.00537 |
| Sumo1 | Co-nodes | Ubiquitin & | Small ubic | Sumo1 | PIC1, GMP | 9.354E-05 | 2.4204E-12 | 0.02718 | 0.00726 |
| Creb1 | Transcriptio | BZIP facto | CREB-like | Creb1 | | 9.41E-05 | 3.5218E-24 | | |
| Fli1 | Transcriptio | Tryptopha | Ets-like | Fli1 | SIC-1, EWS | 9.532E-05 | 2.0803E-24 | | |
| Hnf4g | Receptors | Nuclear re | Hepatocyt | Hnf4g | NR2A2 | 9.545E-05 | 0.00017431 | 0.0005 | |
| Myog | Transcriptio | BHLH facto | Myogenic | Myog | bHLHc3 | 9.561E-05 | 8.0206E-13 | 0.00069 | 8.3E-12 |
| Dpy30 | Enzymes | Regulatory | Dpy-30 his | Dpy30 | Saf19, HDI | 9.917E-05 | 2.3909E-22 | | 0.04801 |
| Bcl6 | Transcriptio | C2H2 Zn fi | BCL6 facto | Bcl6 | ZBTB27, L | 0.0001173 | 1.1124E-09 | 9.8E-06 | 1.2E-07 |
| Nacc1 | Co-nodes | BTB doma | Nucleus ac | Nacc1 | NAC1, NA | 0.0001347 | 8.9906E-23 | | |
| Jmjd6 | Enzymes | Demethyla | Jumonji do | Jmjd6 | PTDSR1, K | 0.0001408 | 0.00281161 | | 8.5E-05 |
| Prox1 | Transcriptio | Homeo do | Prospero h | Prox1 | | 0.0001535 | 0.00190499 | 0.00038 | 3.1E-07 |
| Hnf1a | Transcriptio | Homeo do | HNF1-like | Hnf1a | HNF1, LFB | 0.0001571 | | 0.00654 | 0.00336 |
| Ebf2 | Transcriptio | Rel Homol | Early B-Ce | Ebf2 | FLJ11500, | 0.0001639 | 1.3072E-12 | 2.8E-08 | 3.9E-05 |
| Chd1 | Enzymes | Helicases | Chromodo | Chd1 | | 0.0001743 | 2.0698E-10 | 0.00694 | |
| Esrra | Receptors | Nuclear re | Estrogen-r | Esrra | ERR1, ERR | 0.0001894 | 1.1488E-09 | 0.0039 | 3.4E-06 |
| Ncor2 | Transcriptio | Tryptopha | Nuclear re | Ncor2 | SMRT, SM | 0.0001918 | 9.7752E-23 | 4.4E-05 | 9.2E-06 |
| Hand2 | Transcriptio | BHLH facto | Twist-like | Hand2 | dHand, Thi | 0.0002086 | 3.2809E-13 | 9.8E-06 | 4E-06 |
| Tmem131 | Co-nodes | Membrane | Transmem | Tmem131 | CC28, YR-2 | 0.000212 | 4.165E-11 | | 0.00062 |
| Ascl2 | Transcriptio | BHLH facto | Achaete-S | Ascl2 | ASH2, HAS | 0.0002459 | 3.2436E-16 | 0.01234 | 4.1E-05 |
| Usf2 | Transcriptio | BHLH facto | USF factor | Usf2 | FIP, bHLHb | 0.000285 | 6.2286E-18 | | |
| Kdm5a | Enzymes | Demethyla | Histone-H3 | Kdm5a | | 0.0003011 | 1.6575E-33 | | |

| | | | | | | | | | |
|---|---|---|---|---|---|---|---|---|---|
| Lhx2 | Transcriptio | Homeo do | Lhx-2-like | Lhx2 | LH-2, hLhx | 0.0003246 | | | 0.00326 |
| Taf7l | Co-nodes | General tr | TATA-box | Taf7l | CT40 | 0.000342 | 3.7812E-11 | 0.0094 | 2.7E-06 |
| Nelfe | Co-nodes | RNA bindi | Negative e | Nelfe | RD, D6S45 | 0.000342 | 4.0018E-22 | 0.00575 | 0.04692 |
| Tbp | Co-nodes | General tr | TATA-box | Tbp | TFIID | 0.0003697 | 4.891E-21 | 3.5E-07 | 7E-05 |
| Gps2 | Enzymes | Regulatory | G protein | Gps2 | | 0.0003778 | 0.00360504 | 0.00029 | 6.8E-09 |
| Wapl | Co-nodes | Other co-r | WAPL coh | Wapl | FOE | 0.0003912 | | | 0.00268 |
| Ets1 | Transcriptio | Tryptopha | Ets-like | Ets1 | FLJ10768, | 0.0004006 | 8.3953E-23 | | |
| Tbpl1 | Co-nodes | General tr | TATA-box | Tbpl1 | TLP, STUD | 0.0004085 | 8.4823E-12 | | |
| Brd2 | Co-nodes | Bromodon | Bromodon | Brd2 | KIAA9001, | 0.0004114 | 1.0839E-20 | | |
| Atrx | Enzymes | Helicases | ATRX chro | Atrx | XH2, XNP | 0.0004388 | 0.00165398 | 0.00139 | 2.3E-07 |
| Tfe3 | Transcriptio | BHLH fact | TFE3-like | Tfe3 | TFEA, bHL | 0.000452 | 7.0517E-05 | | 0.00584 |
