## Supplementary Data 5 for "*Let-7* restrains an oncogenic circuit in AT2 cells to prevent fibrogenic cell intermediates in pulmonary fibrosis"

### Supplementary Data 5: CUT&RUN H3K27ac Results

#### CUT&RUN H3K27ac Results

| Chromosome | Start | Stop | Direction | - LOG10(p-value) | - LOG10(q-value | GeneBody | GeneBody+/-10kb |
|---|---|---|---|---|---|---|---|
| chr1 | 3029001 | 3030000 | Up | 3.295960173 | 2.013652942 | | |

CUT&RUN-seq data has been deposited in the NCBI Gene Expression Omnibus database.

#### CUT&RUN H3K27me3 Results

| Chromosome | Start | Stop | Direction | - LOG10(p-value) | - LOG10(q-value | GeneBody | GeneBody+/-10kb |
|---|---|---|---|---|---|---|---|
| chr1 | 4455101 | 4457700 | Down | 2.288278621 | 1.158271748 | | |

CUT&RUN-seq data has been deposited in the NCBI Gene Expression Omnibus database.
