## Supplementary Data 6 for "*Let-7* restrains an oncogenic circuit in AT2 cells to prevent fibrogenic cell intermediates in pulmonary fibrosis"

### Supplementary Data 6: GREAT tool GO pathway enrichment.

Upregulated H3K27ac peaks in let-7afd-/- AT2 cells vs controls

| # Ontology | ID | Desc | BinomRank | BinomP | BinomBonfP | BinomFdrQ |
|---|---|---|---|---|---|---|
| GO Biological Process | GO:0022402 | cell cycle process | 1 | **3.25E-20** | 4.18E-16 | 4.18E-16 |
| GO Biological Process | GO:0051726 | regulation of cell cycle | 2 | **3.84E-18** | 4.94E-14 | 2.47E-14 |
| GO Biological Process | GO:0000278 | mitotic cell cycle | 3 | **1.20E-15** | 1.55E-11 | 5.15E-12 |
| GO Biological Process | GO:1901701 | cellular response to oxygen-containing compo | 4 | **1.95E-15** | 2.51E-11 | 6.28E-12 |
| GO Biological Process | GO:0012501 | programmed cell death | 5 | **4.20E-15** | 5.40E-11 | 1.08E-11 |
| GO Biological Process | GO:0006974 | cellular response to DNA damage stimulus | 6 | **4.54E-15** | 5.84E-11 | 9.74E-12 |
| GO Biological Process | GO:0042327 | positive regulation of phosphorylation | 7 | **6.68E-15** | 8.59E-11 | 1.23E-11 |
| GO Biological Process | GO:1903047 | mitotic cell cycle process | 8 | **7.83E-15** | 1.01E-10 | 1.26E-11 |
| GO Biological Process | GO:0001934 | positive regulation of protein phosphorylation | 9 | **8.21E-15** | 1.06E-10 | 1.17E-11 |
| GO Biological Process | GO:0016192 | vesicle-mediated transport | 10 | **1.02E-14** | 1.31E-10 | 1.31E-11 |
| GO Biological Process | GO:0008219 | cell death | 11 | **1.34E-14** | 1.73E-10 | 1.57E-11 |
| GO Biological Process | GO:0007346 | regulation of mitotic cell cycle | 12 | **2.07E-14** | 2.66E-10 | 2.21E-11 |
| GO Biological Process | GO:0051338 | regulation of transferase activity | 13 | **2.29E-14** | 2.94E-10 | 2.26E-11 |
| GO Biological Process | GO:0002684 | positive regulation of immune system process | 14 | **9.06E-14** | 1.17E-09 | 8.33E-11 |
| GO Biological Process | GO:0006915 | apoptotic process | 15 | **1.01E-13** | 1.30E-09 | 8.64E-11 |
| GO Biological Process | GO:1902533 | positive regulation of intracellular signal transd | 16 | **1.52E-13** | 1.96E-09 | 1.23E-10 |
| GO Biological Process | GO:0060548 | negative regulation of cell death | 17 | **2.23E-13** | 2.87E-09 | 1.69E-10 |
| GO Biological Process | GO:0051345 | positive regulation of hydrolase activity | 18 | **2.98E-13** | 3.83E-09 | 2.13E-10 |
| GO Biological Process | GO:0072359 | circulatory system development | 19 | **3.81E-13** | 4.90E-09 | 2.58E-10 |
| GO Biological Process | GO:0043207 | response to external biotic stimulus | 20 | **3.92E-13** | 5.05E-09 | 2.52E-10 |
| GO Biological Process | GO:0010564 | regulation of cell cycle process | 21 | **4.92E-13** | 6.33E-09 | 3.01E-10 |
| GO Biological Process | GO:0043549 | regulation of kinase activity | 22 | **8.30E-13** | 1.07E-08 | 4.85E-10 |
| GO Biological Process | GO:0071495 | cellular response to endogenous stimulus | 23 | **1.10E-12** | 1.41E-08 | 6.13E-10 |
| GO Biological Process | GO:0051707 | response to other organism | 24 | **1.23E-12** | 1.58E-08 | 6.58E-10 |
| GO Biological Process | GO:0009607 | response to biotic stimulus | 25 | **1.41E-12** | 1.81E-08 | 7.23E-10 |
| GO Biological Process | GO:0006461 | protein complex assembly | 26 | **1.48E-12** | 1.90E-08 | 7.32E-10 |
| GO Biological Process | GO:0070271 | protein complex biogenesis | 27 | **1.55E-12** | 1.99E-08 | 7.37E-10 |
| GO Biological Process | GO:1901135 | carbohydrate derivative metabolic process | 28 | **1.71E-12** | 2.20E-08 | 7.84E-10 |
| GO Biological Process | GO:0048646 | anatomical structure formation involved in mor | 29 | **6.13E-12** | 7.88E-08 | 2.72E-09 |
