## Supplementary Data 7 for "*Let-7* restrains an oncogenic circuit in AT2 cells to prevent fibrogenic cell intermediates in pulmonary fibrosis"

Supplementary Data 7: Known TF Motifs associated with increased H3K27ac peaks in *let-7afd-/-* AT2 cells

| Rank | Consensus Motif | P-value | q-value | Motif Name |
|---|---|---|---|---|
| 1 | 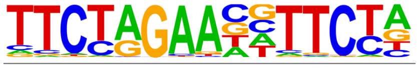 | 1E-15 | 4.00E-04 | HRE(HSF)/Striatum-HSF1-ChIP-Seq(GSE38000)/Homer |
| 2 | 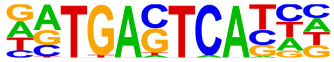 | 1E-15 | 1.60E-03 | Atf3(bZIP)/GBM-ATF3-ChIP-Seq(GSE33912)/Homer |
| 3 | 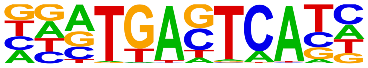 | 1E-15 | 1.60E-03 | Fra1(bZIP)/BT549-Fra1-ChIP-Seq(GSE46166)/Homer |
| 4 | 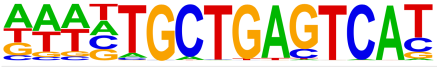 | 1E-15 | 1.60E-03 | Bach1(bZIP)/K562-Bach1-ChIP-Seq(GSE31477)/Homer |
