## Supplementary Data 8 for "*Let-7* restrains an oncogenic circuit in AT2 cells to prevent fibrogenic cell intermediates in pulmonary fibrosis"

**Supplementary Data 8: Induced genes with H3K27ac peaks**

| Gene Symbol | Category | Class | Family | ChIP-Seq consensome percentile | | | | | | | | |
|---|---|---|---|---|---|---|---|---|---|---|---|---|
| | | | | Arid3a | Bach1 | E2f1 | E2f3 | Ezh2 | Hif1a | Myc | | Count |
| *Pim1* | Enzymes | Kinases | Pim-1 kinases (PIM) | 87 | 61 | 95 | 99 | 65 | 54 | 72 | | 7 |
| *Aurkb* | Enzymes | Kinases | Aurora-related kinases (AURK) | 76 | | 63 | 96 | 59 | 72 | 75 | | 6 |
| *Hexb* | Enzymes | Glycosidases | Beta-N-acetylhexosaminidases (HEX) | 97 | 73 | | 99 | 99 | 99 | 99 | | 6 |
| *Mpzl1* | Co-nodes | CNS proteins | Myelin protein zero like | 97 | | 66 | 80 | 78 | 62 | 77 | | 6 |
| *Alg8* | Enzymes | Glycosyltransferases | Asparagine-linked glycosylation protein 6 ho | 58 | 60 | 78 | 98 | | | 97 | | 5 |
| *Arid3a* | Transcription factors | ARID domain | ARID3 family | 84 | 87 | | 74 | 89 | | 87 | | 5 |
| *Atad2* | Enzymes | Phosphatases | ATPase AAA domain (ATAD) | | 61 | 95 | 99 | 58 | | 91 | | 5 |
| *Bach1* | Transcription factors | BZIP factors | NF-E2-like factor | 75 | 93 | 52 | 62 | | | 77 | | 5 |
| *Cd68* | Receptors | Other receptors | Scavenger receptors | | 56 | 64 | 77 | 52 | | 99 | | 5 |
| *Cdca2* | Enzymes | Enzyme regulators | Protein phosphatase 1 regulatory subunits | | 81 | 86 | 94 | 65 | | 99 | | 5 |
| *Cdca5* | Co-nodes | Cell cycle, cell division and DNA repair | Cell division cycle associated | 92 | 90 | 81 | 95 | | | 79 | | 5 |
| *Cks2* | Enzymes | Enzyme regulators | CDC28 protein kinase regulatory subunit | 92 | 53 | | 70 | 55 | | 99 | | 5 |
| *Eif4a1* | Enzymes | Helicases (RNA) | Eukaryotic initiation factors 4A (EIF4A) | | 56 | 64 | 90 | 52 | | 99 | | 5 |
| *Gins1* | Co-nodes | Cell cycle, cell division and DNA repair | GINS complex subunit | | 54 | 82 | 94 | 62 | | 75 | | 5 |
| *Hat1* | Enzymes | Acetyltransferases | Histone acetyltransferases | 82 | | 58 | 60 | | 71 | 81 | | 5 |
| *Nuf2* | Co-nodes | Cell cycle, cell division and DNA repair | NUF2 component of NDC80 kinetochore com | 91 | 87 | 67 | 97 | | | 69 | | 5 |
| *Palb2* | Co-nodes | Cell cycle, cell division and DNA repair | Partner and localizer of BRCA2 | 95 | 63 | 58 | | | 92 | 70 | | 5 |
| *Rad51ap1* | Co-nodes | Cell cycle, cell division and DNA repair | Cell cycle checkpoint proteins (RAD) | 65 | 62 | 67 | 92 | | | 88 | | 5 |
| *Tyro3* | Receptors | Catalytic receptors | TAM (TYRO3-, AXL- and MER-TK) receptor fa | 83 | 55 | 62 | | 83 | | 59 | | 5 |
| *Abhd17c* | Enzymes | Esterases | Abhydrolases domain containing (ABHD) | | | 79 | 51 | 56 | | 92 | | 4 |
| *Adam10* | Enzymes | Peptidases | ADAM metallopeptidase domain (ADAM) | | 78 | | 63 | 74 | | 61 | | 4 |
| *Asap1* | Enzymes | Enzyme regulators | ArfGAP with SH3 domain, ankyrin repeat and PH domai | | 89 | | 52 | 83 | | 77 | | 4 |
| *Asxl1* | Co-nodes | Transcriptional coregulators | ASXL transcriptional regulator | | | 98 | 95 | | 56 | 88 | | 4 |
| *B4galt1* | Enzymes | Glycosyltransferases | Beta-1,4-galactosyl transferases (B4GALT) | 62 | | | 54 | 62 | | 55 | | 4 |
| *Bub1b* | Enzymes | Kinases | BUB mitotic checkpoint proteins | 70 | | 76 | 75 | | | 89 | | 4 |
| *Chek1* | Enzymes | Kinases | Checkpoint kinases (CHEK) | | 55 | 63 | 96 | | | 79 | | 4 |
| *Cks1b* | Enzymes | Enzyme regulators | CDC28 protein kinase regulatory subunit | | 54 | 76 | 92 | | | 95 | | 4 |
| *Cntrl* | Co-nodes | Cell cycle, cell division and DNA repair | Centriolin | | | 89 | 94 | | 52 | 87 | | 4 |
| *Diaph3* | Co-nodes | Other co-nodes | Diaphanous related formin | 70 | | 64 | 86 | | | 61 | | 4 |
| *Eno1* | Enzymes | Lyases | Enolases (ENO) | 87 | 85 | | | | 99 | 94 | | 4 |
| *Espl1* | Enzymes | Peptidases | Extra spindle pole bodies like, separases | | 56 | 64 | 89 | | | 93 | | 4 |
| *Ezh2* | Co-nodes | Polycomb group (PcG) proteins | Enhancer of zeste polycomb repressive complex subuni | | 50 | 98 | 99 | | | 94 | | 4 |
| *Fancd2* | Co-nodes | Cell cycle, cell division and DNA repair | FA complementation group | 94 | | 59 | 83 | | | 66 | | 4 |
| *Gabpb1* | Transcription factors | Tryptophan cluster | GA binding protein transcription factor subu | 88 | | 80 | | 50 | | 80 | | 4 |
| *Glg1* | Co-nodes | Endosomal, lysosomal, ER and Golgi pr | Golgi glycoprotein | | | | 91 | 65 | 78 | 61 | | 4 |
| *Haus6* | Co-nodes | Cell cycle, cell division and DNA repair | HAUS augmin like complex subunit | 81 | | 57 | 98 | | | 79 | | 4 |
| *Hif1a* | Transcription factors | BHLH factors | Ahr-like family | | 73 | | 71 | 53 | | 76 | | 4 |
| *Hk1* | Enzymes | Kinases | Hexokinase kinases (HK) | 53 | | 58 | | | 92 | 53 | | 4 |
| *Hmox1* | Enzymes | Oxidoreductases | Heme oxygenases (HMOX) | | 99 | | | 84 | 95 | 62 | | 4 |
| *Lpgat1* | Enzymes | Acyltransferases | Lysophosphatidylglycerol acyltransferases | | | 61 | 56 | 78 | | 52 | | 4 |
| *Mms22l* | Co-nodes | Cell cycle, cell division and DNA repair | MMS22 like, DNA repair protein | | | 55 | 98 | 65 | | 62 | | 4 |
| *Ncapd2* | Co-nodes | Cell cycle, cell division and DNA repair | Non-SMC condensin complex subunits | | 65 | 64 | 95 | | | 90 | | 4 |
| *Ncapg2* | Co-nodes | Cell cycle, cell division and DNA repair | Non-SMC condensin complex subunits | | | 94 | 99 | 50 | | 68 | | 4 |
| *Nipa2* | Co-nodes | Transporters and transport proteins | NIPA magnesium transporter | 68 | | | 73 | 59 | | 63 | | 4 |
| *Noxo1* | Enzymes | Enzyme regulators | NADPH oxidase organizer | | | 88 | 84 | 72 | | 81 | | 4 |
| *Pik3cd* | Enzymes | Kinases | Phosphatidylinositol-4-phosphate 3 kinases ( | 93 | | | | 75 | 72 | 66 | | 4 |
| *Pold1* | Enzymes | Nucleotidyltransferases | DNA-directed DNA polymerases (POL) | | | 65 | 86 | | 69 | 76 | | 4 |
| *Pole2* | Enzymes | Nucleotidyltransferases | DNA-directed DNA polymerases (POL) | 66 | | 70 | 95 | | | 58 | | 4 |
| *Rrm1* | Enzymes | Reductases | Ribonucleoside-diphosphate reductases (RR | 75 | | 88 | 97 | | | 84 | | 4 |
| *Serpine1* | Receptors | Ligands | Serpins | 66 | 61 | | | | 70 | 65 | | 4 |
| *Slc20a1* | Co-nodes | Transporters and transport proteins | Solute carrier superfamily member | | 95 | | 66 | 53 | | 98 | | 4 |
| *Slc7a6* | Co-nodes | Transporters and transport proteins | Solute carrier superfamily member | | 52 | 70 | | 75 | | 90 | | 4 |
| *Smc6* | Co-nodes | Cell cycle, cell division and DNA repair | Structural maintenance of chromosomes | | | 75 | 95 | 54 | | 84 | | 4 |
| *Ube2m* | Enzymes | E2 ubiquitin conjugating enzymes | Ubiquitin conjugating enzymes E2 (UBE2) | 55 | | 56 | | 87 | | 69 | | 4 |
| *Wdhd1* | Co-nodes | WD repeat proteins | WD repeat and HMG-box DNA binding protein | | | 78 | 99 | 52 | | 84 | | 4 |
| *Wdr76* | Co-nodes | WD repeat proteins | WD repeat domain | 80 | | 99 | 99 | | | 55 | | 4 |
| *Acvr1b* | Receptors | Catalytic receptors | Transforming growth factor-? receptor fami | 78 | 68 | | | 56 | | | | 3 |
| *Arhgap11a* | Enzymes | Enzyme regulators | Rho GTPase activating proteins | 60 | | 67 | | | | 65 | | 3 |
| *Ccna2* | Enzymes | Enzyme regulators | Cyclins (CCN) | | | 88 | 87 | | | 75 | | 3 |
| *Ccnf* | Enzymes | Enzyme regulators | Cyclins (CCN) | | | 90 | 90 | | | 67 | | 3 |
| *Cdk1* | Enzymes | Kinases | Cyclin-dependent kinases (CDK) | | | 97 | 98 | | | 76 | | 3 |
| *Cenpf* | Co-nodes | Cell cycle, cell division and DNA repair | Centromere protein | | | | 95 | | 81 | 67 | | 3 |
| *Coq7* | Enzymes | Other enzymes | Coenzyme Q | 56 | | 57 | | | | 63 | | 3 |
| *Crlf2* | Receptors | Catalytic receptors | Interleukin 2 (IL2) receptor family | 65 | 62 | | | | | 51 | | 3 |
| *Dck* | Enzymes | Kinases | Deoxycytidine kinases | | | | 96 | 56 | | 61 | | 3 |
| *Efna3* | Receptors | Ligands | Ephrin | 65 | | | | 95 | | 56 | | 3 |
| *Fam107b* | Co-nodes | Family with sequence similarity | Family with sequence similarity member | | 82 | | | 71 | | 60 | | 3 |
| *Frrs1* | Enzymes | Oxidoreductases | Ferric chelate reductases | 69 | | | | 66 | | 67 | | 3 |
| *Gcnt2* | Enzymes | Glycosyltransferases | Beta-1,6-N-acetylglucosaminyltransferases | 90 | | | 71 | | | 56 | | 3 |
| *Ifngr2* | Receptors | Catalytic receptors | Interferon gamma receptor | 63 | | | | 73 | | 59 | | 3 |
| *Inf2* | Co-nodes | Other co-nodes | Inverted formin | | | | 84 | 76 | | 80 | | 3 |
| *Iqgap3* | Enzymes | Enzyme regulators | IQ motif containing GTPase activating proteins | | | 69 | 97 | | | 65 | | 3 |
| *Itga6* | Receptors | Catalytic receptors | Integrins | | | 90 | 78 | 59 | | | | 3 |
| *Lamc1* | Receptors | Ligands | Laminin subunit | | 96 | | | 65 | | 63 | | 3 |
| *Lcn2* | Co-nodes | Transporters and transport proteins | Lipocalin | | | | | 57 | 69 | 86 | | 3 |
| *Lin9* | Co-nodes | Transcriptional coregulators | Lin-9 DREAM MuvB core complex component | | | 83 | 83 | | | 53 | | 3 |
| *Man2a2* | Enzymes | Glycosidases | Mannosidases (MAN) | 78 | | | 64 | | | 54 | | 3 |
| *Me1* | Enzymes | Dehydrogenases | Malate, oxaloacetate-decarboxylating (ME) | | 95 | | | 79 | 59 | | | 3 |
| *Metap1* | Enzymes | Peptidases | Methionyl aminopeptidases (METAP) | | 72 | | 52 | | | 91 | | 3 |
| *Mreg* | Co-nodes | Endosomal, lysosomal, ER and Golgi pr | Melanoregulin | | | 95 | | 72 | | 70 | | 3 |
| *Ndufab1* | Enzymes | Oxidoreductases | NADH:ubiquinone oxidoreductase subunits | | | 64 | 83 | | | 75 | | 3 |
| *Nek2* | Enzymes | Kinases | NimA related kinases (NEK) | | 76 | | | | 74 | 72 | | 3 |
| *Nras* | Enzymes | GTPases | Ras Type GTPases | 54 | 77 | | | | | 91 | | 3 |
| *Nrm* | Co-nodes | Other co-nodes | Nurim | | 57 | | 89 | | | 79 | | 3 |

| | | | | | | | | | | | | |
|---|---|---|---|---|---|---|---|---|---|---|---|---|
| *Nudt4* | Enzymes | Phosphatases | Nudix (NUDT) | | | 91 | 86 | | | 69 | | 3 |
| *Nusap1* | Co-nodes | Ribosomes and ribosome biogenesis | Nucleolar and spindle associated protein | | | 81 | 77 | | | 77 | | 3 |
| *Oip5* | Co-nodes | Interacting proteins | Opa interacting protein | | | 81 | 77 | | | 77 | | 3 |
| *Orc1* | Co-nodes | Cell cycle, cell division and DNA repair | Origin recognition complex subunit | | | | 56 | | 50 | 66 | | 3 |
| *Pidd1* | Co-nodes | Apoptosis and apoptosis regulators | P53-induced death domain protein | | | 90 | 98 | | | 94 | | 3 |
| *Plagl2* | Transcription factors | C2H2 Zn finger factors | PLAG Zinc Finger | | | 87 | 89 | | | 68 | | 3 |
| *Plch1* | Enzymes | Lipases | Phosphoinositide phospholipases C (PLC) | | | | 82 | 57 | 78 | | | 3 |
| *Pmepa1* | Co-nodes | Androgen regulated | Prostate transmembrane protein, androgen | 51 | | 67 | | 57 | | | | 3 |
| *Poc1a* | Co-nodes | Cell cycle, cell division and DNA repair | POC1 centriolar protein | 71 | 86 | 50 | | | | | | 3 |
| *Pola1* | Enzymes | Nucleotidyltransferases | DNA-directed DNA polymerases (POL) | | | 61 | 90 | | | 59 | | 3 |
| *Prr11* | Co-nodes | Proline rich proteins | Proline rich | 58 | | | 80 | | | 56 | | 3 |
| *Prrg4* | Co-nodes | Proline rich proteins | Proline rich and Gla domain | 56 | | | | 71 | | 52 | | 3 |
| *Rfc5* | Co-nodes | Cell cycle, cell division and DNA repair | Replication factor C subunit | | | | 90 | | 82 | 92 | | 3 |
| *Rnh1* | Enzymes | Enzyme regulators | Ribonuclease/angiogenin inhibitor | | 75 | | | 56 | | 91 | | 3 |
| *Smc4* | Co-nodes | Cell cycle, cell division and DNA repair | Structural maintenance of chromosomes | | | 85 | 88 | | | 65 | | 3 |
| *Spata13* | Enzymes | Enzyme regulators | Rho guanine nucleotide exchange factors | | | | 87 | 57 | | 68 | | 3 |
| *Synpo* | Co-nodes | Cytoskeleton components and regulator | Synaptopodin | | 62 | | | 72 | | 53 | | 3 |
| *Timeless* | Co-nodes | Circadian factors | Timeless circadian regulator | | | 68 | 96 | | | 52 | | 3 |
| *Tk1* | Enzymes | Kinases | Thymidine kinases (TK) | | | | 90 | | 93 | 75 | | 3 |
| *Tubgcp2* | Co-nodes | Cytoskeleton components and regulator | Tubulin gamma complex associated protein | | | | 96 | 63 | | 74 | | 3 |
| *Wwc1* | Enzymes | Enzyme regulators | Protein phosphatase 1 regulatory subunits | | | 64 | | 76 | | 56 | | 3 |
| *Ybx2* | Transcription factors | Cold shock domain | FRG Y2-like | | 54 | | 72 | 88 | | | | 3 |
| *Zmat3* | Co-nodes | Zinc finger proteins | Zinc finger matrin-type | | | | 82 | 73 | | 82 | | 3 |
| *Cldn4* | Co-nodes | Junction proteins | Claudin | 94 | 84 | | 87 | | | | | 3 |
| *E2f2* | Transcription factors | E2F/FOX | E2F | | | 61 | 95 | | | 79 | | 3 |
| *1700066B1* | Co-nodes | Uncharacterized transcripts | Riken genes | | | | 96 | | | 51 | | 2 |
| *Agrn* | Receptors | Ligands | Agrin | | | 81 | | 95 | | | | 2 |
| *Arhgdib* | Enzymes | Enzyme regulators | Rho GDP-dissociation inhibitors | | | | | 67 | 64 | | | 2 |
| *Cacna1c* | Ion channels | Voltage gated ion channels | Calcium voltage-gated channel subunit | | | | | 99 | | 66 | | 2 |
| *Cd300e* | Co-nodes | Cell surface proteins | Cluster of differentiation | 99 | | | 85 | | | | | 2 |
| *Cdc20* | Co-nodes | Cell cycle, cell division and DNA repair | Cell division cycle | | | | 59 | | | 77 | | 2 |
| *Clcn5* | Ion channels | Other ion channels | Chloride voltage-gated channel | | | | 68 | 74 | | | | 2 |
| *Csf1r* | Receptors | Catalytic receptors | M-CSF/CSF1 receptor | | 58 | | | 81 | | | | 2 |
| *Dab2* | Co-nodes | Adaptor, docking and scaffolding nodes | DAB adaptor protein | | | | | 66 | | 66 | | 2 |
| *Ddx39* | Enzymes | Helicases (RNA) | DEAD box (DDX) | | | | | 61 | | 89 | | 2 |
| *Dhx58* | Receptors | Catalytic receptors | RIG-I-like receptors | | 61 | | | | | 98 | | 2 |
| *Dlgap5* | Co-nodes | Cell cycle, cell division and DNA repair | DLG associated protein | | | 70 | 63 | | | | | 2 |
| *Fam221a* | Co-nodes | Family with sequence similarity | Family with sequence similarity member | 54 | | | 62 | | | | | 2 |
| *Fam83d* | Co-nodes | Family with sequence similarity | Family with sequence similarity member | | | | 85 | | | 54 | | 2 |
| *Fetub* | Co-nodes | Other co-nodes | Fetuin B | 77 | 79 | | | | | | | 2 |
| *Fmnl1* | Co-nodes | Cytoskeleton components and regulator | Formin like | | | | | 88 | | 67 | | 2 |
| *Focad* | Co-nodes | Adhesion molecules | Focadhesin | | | | | 73 | | 97 | | 2 |
| *Galnt2* | Enzymes | Glycosyltransferases | Polypeptide GalNAc transferases (GALNT) | | | 66 | | 53 | | | | 2 |
| *Gdf11* | Receptors | Ligands | Growth differentiation factor | | 55 | | | 73 | | | | 2 |
| *Ggta1* | Enzymes | Glycosyltransferases | Glycoprotein galactosyltransferase | | | | 81 | | 61 | | | 2 |
| *Gpc3* | Receptors | Ligands | Glypican | | | | 55 | 69 | | | | 2 |
| *Hmmr* | Co-nodes | Cell surface proteins | Hyaluronan mediated motility receptor | | | | 94 | | | 95 | | 2 |
| *Igf2bp2* | Co-nodes | RNA binding and RB motif proteins | Insulin like growth factor mRNA binding protein | | | 61 | | 92 | | | | 2 |
| *Kif14* | Co-nodes | Cell cycle, cell division and DNA repair | Kinesin | | | | 76 | | | 61 | | 2 |
| *Lacc1* | Co-nodes | Other co-nodes | Laccase domain containing | | | | 60 | 81 | | | | 2 |
| *Lama5* | Receptors | Ligands | Laminin subunit | | 89 | | | 78 | | | | 2 |
| *Lrcol1* | Co-nodes | Leucine rich proteins | Leucine rich colipase like | 73 | | | | 69 | | | | 2 |
| *Ly75* | Receptors | Other receptors | Scavenger receptors | | | | | 66 | | 56 | | 2 |
| *Lztr1* | Co-nodes | BTB domain containing | Leucine zipper like transcription regulator | 56 | | | | | | 83 | | 2 |
| *Mad2l1* | Co-nodes | Cell cycle, cell division and DNA repair | Mitotic arrest deficient like | | | | 92 | | | 60 | | 2 |
| *Malt1* | Co-nodes | Other enzymes | MALT1 paracaspase | | | | | 85 | | 61 | | 2 |
| *Map6* | Co-nodes | Cytoskeleton components and regulator | Microtubule associated protein | | | | | 93 | 82 | | | 2 |
| *Melk* | Enzymes | Kinases | Maternal embryonic leucine zipper kinases | | | 63 | 79 | | | | | 2 |
| *Mrpl11* | Co-nodes | Ribosomes and ribosome biogenesis | Mitochondrial ribosomal protein | 90 | | | | | | 89 | | 2 |
| *Mrps18b* | Co-nodes | Ribosomes and ribosome biogenesis | Mitochondrial ribosomal protein | | | | 84 | | | 95 | | 2 |
| *Ncf4* | Co-nodes | Other co-nodes | Neutrophil cytosolic factor | 67 | | | 72 | | | | | 2 |
| *Ncl* | Co-nodes | RNA binding and RB motif proteins | Nucleolin | | | | 83 | | | 99 | | 2 |
| *Ola1* | Enzymes | ATPases | Obg like ATPases | | | | | 59 | | 91 | | 2 |
| *Pigs* | Co-nodes | Other co-nodes | Phosphatidylinositol glycan subunits (PIG) | | | | | | 85 | 59 | | 2 |
| *Plaur* | Receptors | Other receptors | Plasminogen activator, urokinase receptor | | 97 | | | | | 62 | | 2 |
| *Psmc1* | Co-nodes | Proteasome | Proteasome 26S subunit, ATPase | | | 73 | | | | 63 | | 2 |
| *Rab15* | Enzymes | GTPases | RAB, member RAS oncogene | | | | 64 | 85 | | | | 2 |
| *Rab27a* | Enzymes | GTPases | RAB, member RAS oncogene | 68 | | | | | | 88 | | 2 |
| *Rab8b* | Enzymes | GTPases | RAB, member RAS oncogene | | | | | 92 | | 51 | | 2 |
| *Rac2* | Enzymes | GTPases | Rho GTPases | | 69 | | | | | 65 | | 2 |
| *Rad21* | Co-nodes | Cohesin complex | Cell cycle checkpoint proteins (RAD) | | | 88 | 79 | | | | | 2 |
| *Rasa3* | Enzymes | Enzyme regulators | RAS p21 protein activator | | | | | 56 | | 67 | | 2 |
| *Rnf26* | Co-nodes | Ring finger proteins | Ring finger protein | | | 77 | 87 | | | | | 2 |
| *Sbno2* | Enzymes | Helicases (RNA) | Strawberry notch homolog | | 78 | | | | | 71 | | 2 |
| *Shcbp1* | Co-nodes | Other co-nodes | SHC binding and spindle associated | 74 | | 83 | | | | | | 2 |
| *Slc16a3* | Co-nodes | Transporters and transport proteins | Solute carrier superfamily member | | | | | 60 | 99 | | | 2 |
| *Spns2* | Co-nodes | Lipid metabolism | Sphingolipid transporter | | | | | 89 | | 75 | | 2 |
| *St3gal4* | Enzymes | Glycosyltransferases | Beta-galactoside alpha-2,3-sialyltransferases (ST3GAL) | | | | | 64 | | 89 | | 2 |
| *Susd1* | Co-nodes | Other co-nodes | Sushi domain containing | | | | | 91 | | 51 | | 2 |
| *Tnfrsf1b* | Receptors | Catalytic receptors | Tumor necrosis factor receptors | | | | | 58 | | 70 | | 2 |
| *Xpnpep1* | Enzymes | Peptidases | Xaa-Pro aminopeptidases (XPNPEP) | | | | | | 55 | 85 | | 2 |
| *Xrcc6* | Co-nodes | Cell cycle, cell division and DNA repair | X-ray repair cross complementing | | | | 84 | | | 95 | | 2 |
| *Xrn1* | Enzymes | Ribonucleases | 5'-3' exoribonucleases | 90 | | | | | | 65 | | 2 |
| *Myc* | Transcription factors | BHLH factors | Myc/Max factor | | 57 | | 95 | | | | | 2 |
| *Abcg1* | Co-nodes | Transporters and transport proteins | ATP binding cassette | | | | | | | 55 | | 1 |
| *Akt1* | Enzymes | Kinases | Protein kinase B/Akt kinases (AKT) | | | | | 92 | | | | 1 |
| *Alox5ap* | Enzymes | Enzyme regulators | Arachidonate 5-lipoxygenase activating protein | | 51 | | | | | | | 1 |
| *Capzb* | Co-nodes | Cytoskeleton components and regulator | Capping actin protein of muscle Z-line subunit | | | | | | | 63 | | 1 |
| *Cd276* | Co-nodes | Cell surface proteins | Cluster of differentiation | | 92 | | | | | | | 1 |
| *Cdkn3* | Enzymes | Enzyme regulators | Cyclin-dependent kinase inhibitors (CDKN) | | | 50 | | | | | | 1 |
| *Cenpe* | Enzymes | Enzyme regulators | Protein phosphatase 1 regulatory subunits | | | | 65 | | | | | 1 |
| *Cnih1* | Co-nodes | Receptor associated factors | Cornichon AMPA receptor auxiliary protein | | | | | | | 64 | | 1 |
| *Coq3* | Enzymes | Methyltransferases | 3-demethylubiquinol 3-O-methyltransferases (COQ) | | | | | | | 68 | | 1 |

| | | | | | | | | | | | | |
|---|---|---|---|---|---|---|---|---|---|---|---|---|
| *Ctse* | Enzymes | Peptidases | Cathepsins (CTS) | | | | | 74 | | | | 1 |
| *Dock10* | Co-nodes | Cell cycle, cell division and DNA repair | Dedicator of cytokinesis | | | | | 94 | | | | 1 |
| *Dynlt1b* | Co-nodes | Cell cycle, cell division and DNA repair | Dynein light chain Tctex-type | 95 | | | | | | | | 1 |
| *Emilin2* | Co-nodes | Other co-nodes | Elastin microfibril interfacer | | | | | 94 | | | | 1 |
| *Flna* | Co-nodes | Cytoskeleton components and regulator | Filamin | | | | | | | 62 | | 1 |
| *Gla* | Enzymes | Glycosidases | Galactosidase alpha | | 96 | | | | | | | 1 |
| *Gpnmb* | Co-nodes | Glycoproteins | Glycoprotein nmb | | | | 76 | | | | | 1 |
| *Il1b* | Receptors | Ligands | Interleukin | | | | 54 | | | | | 1 |
| *Il1rl2* | Receptors | Catalytic receptors | Interleukin 1 receptor like | | | | | 78 | | | | 1 |
| *Ints2* | Co-nodes | Transcriptional coregulators | Integrator complex subunit | | | | | | | 77 | | 1 |
| *Itga2* | Receptors | Catalytic receptors | Integrins | | | | 63 | | | | | 1 |
| *Kifc5b* | Co-nodes | Cell cycle, cell division and DNA repair | Kinesin | | | | | | | 96 | | 1 |
| *Klhdc8a* | Co-nodes | Kelch domain proteins | Kelch domain containing | | | | | 94 | | | | 1 |
| *Klhl32* | Co-nodes | Kelch domain proteins | Kelch like | | | | | 81 | | | | 1 |
| *Lamb1* | Receptors | Ligands | Laminin subunit | | | | | 76 | | | | 1 |
| *Lamc2* | Receptors | Ligands | Laminin subunit | | | | | 78 | | | | 1 |
| *Lgals9* | Co-nodes | Lectins | Galectin | 86 | | | | | | | | 1 |
| *Lpcat3* | Enzymes | Acyltransferases | Lysophosphatidylcholine (LPCAT) | | | | | | | 54 | | 1 |
| *Lrg1* | Co-nodes | Glycoproteins | Leucine rich alpha-2-glycoprotein | | | | | 83 | | | | 1 |
| *Mab21l3* | Co-nodes | Other co-nodes | mab-21 like | | 85 | | | | | | | 1 |
| *Mcm10* | Co-nodes | Cell cycle, cell division and DNA repair | Minichromosome maintenance (MCM) | | | | 89 | | | | | 1 |
| *Mpzl2* | Co-nodes | CNS proteins | Myelin protein zero like | | | | | 65 | | | | 1 |
| *Ms4a6d* | Co-nodes | Membrane proteins | Membrane spanning-domains | | 59 | | | | | | | 1 |
| *Nlrc4* | Receptors | Catalytic receptors | NOD-like receptors | | | | 56 | | | | | 1 |
| *Pdzd2* | Co-nodes | PDZ domain proteins | PDZ domain containing | | | | | 79 | | | | 1 |
| *Pglyrp1* | Co-nodes | Antimicrobial factors | Peptidoglycan recognition protein | | | | | | | 76 | | 1 |
| *Pigt* | Co-nodes | Other co-nodes | Phosphatidylinositol glycan subunits (PIG) | | | | | | | 82 | | 1 |
| *Pla2g7* | Enzymes | Lipases | Phospholipases (PLA2) | | | | | 57 | | | | 1 |
| *Plek* | Co-nodes | Pleckstrin domain | Pleckstrin | | | | 81 | | | | | 1 |
| *Ppa1* | Enzymes | Phosphatases | Inorganic pyro- (PPA) | | | | | | | 89 | | 1 |
| *Prkch* | Enzymes | Kinases | Protein kinase C (PKC) | | | | | 91 | | | | 1 |
| *Pros1* | Receptors | Ligands | Protein S | 57 | | | | | | | | 1 |
| *Rasl12* | Enzymes | GTPases | Ras Type GTPases | | | | | 94 | | | | 1 |
| *Reln* | Receptors | Ligands | Reelin | | | | | 96 | | | | 1 |
| *Saa3* | Co-nodes | Amyloid proteins | Serum amyloid | | | | | | | 75 | | 1 |
| *Sdc3* | Co-nodes | Cell surface proteins | Syndecan | | | | | 91 | | | | 1 |
| *Spats2l* | Co-nodes | Testis, sperm and spermatogenesis | Spermatogenesis associated serine rich like | | | | | 91 | | | | 1 |
| *Spdl1* | Co-nodes | Coiled coil domain | Spindle apparatus coiled-coil protein | | | | | | | 53 | | 1 |
| *Sphk1* | Enzymes | Kinases | Sphingosine kinases (SPHK) | | | | | 95 | | | | 1 |
| *Spi1* | Transcription factors | Tryptophan cluster | Spi-like | | | | 82 | | | | | 1 |
| *Syt2* | Co-nodes | CNS proteins | Synaptotagmin | | | | | 99 | | | | 1 |
| *Tc2n* | Co-nodes | Other co-nodes | Tandem C2 domains, nuclear | | | | | 87 | | | | 1 |
| *Tceanc* | Co-nodes | General transcription factors | Transcription elongation factor A N-terminal and central domain containing | | | | | | | 53 | | 1 |
| *Thnsl2* | Co-nodes | Other co-nodes | Threonine synthase like | | | 63 | | | | | | 1 |
| *Tnfaip2* | Co-nodes | Immune system components | TNF alpha induced proteins | | | | | 99 | | | | 1 |
| *Troap* | Co-nodes | Interacting proteins | Trophinin associated protein | | | | | | | 55 | | 1 |
| *Ubash3b* | Enzymes | Phosphatases | Ubiquitin associated and SH3 domain containing | | | | | 88 | | | | 1 |
| *Vnn1* | Enzymes | Other enzymes | Vanin (VNN) | | | | | 70 | | | | 1 |
| *Wdfy4* | Co-nodes | Other co-nodes | WD repeat and FYVE domain containing | | | | | | | 58 | | 1 |
| *Arl11* | Enzymes | GTPases | ADP-ribosylation factor like GTPases | | | | | | | | | 0 |
| *Bcl2l15* | Co-nodes | Other co-nodes | BCL2 like | | | | | | | | | 0 |
| *C1ra* | Enzymes | Peptidases | Complement components and factors (C/CF) | | | | | | | | | 0 |
| *Ccdc36* | Co-nodes | Other co-nodes | Interactor of HORMAD | | | | | | | | | 0 |
| *Cd48* | Co-nodes | Cell surface proteins | Cluster of differentiation | | | | | | | | | 0 |
| *Cd53* | Co-nodes | Cell surface proteins | Cluster of differentiation | | | | | | | | | 0 |
| *Cep41* | Co-nodes | Cell cycle, cell division and DNA repair | Centrosomal protein | | | | | | | | | 0 |
| *Cit* | Enzymes | Kinases | Citron rho-interacting serine/threonine kinases | | | | | | | | | 0 |
| *Clu* | Co-nodes | Other co-nodes | Clusterin | | | | | | | | | 0 |
| *Col5a2* | Receptors | Ligands | Collagen chain | | | | | | | | | 0 |
| *Eln* | Co-nodes | Other co-nodes | Elastin | | | | | | | | | 0 |
| *Fam96a* | NA | NA | NA | | | | | | | | | 0 |
| *Fes* | Enzymes | Kinases | FES proto-oncogene, tyrosine kinases | | | | | | | | | 0 |
| *Fgr* | Enzymes | Kinases | Src kinases | | | | | | | | | 0 |
| *Filip1* | Co-nodes | Extracellular matrix | Filamin A interacting protein | | | | | | | | | 0 |
| *Fxyd4* | Co-nodes | Transporters and transport proteins | FXYD domain containing ion transport regulator | | | | | | | | | 0 |
| *Fyb* | Co-nodes | Other co-nodes | FYN binding protein | | | | | | | | | 0 |
| *Gpr132* | Receptors | G protein coupled receptors | G protein-coupled receptor | | | | | | | | | 0 |
| *Hcls1* | Co-nodes | Substrate proteins | Hematopoietic cell-specific Lyn substrate | | | | | | | | | 0 |
| *Hist1h1d* | Co-nodes | Histones and histone associated | H1 linker histone | | | | | | | | | 0 |
| *Hist1h2bg* | Co-nodes | Histones and histone associated | H2B clustered histone | | | | | | | | | 0 |
| *Igsf6* | Co-nodes | Immune system components | Immunoglobulin superfamily member | | | | | | | | | 0 |
| *Il18bp* | Co-nodes | Other co-nodes | Interleukin binding protein | | | | | | | | | 0 |
| *Il18rap* | Receptors | Catalytic receptors | Interleukin 18 receptor accessory protein | | | | | | | | | 0 |
| *Inpp5d* | Enzymes | Phosphatases | Inositol poly- (INPP) | | | | | | | | | 0 |
| *Itm2a* | Co-nodes | Membrane proteins | Integral membrane protein | | | | | | | | | 0 |
| *Kif4* | Co-nodes | Cell cycle, cell division and DNA repair | Kinesin | | | | | | | | | 0 |
| *Mat1a* | Enzymes | Transferases | Methionine adenosyltransferases (MAT) | | | | | | | | | 0 |
| *Mcoln3* | Ion channels | Voltage gated ion channels | Mucolipin | | | | | | | | | 0 |
| *Mid1* | Enzymes | E3 ubiquitin ligases | Midline (MID) | | | | | | | | | 0 |
| *Ms4a6c* | Co-nodes | Membrane proteins | Membrane spanning-domains | | | | | | | | | 0 |
| *Muc4* | Co-nodes | Glycoproteins | Mucin, cell surface associated | | | | | | | | | 0 |
| *Myo1f* | Co-nodes | Cytoskeleton components and regulator | Myosin | | | | | | | | | 0 |
| *Olr1* | Receptors | Other receptors | Scavenger receptors | | | | | | | | | 0 |
| *Orm1* | Receptors | Ligands | Orosomucoid | | | | | | | | | 0 |
| *Pde1c* | Enzymes | Esterases | Phosphodiesterases (PDE) | | | | | | | | | 0 |
| *Pfdn1* | Co-nodes | Chaperones | Prefoldin subunit | | | | | | | | | 0 |
| *Pkhd1* | Co-nodes | Other co-nodes | PKHD1 ciliary IPT domain containing fibrocystin/polyductin | | | | | | | | | 0 |
| *Plscr1* | Enzymes | Scramblases | Phospholipid scramblases | | | | | | | | | 0 |
| *Prg4* | Co-nodes | Extracellular matrix | Proteoglycan, pro eosinophil major basic protein | | | | | | | | | 0 |
| *Ptafr* | Receptors | G protein coupled receptors | Platelet-activating factor receptor | | | | | | | | | 0 |
| *Samsn1* | Co-nodes | Nuclear proteins | SAM domain, SH3 domain and nuclear localization signals | | | | | | | | | 0 |
| *Sla* | Co-nodes | SH2 domain containing | Src like adaptor | | | | | | | | | 0 |
| *Spc24* | Co-nodes | Cell cycle, cell division and DNA repair | SPC component of NDC80 kinetochore complex | | | | | | | | | 0 |

| | | | | | | | | | | | | |
|---|---|---|---|---|---|---|---|---|---|---|---|---|
| *Tmem173* | Co-nodes | Membrane proteins | Stimulator of interferon response cGAMP interactor | | | | | | | | | 0 |
| *Tmem37* | Co-nodes | Membrane proteins | Transmembrane protein | | | | | | | | | 0 |
| *Tnc* | Receptors | Ligands | Tenascin | | | | | | | | | 0 |
| *Trpv2* | Ion channels | Voltage gated ion channels | Transient receptor potential cation channel | | | | | | | | | 0 |
| *Vav1* | Enzymes | Enzyme regulators | Rho guanine nucleotide exchange factors | | | | | | | | | 0 |
