## Supplementary Data 9 for "*Let-7* restrains an oncogenic circuit in AT2 cells to prevent fibrogenic cell intermediates in pulmonary fibrosis"

*Let-7* targetome DEGs (IPF AB cells vs control AT2s)

| Gene | p_val | avg_log2FC | pct.1 | pct.2 | p_val_adj | Direction |
|---|---|---|---|---|---|---|
| Tmsb10 | 5.69E-94 | 2.146644556 | 0.942 | 0.833 | 2.62E-89 | Upregulated |
| Col1a1 | 0 | 2.121800638 | 0.752 | 0.056 | 0 | Upregulated |
| Tpm1 | 1.39E-191 | 1.813866337 | 0.75 | 0.174 | 6.39E-187 | Upregulated |
| Serpine1 | 1.30E-198 | 1.648077386 | 0.395 | 0.012 | 5.95E-194 | Upregulated |
| Slco2a1 | 0 | 1.555661532 | 0.667 | 0.02 | 0 | Upregulated |
| Apbb2 | 2.83E-129 | 1.553292126 | 0.868 | 0.365 | 1.30E-124 | Upregulated |
| Asap1 | 6.65E-166 | 1.471623475 | 0.815 | 0.241 | 3.06E-161 | Upregulated |
| Tmsb4x | 1.34E-58 | 1.446005362 | 0.973 | 0.924 | 6.16E-54 | Upregulated |
| Col6a2 | 0 | 1.43588413 | 0.583 | 0.006 | 0 | Upregulated |
| Lpp | 3.20E-116 | 1.290009635 | 0.982 | 0.839 | 1.47E-111 | Upregulated |
| Crip2 | 0 | 1.266059163 | 0.679 | 0.032 | 0 | Upregulated |
| Actb | 2.60E-49 | 1.257628797 | 0.926 | 0.833 | 1.20E-44 | Upregulated |
| Trim2 | 1.03E-127 | 1.208766655 | 0.808 | 0.287 | 4.75E-123 | Upregulated |
| S100a11 | 3.24E-46 | 1.183343883 | 0.842 | 0.751 | 1.49E-41 | Upregulated |
| Ntn1 | 9.53E-198 | 1.175115649 | 0.554 | 0.052 | 4.38E-193 | Upregulated |
| Igf2bp2 | 8.13E-96 | 1.11416703 | 0.828 | 0.383 | 3.73E-91 | Upregulated |
| Cfl1 | 4.33E-57 | 1.096735604 | 0.804 | 0.652 | 1.99E-52 | Upregulated |
| Zmat3 | 8.11E-228 | 1.059856913 | 0.636 | 0.067 | 3.73E-223 | Upregulated |
| Spint1 | 1.48E-90 | 1.055832739 | 0.714 | 0.328 | 6.79E-86 | Upregulated |
| Timp2 | 4.50E-172 | 0.955096018 | 0.645 | 0.108 | 2.07E-167 | Upregulated |
| Tpm4 | 3.31E-67 | 0.953650323 | 0.763 | 0.455 | 1.52E-62 | Upregulated |
| Golm1 | 1.22E-141 | 0.953619482 | 0.69 | 0.16 | 5.60E-137 | Upregulated |
| Serpinb9 | 1.99E-49 | 0.941856668 | 0.31 | 0.073 | 9.16E-45 | Upregulated |
| Tagln2 | 3.21E-59 | 0.934582527 | 0.75 | 0.498 | 1.48E-54 | Upregulated |
| Agrn | 1.44E-142 | 0.920941386 | 0.627 | 0.134 | 6.60E-138 | Upregulated |
| Lims1 | 8.57E-81 | 0.906249009 | 0.837 | 0.461 | 3.94E-76 | Upregulated |
| Marcks | 1.59E-135 | 0.901261918 | 0.609 | 0.13 | 7.29E-131 | Upregulated |
| Pfn1 | 4.28E-36 | 0.873690158 | 0.806 | 0.664 | 1.97E-31 | Upregulated |
| Pim1 | 2.50E-110 | 0.848568733 | 0.618 | 0.167 | 1.15E-105 | Upregulated |
| Edil3 | 4.44E-64 | 0.826525888 | 0.703 | 0.322 | 2.04E-59 | Upregulated |
| Itgb6 | 2.46E-44 | 0.820458446 | 0.886 | 0.704 | 1.13E-39 | Upregulated |
| Flna | 1.01E-53 | 0.819692051 | 0.641 | 0.322 | 4.63E-49 | Upregulated |
| Ywhaz | 2.57E-61 | 0.808945319 | 0.824 | 0.577 | 1.18E-56 | Upregulated |
| Ctgf | 6.14E-38 | 0.795971618 | 0.375 | 0.134 | 2.82E-33 | Upregulated |
| Grb10 | 3.99E-93 | 0.79178052 | 0.652 | 0.203 | 1.83E-88 | Upregulated |
| Rab8b | 2.51E-88 | 0.789260181 | 0.58 | 0.168 | 1.15E-83 | Upregulated |
| Edn1 | 7.24E-78 | 0.758274451 | 0.569 | 0.173 | 3.33E-73 | Upregulated |
| Spock2 | 9.42E-210 | 0.739556877 | 0.393 | 0.008 | 4.33E-205 | Upregulated |
| Plxnb2 | 2.29E-84 | 0.738329269 | 0.656 | 0.232 | 1.05E-79 | Upregulated |
| Picalm | 5.38E-51 | 0.705187305 | 0.848 | 0.546 | 2.47E-46 | Upregulated |
| Col6a1 | 2.72E-241 | 0.698502182 | 0.449 | 0.009 | 1.25E-236 | Upregulated |
| Clic5 | 3.53E-90 | 0.680726455 | 0.538 | 0.129 | 1.62E-85 | Upregulated |
| Nek6 | 3.69E-81 | 0.674395185 | 0.596 | 0.184 | 1.69E-76 | Upregulated |
| Bcl2l1 | 2.02E-51 | 0.657231296 | 0.868 | 0.545 | 9.27E-47 | Upregulated |
| Srgap1 | 7.52E-131 | 0.655755556 | 0.516 | 0.079 | 3.45E-126 | Upregulated |
| Cttnbp2nl | 1.74E-94 | 0.628249513 | 0.592 | 0.154 | 8.02E-90 | Upregulated |
| Plaur | 2.05E-40 | 0.621762758 | 0.65 | 0.349 | 9.43E-36 | Upregulated |
| Col18a1 | 8.66E-222 | 0.608265817 | 0.46 | 0.017 | 3.98E-217 | Upregulated |
| Ifngr2 | 3.53E-50 | 0.600698485 | 0.714 | 0.387 | 1.62E-45 | Upregulated |
| Col4a3 | 4.03E-57 | 0.597610495 | 0.491 | 0.16 | 1.85E-52 | Upregulated |
| Sp100 | 2.34E-70 | 0.587661452 | 0.647 | 0.235 | 1.07E-65 | Upregulated |
| Sparc | 1.53E-187 | 0.576590382 | 0.382 | 0.012 | 7.03E-183 | Upregulated |

| | | | | | | |
|---|---|---|---|---|---|---|
| Smarcc1 | 1.23E-50 | 0.549480339 | 0.819 | 0.489 | 5.63E-46 | Upregulated |
| Angptl4 | 1.12E-31 | 0.544326281 | 0.272 | 0.084 | 5.15E-27 | Upregulated |
| Rrm2b | 8.50E-100 | 0.506952256 | 0.498 | 0.1 | 3.90E-95 | Upregulated |
| Thsd4 | 1.17E-32 | 0.502711424 | 0.879 | 0.673 | 5.39E-28 | Upregulated |
| Sema7a | 3.45E-168 | 0.485245018 | 0.346 | 0.011 | 1.59E-163 | Upregulated |
| Gprc5a | 2.11E-34 | 0.467441878 | 0.951 | 0.87 | 9.69E-30 | Upregulated |
| Adam12 | 4.64E-93 | 0.452958962 | 0.283 | 0.027 | 2.13E-88 | Upregulated |
| Eif4g2 | 5.37E-20 | 0.446329545 | 0.703 | 0.542 | 2.47E-15 | Upregulated |
| Zyx | 8.85E-80 | 0.434222993 | 0.46 | 0.106 | 4.07E-75 | Upregulated |
| Trabd2b | 7.76E-95 | 0.392820285 | 0.212 | 0.009 | 3.56E-90 | Upregulated |
| **Bach1** | **3.35E-29** | **0.390877714** | **0.723** | **0.437** | **1.54E-24** | **Upregulated** |
| Sec14l1 | 1.00E-59 | 0.386832236 | 0.558 | 0.185 | 4.61E-55 | Upregulated |
| Ppic | 8.31E-36 | 0.386707578 | 0.491 | 0.217 | 3.82E-31 | Upregulated |
| Lcn2 | 4.71E-11 | 0.378885076 | 0.326 | 0.182 | 2.16E-06 | Upregulated |
| Gpd2 | 5.02E-39 | 0.376792405 | 0.547 | 0.227 | 2.31E-34 | Upregulated |
| Lpgat1 | 1.32E-56 | 0.37085342 | 0.404 | 0.109 | 6.08E-52 | Upregulated |
| Krt80 | 6.01E-149 | 0.367180362 | 0.308 | 0.01 | 2.76E-144 | Upregulated |
| Gclc | 1.14E-88 | 0.363080679 | 0.324 | 0.042 | 5.22E-84 | Upregulated |
| Ap1s1 | 1.91E-48 | 0.360106554 | 0.406 | 0.129 | 8.77E-44 | Upregulated |
| Hipk2 | 3.92E-36 | 0.360067638 | 0.627 | 0.295 | 1.80E-31 | Upregulated |
| Ube2j1 | 2.66E-42 | 0.354486409 | 0.513 | 0.206 | 1.22E-37 | Upregulated |
| Prr5l | 3.33E-102 | 0.349899804 | 0.241 | 0.012 | 1.53E-97 | Upregulated |
| Kazn | 2.69E-15 | 0.349100179 | 0.614 | 0.392 | 1.24E-10 | Upregulated |
| Psmd1 | 1.18E-34 | 0.346958961 | 0.536 | 0.237 | 5.44E-30 | Upregulated |
| Rbm38 | 2.24E-94 | 0.312169056 | 0.319 | 0.037 | 1.03E-89 | Upregulated |
| Cd276 | 1.23E-50 | 0.307138472 | 0.386 | 0.109 | 5.67E-46 | Upregulated |
| Nol4l | 5.48E-28 | 0.305218186 | 0.513 | 0.241 | 2.52E-23 | Upregulated |
| Rbm3 | 2.00E-20 | 0.302332319 | 0.629 | 0.377 | 9.21E-16 | Upregulated |
| Cux1 | 1.19E-23 | 0.295050432 | 0.958 | 0.864 | 5.49E-19 | Upregulated |
| TP53INP1 | 2.30E-67 | 0.288916248 | 0.335 | 0.062 | 1.06E-62 | Upregulated |
| Prim2 | 7.83E-47 | 0.288181305 | 0.453 | 0.146 | 3.60E-42 | Upregulated |
| Cdc34 | 2.65E-47 | 0.283017454 | 0.321 | 0.081 | 1.22E-42 | Upregulated |
| Lasp1 | 1.89E-35 | 0.282043501 | 0.417 | 0.153 | 8.69E-31 | Upregulated |
| Mpzl1 | 5.36E-31 | 0.270233387 | 0.594 | 0.292 | 2.46E-26 | Upregulated |
| Hmgn2p46 | 1.20E-62 | 0.270103322 | 0.243 | 0.034 | 5.51E-58 | Upregulated |
| Tgm2 | 9.26E-38 | 0.266602547 | 0.234 | 0.055 | 4.25E-33 | Upregulated |
| Atl3 | 3.13E-39 | 0.266429798 | 0.417 | 0.146 | 1.44E-34 | Upregulated |
| Gnas | 1.61E-14 | 0.260155557 | 0.864 | 0.771 | 7.40E-10 | Upregulated |
| Lipa | 1.29E-35 | 0.260022828 | 0.408 | 0.151 | 5.93E-31 | Upregulated |
| Ctsb | 1.02E-16 | 0.251894043 | 0.826 | 0.682 | 4.70E-12 | Upregulated |
| Mal2 | 4.57E-08 | 0.251441931 | 0.714 | 0.598 | 0.002102 | Upregulated |
| Dab2ip | 3.39E-37 | 0.250886312 | 0.357 | 0.116 | 1.56E-32 | Upregulated |
| Dcakd | 3.69E-38 | 0.247400852 | 0.29 | 0.079 | 1.69E-33 | Upregulated |
| Itgb3 | 1.54E-82 | 0.244790504 | 0.185 | 0.008 | 7.09E-78 | Upregulated |
| Ehd2 | 2.75E-26 | 0.227573776 | 0.395 | 0.167 | 1.26E-21 | Upregulated |
| Abhd2 | 4.64E-11 | 0.225025994 | 0.859 | 0.728 | 2.13E-06 | Upregulated |
| Golt1b | 1.13E-21 | 0.222777626 | 0.295 | 0.117 | 5.21E-17 | Upregulated |
| Epb41l2 | 1.83E-23 | 0.2193333 | 0.326 | 0.131 | 8.39E-19 | Upregulated |
| Arhgap1 | 2.25E-23 | 0.218249924 | 0.337 | 0.138 | 1.03E-18 | Upregulated |
| Capn6 | 1.82E-111 | 0.216317517 | 0.188 | 0 | 8.34E-107 | Upregulated |
| Pcyt1a | 2.70E-37 | 0.214575339 | 0.35 | 0.11 | 1.24E-32 | Upregulated |
| Dera | 7.06E-25 | 0.21223684 | 0.529 | 0.264 | 3.25E-20 | Upregulated |
| Mrps23 | 9.47E-19 | 0.211586337 | 0.4 | 0.198 | 4.35E-14 | Upregulated |
| Tmem165 | 9.15E-13 | 0.209937887 | 0.895 | 0.769 | 4.20E-08 | Upregulated |
| Mtch2 | 4.29E-28 | 0.206092794 | 0.48 | 0.214 | 1.97E-23 | Upregulated |
| Epha4 | 3.66E-10 | 0.204035392 | 0.578 | 0.407 | 1.68E-05 | Upregulated |

| | | | | | | |
|---|---|---|---|---|---|---|
| Rnf122 | 2.71E-33 | 0.202292062 | 0.123 | 0.016 | 1.24E-28 | Upregulated |
| Pphln1 | 4.11E-21 | 0.201183755 | 0.641 | 0.36 | 1.89E-16 | Upregulated |
| Pafah1b2 | 8.90E-16 | 0.201178118 | 0.598 | 0.375 | 4.09E-11 | Upregulated |
| Rab12 | 1.07E-24 | 0.199381817 | 0.353 | 0.141 | 4.92E-20 | Upregulated |
| Sdc3 | 4.18E-63 | 0.195561704 | 0.221 | 0.026 | 1.92E-58 | Upregulated |
| Ralb | 4.93E-34 | 0.194733395 | 0.286 | 0.083 | 2.26E-29 | Upregulated |
| **Nras** | **1.01E-30** | **0.192782181** | **0.288** | **0.091** | **4.64E-26** | **Upregulated** |
| Mlf2 | 4.43E-14 | 0.191677685 | 0.496 | 0.297 | 2.04E-09 | Upregulated |
| Dpy19l1 | 2.94E-14 | 0.191152233 | 0.683 | 0.446 | 1.35E-09 | Upregulated |
| Tpcn1 | 2.68E-28 | 0.190511075 | 0.388 | 0.153 | 1.23E-23 | Upregulated |
| Tmem30a | 4.63E-10 | 0.188074363 | 0.511 | 0.343 | 2.13E-05 | Upregulated |
| Myo1c | 6.88E-18 | 0.184215918 | 0.498 | 0.264 | 3.16E-13 | Upregulated |
| Casp3 | 1.37E-34 | 0.176929731 | 0.241 | 0.061 | 6.28E-30 | Upregulated |
| Opa3 | 2.00E-09 | 0.175021803 | 0.317 | 0.177 | 9.21E-05 | Upregulated |
| Gpat4 | 1.76E-31 | 0.174898319 | 0.355 | 0.124 | 8.09E-27 | Upregulated |
| Lamc1 | 4.43E-18 | 0.174590919 | 0.685 | 0.416 | 2.04E-13 | Upregulated |
| Slc25a24 | 8.38E-26 | 0.173751843 | 0.375 | 0.15 | 3.85E-21 | Upregulated |
| Dpp3 | 4.81E-22 | 0.173418106 | 0.246 | 0.087 | 2.21E-17 | Upregulated |
| Bzw1 | 2.14E-15 | 0.171580448 | 0.571 | 0.334 | 9.82E-11 | Upregulated |
| Fxn | 1.44E-26 | 0.169073167 | 0.306 | 0.11 | 6.63E-22 | Upregulated |
| Uhrf1 | 1.70E-26 | 0.167405867 | 0.17 | 0.04 | 7.81E-22 | Upregulated |
| Wipf1 | 2.60E-63 | 0.16252236 | 0.212 | 0.023 | 1.20E-58 | Upregulated |
| Rhoa | 9.49E-09 | 0.158562952 | 0.817 | 0.697 | 0.000436 | Upregulated |
| Eif4e2 | 2.39E-15 | 0.157368385 | 0.453 | 0.25 | 1.10E-10 | Upregulated |
| Dmd | 5.84E-09 | 0.153361863 | 0.554 | 0.371 | 0.0002682 | Upregulated |
| Lamb1 | 3.26E-32 | 0.152440874 | 0.217 | 0.052 | 1.50E-27 | Upregulated |
| Gpn1 | 8.00E-30 | 0.151785884 | 0.228 | 0.062 | 3.67E-25 | Upregulated |
| Marveld2 | 1.84E-31 | 0.151052688 | 0.292 | 0.09 | 8.47E-27 | Upregulated |
| Nudt4 | 4.06E-19 | 0.149114044 | 0.42 | 0.206 | 1.87E-14 | Upregulated |
| Ap1m1 | 3.28E-29 | 0.141055447 | 0.295 | 0.097 | 1.51E-24 | Upregulated |
| Ilk | 3.18E-34 | 0.140315384 | 0.147 | 0.023 | 1.46E-29 | Upregulated |
| Rap2a | 9.80E-34 | 0.138881975 | 0.203 | 0.045 | 4.50E-29 | Upregulated |
| Nhlrc3 | 5.04E-24 | 0.135884968 | 0.257 | 0.087 | 2.32E-19 | Upregulated |
| Vamp3 | 1.73E-14 | 0.130334696 | 0.33 | 0.165 | 7.93E-10 | Upregulated |
| Ints6l | 4.32E-13 | 0.123306201 | 0.138 | 0.048 | 1.98E-08 | Upregulated |
| Myh10 | 2.62E-20 | 0.122700631 | 0.275 | 0.106 | 1.20E-15 | Upregulated |
| Cbx5 | 4.96E-14 | 0.120452507 | 0.393 | 0.206 | 2.28E-09 | Upregulated |
| Mcam | 7.33E-61 | 0.119926747 | 0.114 | 0.002 | 3.37E-56 | Upregulated |
| Nrep | 3.92E-28 | 0.116212921 | 0.292 | 0.097 | 1.80E-23 | Upregulated |
| Acer3 | 1.01E-17 | 0.110358261 | 0.402 | 0.195 | 4.62E-13 | Upregulated |
| Vps25 | 1.01E-12 | 0.109743498 | 0.301 | 0.15 | 4.65E-08 | Upregulated |
| Ctss | 6.76E-07 | 0.107767616 | 0.571 | 0.417 | 0.0310824 | Upregulated |
| Scamp3 | 1.95E-18 | 0.107480988 | 0.317 | 0.137 | 8.95E-14 | Upregulated |
| Pign | 1.13E-25 | 0.095986381 | 0.395 | 0.163 | 5.18E-21 | Upregulated |
| Atxn7l3b | 1.30E-18 | 0.095577569 | 0.205 | 0.07 | 5.99E-14 | Upregulated |
| Galns | 9.66E-19 | 0.094881197 | 0.185 | 0.061 | 4.44E-14 | Upregulated |
| Psme3 | 2.36E-16 | 0.092665842 | 0.306 | 0.136 | 1.08E-11 | Upregulated |
| Thyn1 | 4.98E-11 | 0.0916515 | 0.246 | 0.122 | 2.29E-06 | Upregulated |
| Syk | 4.09E-12 | 0.08457082 | 0.268 | 0.131 | 1.88E-07 | Upregulated |
| Il17re | 1.71E-16 | 0.082388566 | 0.183 | 0.063 | 7.85E-12 | Upregulated |
| Cntrl | 6.22E-21 | 0.081555589 | 0.27 | 0.101 | 2.86E-16 | Upregulated |
| Lrrc59 | 3.97E-09 | 0.081539784 | 0.263 | 0.142 | 0.0001824 | Upregulated |
| Rac2 | 1.45E-26 | 0.081261444 | 0.107 | 0.015 | 6.67E-22 | Upregulated |
| Sesn3 | 4.88E-18 | 0.079256504 | 0.138 | 0.038 | 2.24E-13 | Upregulated |
| Ints2 | 3.03E-13 | 0.078104645 | 0.17 | 0.065 | 1.39E-08 | Upregulated |
| Lrrc20 | 5.66E-13 | 0.077789937 | 0.152 | 0.056 | 2.60E-08 | Upregulated |

| | | | | | | |
|---|---|---|---|---|---|---|
| Pctp | 1.20E-18 | 0.076031589 | 0.109 | 0.024 | 5.52E-14 | Upregulated |
| Rap1b | 1.02E-06 | 0.073989036 | 0.741 | 0.571 | 0.0470249 | Upregulated |
| Ipo9 | 1.44E-12 | 0.072934945 | 0.404 | 0.226 | 6.60E-08 | Upregulated |
| Kctd10 | 3.06E-19 | 0.071681565 | 0.23 | 0.082 | 1.41E-14 | Upregulated |
| Mical1 | 1.03E-16 | 0.071429617 | 0.23 | 0.089 | 4.74E-12 | Upregulated |
| Adamtsl3 | 6.86E-09 | 0.071367337 | 0.114 | 0.045 | 0.0003153 | Upregulated |
| Wdr46 | 6.70E-12 | 0.070724413 | 0.188 | 0.079 | 3.08E-07 | Upregulated |
| Slc35b4 | 1.20E-13 | 0.066445615 | 0.1 | 0.027 | 5.51E-09 | Upregulated |
| Slc26a2 | 3.05E-19 | 0.061557771 | 0.118 | 0.027 | 1.40E-14 | Upregulated |
| Hif1an | 4.10E-13 | 0.060439983 | 0.156 | 0.057 | 1.88E-08 | Upregulated |
| Hif1an.1 | 4.10E-13 | 0.060439983 | 0.156 | 0.057 | 1.88E-08 | Upregulated |
| Fam98a | 3.50E-08 | 0.057550898 | 0.125 | 0.055 | 0.0016096 | Upregulated |
| Nectin3 | 4.13E-07 | 0.057254319 | 0.268 | 0.156 | 0.0189747 | Upregulated |
| Lsp1 | 3.99E-14 | 0.053738715 | 0.121 | 0.036 | 1.83E-09 | Upregulated |
| Elf4 | 1.03E-17 | 0.052806605 | 0.176 | 0.057 | 4.71E-13 | Upregulated |
| Tmem50b | 2.42E-07 | 0.051664378 | 0.357 | 0.222 | 0.0111369 | Upregulated |
| Snx18 | 8.97E-14 | 0.051003529 | 0.154 | 0.054 | 4.12E-09 | Upregulated |
| Ptafr | 8.68E-15 | 0.050331376 | 0.109 | 0.029 | 3.99E-10 | Upregulated |
| Rrm1 | 7.10E-07 | 0.049943054 | 0.174 | 0.091 | 0.0326086 | Upregulated |
| Traf4 | 2.22E-17 | 0.049233535 | 0.109 | 0.026 | 1.02E-12 | Upregulated |
| Tgfbrap1 | 1.84E-12 | 0.045807766 | 0.156 | 0.059 | 8.43E-08 | Upregulated |
| Arl2bp | 3.29E-10 | 0.04461511 | 0.125 | 0.047 | 1.51E-05 | Upregulated |
| Itpripl2 | 1.64E-09 | 0.04053469 | 0.152 | 0.065 | 7.53E-05 | Upregulated |
| Fgr | 7.87E-09 | 0.038723523 | 0.1 | 0.037 | 0.0003615 | Upregulated |
| Stx17 | 2.62E-11 | 0.03754101 | 0.266 | 0.131 | 1.20E-06 | Upregulated |
| Eif1ad | 1.84E-09 | 0.036285548 | 0.208 | 0.101 | 8.47E-05 | Upregulated |
| Emilin2 | 1.54E-07 | 0.030475835 | 0.234 | 0.129 | 0.007095 | Upregulated |
| **Ezh2** | **3.51E-07** | **0.027859595** | **0.199** | **0.107** | **0.0161381** | **Upregulated** |
| **Arid3a** | **3.48E-08** | **0.025063022** | **0.252** | **0.136** | **0.001597** | **Upregulated** |
| Clip1 | 2.52E-11 | 0.015944528 | 0.801 | 0.586 | 1.16E-06 | Upregulated |
| Nipa2 | 1.47E-08 | 0.015724896 | 0.243 | 0.128 | 0.0006771 | Upregulated |
| Aven | 2.55E-11 | 0.014389443 | 0.288 | 0.146 | 1.17E-06 | Upregulated |
| Pqlc2 | 1.05E-07 | 2.18E-05 | 0.172 | 0.084 | 0.0048334 | Upregulated |
| Stk4 | 1.75E-09 | -0.001728265 | 0.279 | 0.147 | 8.05E-05 | Downregulated |
| Ncbp1 | 1.01E-07 | -0.012884492 | 0.232 | 0.124 | 0.0046621 | Downregulated |
| Clic4 | 1.87E-08 | -0.027142781 | 0.739 | 0.56 | 0.0008612 | Downregulated |
| Rbpms-as1 | 3.28E-12 | -0.195244053 | 0.033 | 0.154 | 1.51E-07 | Downregulated |
| Clu | 9.33E-10 | -0.219686589 | 0.078 | 0.189 | 4.29E-05 | Downregulated |
| Slc26a9 | 6.33E-18 | -0.381004257 | 0.04 | 0.209 | 2.91E-13 | Downregulated |
| Cd38 | 1.43E-12 | -0.438329319 | 0.223 | 0.368 | 6.57E-08 | Downregulated |
| Ttc28 | 1.63E-07 | -0.489972622 | 0.645 | 0.678 | 0.007495 | Downregulated |
| Vapa | 1.74E-26 | -0.576695729 | 0.754 | 0.789 | 8.02E-22 | Downregulated |
| Osmr | 1.22E-10 | -0.618245034 | 0.65 | 0.675 | 5.60E-06 | Downregulated |
| Anxa1 | 4.67E-12 | -0.645902657 | 0.842 | 0.807 | 2.14E-07 | Downregulated |
| **Myc** | **4.72E-33** | **-0.663377486** | **0.096** | **0.365** | **2.17E-28** | **Downregulated** |
| Gata6 | 6.06E-12 | -0.688601345 | 0.438 | 0.524 | 2.78E-07 | Downregulated |
| Acsl1 | 9.73E-24 | -0.701821485 | 0.464 | 0.615 | 4.47E-19 | Downregulated |
| Tcf7l1 | 1.17E-08 | -0.704158117 | 0.634 | 0.63 | 0.0005359 | Downregulated |
| **Foxp2** | **4.11E-24** | **-0.847670257** | **0.408** | **0.591** | **1.89E-19** | **Downregulated** |
| Npnt | 6.16E-33 | -0.907264519 | 0.281 | 0.534 | 2.83E-28 | Downregulated |
| Slc6a14 | 1.02E-35 | -1.04133692 | 0.283 | 0.554 | 4.70E-31 | Downregulated |
| Acsl4 | 3.61E-24 | -1.056821681 | 0.56 | 0.643 | 1.66E-19 | Downregulated |
| Ctsh | 1.27E-61 | -1.315580824 | 0.634 | 0.784 | 5.85E-57 | Downregulated |
| Sod2 | 4.81E-136 | -2.079217632 | 0.634 | 0.893 | 2.21E-131 | Downregulated |
