## Supplementary Data 10 for "*Let-7* restrains an oncogenic circuit in AT2 cells to prevent fibrogenic cell intermediates in pulmonary fibrosis"

| GSEA Gene Sets | Source or Reference |
| --- | --- |
| Proliferation Epcam+ cells | https://hschillerlabshiny.shinyapps.io/Bleo_webtool/ |
| ADI Cells | https://hschillerlabshiny.shinyapps.io/Bleo_webtool/ |
| Homeostatic AT2 cells | https://hschillerlabshiny.shinyapps.io/Bleo_webtool/ |
| AT1 Cells | https://hschillerlabshiny.shinyapps.io/Bleo_webtool/ |
| Basal Cells | https://hschillerlabshiny.shinyapps.io/Bleo_webtool/ |
| Club Cells | https://hschillerlabshiny.shinyapps.io/Bleo_webtool/ |
| Cluster7 ADI cells | Wang et al., 2023 (PMID: 37768734) |
| Senescence-associated secretory phenotype | Wang et al., 2023 (PMID: 37768734) |
| Senescence | Wang et al., 2023 (PMID: 37768734) |
| DNA damage | Wang et al., 2023 (PMID: 37768734) |
| Cell death | Wang et al., 2023 (PMID: 37768734) |
